# Multi-omic identification of key transcriptional regulatory programs during endurance exercise training in rats

**DOI:** 10.1101/2023.01.10.523450

**Authors:** Gregory R. Smith, Bingqing Zhao, Malene E. Lindholm, Archana Raja, Mark Viggars, Hanna Pincas, Nicole R. Gay, Yifei Sun, Yongchao Ge, Venugopalan D. Nair, James A. Sanford, Mary Anne S. Amper, Mital Vasoya, Kevin S. Smith, Stephen Montgomery, Elena Zaslavsky, Sue C. Bodine, Karyn A. Esser, Martin J. Walsh, Michael P. Snyder, Stuart C. Sealfon, the MoTrPAC Study Group

**Author notes:** These authors contributed equally (Gregory R. Smith, Bingqing Zhao).

## Abstract

Transcription factors (TFs) play a key role in regulating gene expression. We conducted an integrated analysis of chromatin accessibility, DNA methylation, mRNA expression, protein abundance and phosphorylation across eight tissues in fifty rats of equally represented sexes following endurance exercise training (EET) to identify coordinated epigenomic and transcriptional changes and determine key TFs involved. We uncovered tissue-specific EET associated changes and TF motif enrichment across differentially expressed genes (DEGs), accessible regions (DARs), and methylated regions (DMRs). We discovered distinct routes of EET-induced regulation through either epigenomic alterations providing better access for TFs to affect target genes, or via changes in TF expression or activity enabling target gene responses. We identified TF motifs enriched among correlated epigenomic and transcriptomic alterations, DEGs correlated with exercise-related phenotypic and cell type composition changes, and EET-induced activity changes of TFs whose target genes are enriched for DEGs. This analysis elucidates the unique gene regulatory mechanisms mediating diverse transcriptional responses to EET across tissues.

## Introduction

Regular exercise impacts health and mitigates disease processes throughout the body. Exercise maintains muscle function, improves cardiovascular wellness and cognitive performance, and lowers the risk of cardiovascular disease and many other disorders, including cancers and dementia^1^. The molecular processes mediating the adaptations induced by exercise training across tissues are poorly understood.

As the main regulators of gene transcription, transcription factors (TFs) act via the recruitment of other factors, co-activators, or co-repressors to cis-regulatory elements at the promoter or distal regions of target genes. The access of TFs to cis-motifs partly depends on chromatin organization. Hence, along with changes in chromatin accessibility and other epigenetic modifications, including DNA methylation, TFs govern constitutive gene expression as well as gene responses to stimuli in tissues. TFs are critical exercise-response mediators^2,3^ and, in skeletal muscle, exercise training-induced transcriptomic changes have been associated with different TFs than those induced by acute exercise^4^.

Our companion multi-tissue analysis of the molecular response dynamics during endurance exercise training found that the majority of differentially regulated genes are tissue-specific, whereas a small proportion are shared across multiple tissues^5,6^. Thus, gene responses to training are likely mediated through the combinatorial action of tissue-enriched and shared transcriptional regulators. Shared exercise-induced TF regulation can elicit tissue-specific functions, as seen with PPARγ, which is implicated in PGC1α-stimulated mitochondrial biogenesis in muscle^7^, regulation of adipogenesis^8^, and hippocampal BDNF activity and its cognitive effects^9^. As complex regulatory patterns drive tissue-specific gene regulation^10^, they likely are involved in mediating the diverse effects of exercise training on tissues. Therefore, in order to unravel the complex regulatory processes driving EET response, we need epigenomic as well as transcriptomic data covering diverse tissues that can all be affected by exercise.

However, few studies have evaluated training-induced genome-wide changes in RNA expression, chromatin accessibility, and DNA methylation^11^, and the majority have concentrated on skeletal muscle.

We leveraged the study design of the Molecular Transducers of Physical Activity Consortium (MoTrPAC) endurance exercise training (EET) study in rats^5^ to characterize the TFs mediating gene responses to training across multiple tissues. During 8 weeks of EET, genome-wide transcriptome, chromatin accessibility, and DNA methylation were assayed in 8 tissues from age-matched male and female rats: gastrocnemius (SKM-GN), heart, hippocampus (HIPPOC), kidney, liver, lung, brown adipose (BAT), and white adipose (WAT-SC). Protein abundance and phosphorylation data was assayed in six of these tissues (SKM-GN, heart, kidney, liver, lung, and WAT-SC) and we identified relationships between EET responses in TF activity at the proteomic level with changes in TF target EET response. Through integrative analysis of all omes for each tissue, we establish a map of the regulatory transcriptional responses to training across tissues.

## Results

### Characterization of epigenetic and transcriptional responses to endurance training

To understand the epigenetic and transcriptional response mechanisms elicited during eight weeks of EET, we analyzed ATAC-seq, RNA-seq and RRBS profiles generated in skeletal muscle (gastrocnemius; SKM-GN), heart, hippocampus (HIPPOC), kidney, liver, lung, brown adipose tissue (BAT), and subcutaneous white adipose tissue (WAT-SC) from fifty rats divided into groups of 10 animals (5 males, 5 females). Animals were subjected to either 1, 2, 4, or 8 weeks of training (Fig 1a) or to no training (untrained controls), consisting of 8 weeks of housing and stationary treadmills. Protein abundance and protein phosphorylation were also measured in six of the eight tissues: SKM-GN, heart, kidney, liver, lung, and WAT-SC. We identified differentially accessible regions (DARs; F test adjusted p-value < 0.1), differentially methylated regions (DMRs: F test adjusted p-value < 0.1), and differentially expressed genes (DEGs; F test adjusted p-value < 0.1) between EET and control groups (Fig 1b). To characterize the transcriptional and epigenomic changes induced by EET across tissues, we evaluated the tissue-specificity of DARs, DMRs, and DEGs. Although most expressed genes, open chromatin sites, and methylated CpG sites were detectable in multiple tissues, the majority of DARs (90%), DMRs (91%), and DEGs (66%) were identified in only one tissue (Fig 1c, Fig. S1). This suggested that gene regulatory responses to EET were largely specific to individual tissues, which was in line with another MoTrPAC publication^6^.

**Figure 1:**
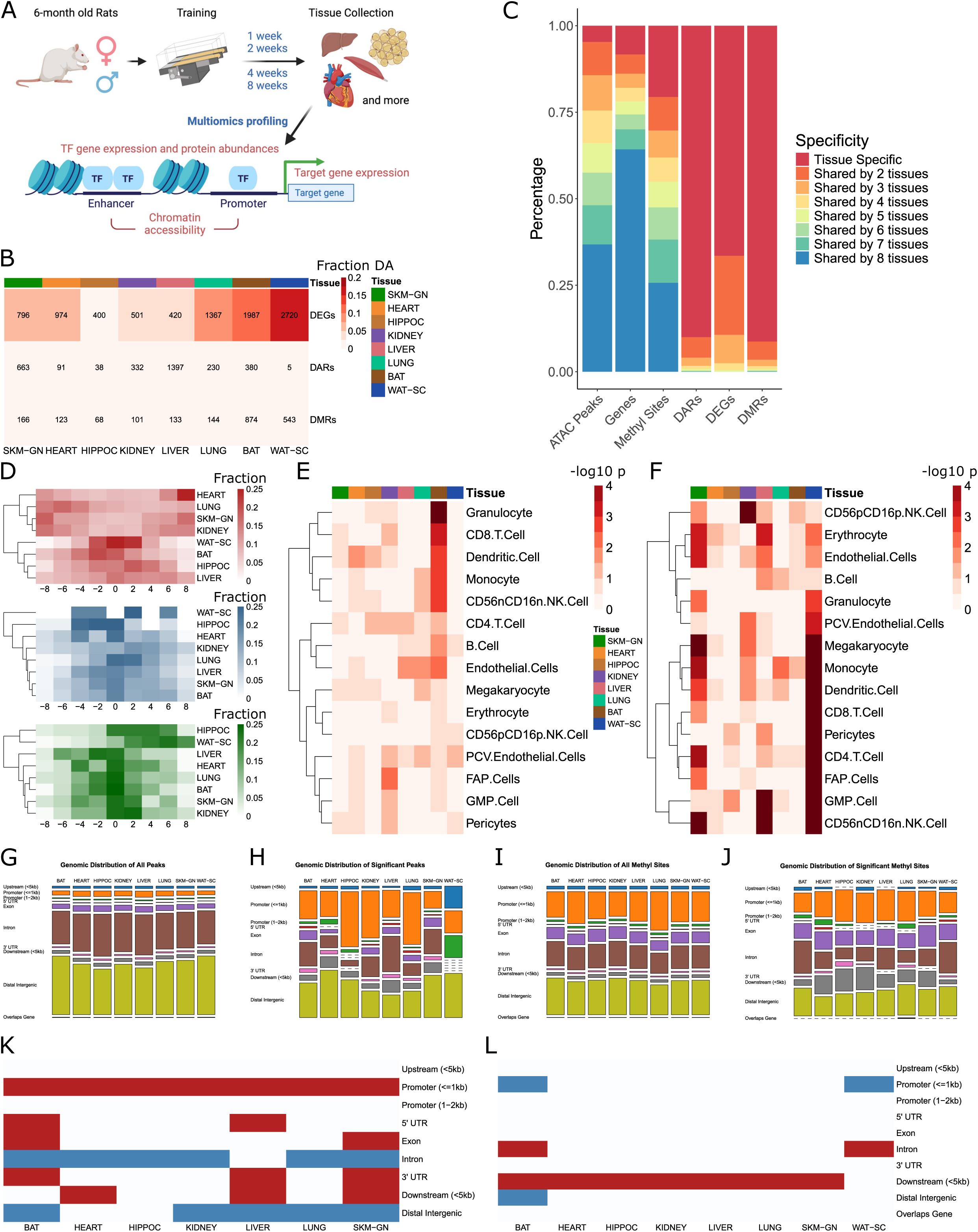
Epigenetic and transcriptional responses to training programs. (a) 6-month old rats of both sexes underwent endurance exercise training (EET) programs. Tissues were collected and subjected to multiomics profiling, including ATAC-seq, RNA-seq, proteomics. (b) A higher number and percentage of differentially expressed genes (DEGs) were identified than differentially accessible regions (DARs) and differentially methylated regions (DMRs) after training in most tissues (F test FDR<0.1). (c) Many accessible regions, methylation sites, and genes were identified in all tissues while training-induced features were highly tissue-specific. (d) Distribution of L2FC positive/negative consistency in RNA-seq (in red), ATAC-seq (in blue) and RRBS differential analytes (in green) across tissues. The sum of the sign(L2FC) at each time point in males and females for each analyte is calculated with values ranging from -8 (negative L2FC at all time points and sexes) to 0 (half positive and half negative L2FC) to 8 (positive L2FC at all time points and sexes). Heart and SKM-GN RNA-seq DEGs are more consistently up or down-regulated while WAT-SC RNA-seq DEGs are less consistent, for example. (e,f) Cell type deconvolution analysis-generated -log10 p-values of Kruskal-Wallis test measuring significant predicted changes in tissue cell type composition based on training (e) or sex (f). Brown adipose exhibited increased proportions of immune cell types after training. White adipose exhibited sex-specific changes in proportions of immune cell types and pericytes. (g-j) Distribution of genomic locations of all accessible regions (g), DARs (h), all methylation sites (i) and DMRs (j). (k) DARs enriched for the proximal promoter compared to all accessible regions. (l) DMRs are enriched for the downstream region compared to all methylation sites.

We then examined the distribution patterns of log2 fold change (L2FC) in gene expression, L2FC in chromatin accessibility, and L2FC in methylation across time points and sexes (Fig. S2, S3, S4 respectively). While the ratios of up-to down-regulated analytes (i.e. DEGs, DARs, DMRs) were similar across the majority of tissues, L2FC patterns differed between tissues and omes. Overall, methylated regions showed the largest spread of L2FC values compared to expressed genes and chromatin accessibility peaks, a pattern that was even more pronounced when comparing DMRs to DEGs and DARs. DEGs and DMRs were more likely to have larger L2FC values than non-significant features in their respective omes; however, DARs did not share this trend. Multiomic power analysis of the five omes included in this study identified RNA-seq as the ome with a consistently higher power and the phosphoproteome with consistently the least power (Fig. S5).

**Figure 2:**
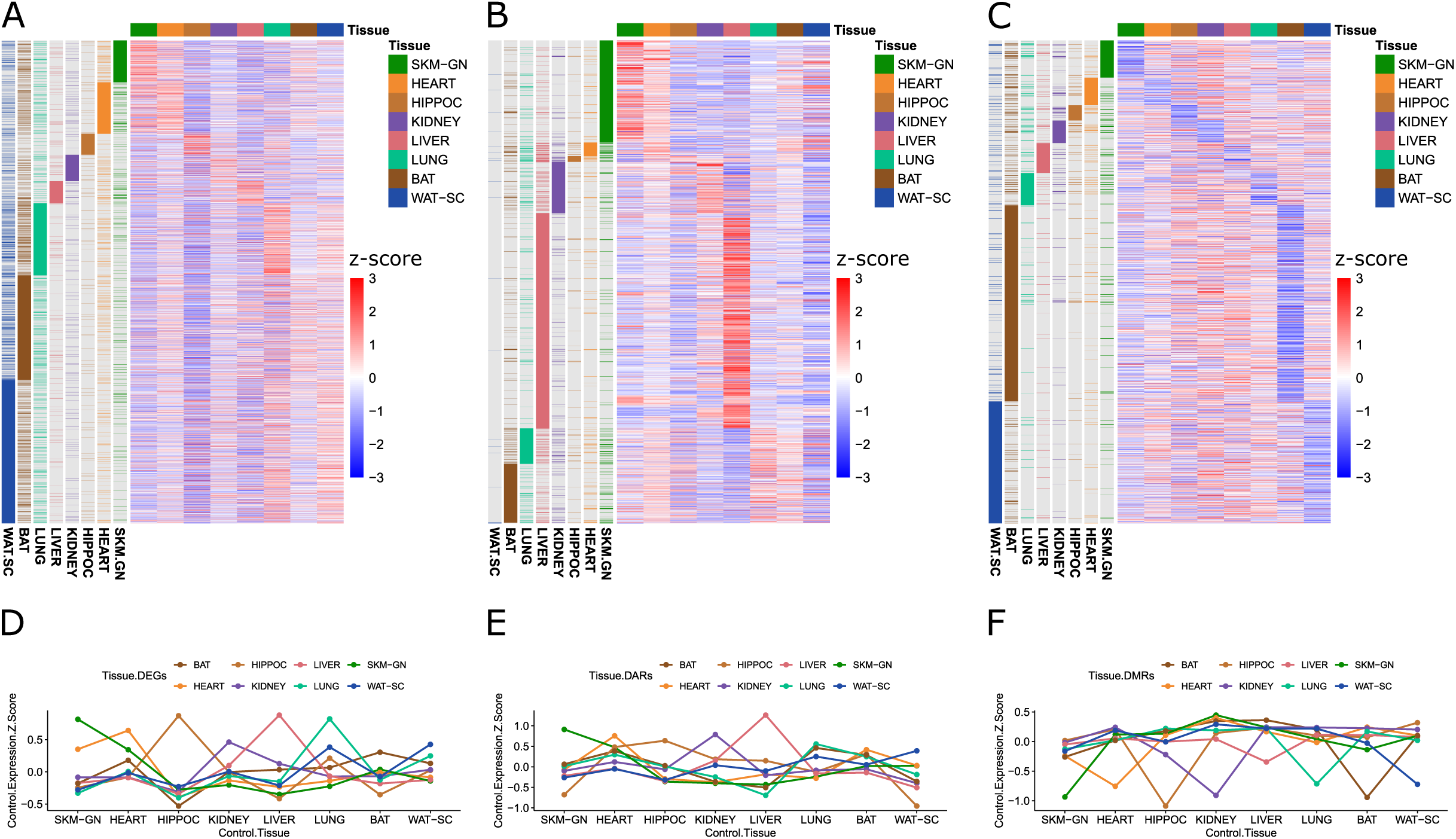
In general, genes are more highly expressed, chromatin peaks are more highly accessible and methylation sites are more hypomethylated in the tissues in which they exhibit a differential training response. (a-c) heatmaps of all differential analytes across eight tissues in RNA-seq (a), ATAC-seq (b), and RRBS (c). Columns reflect z scores of baseline expression (a), accessibility (b), or methylation (c), across the eight tissues. Rows are annotated by the tissues in which each analyte exhibits a differential training response. (d-f) Mean z-score of control gene expression (d), chromatin peak accessibility (e), and site methylation (f), for the differential training response analytes for each tissue.

**Figure 3:**
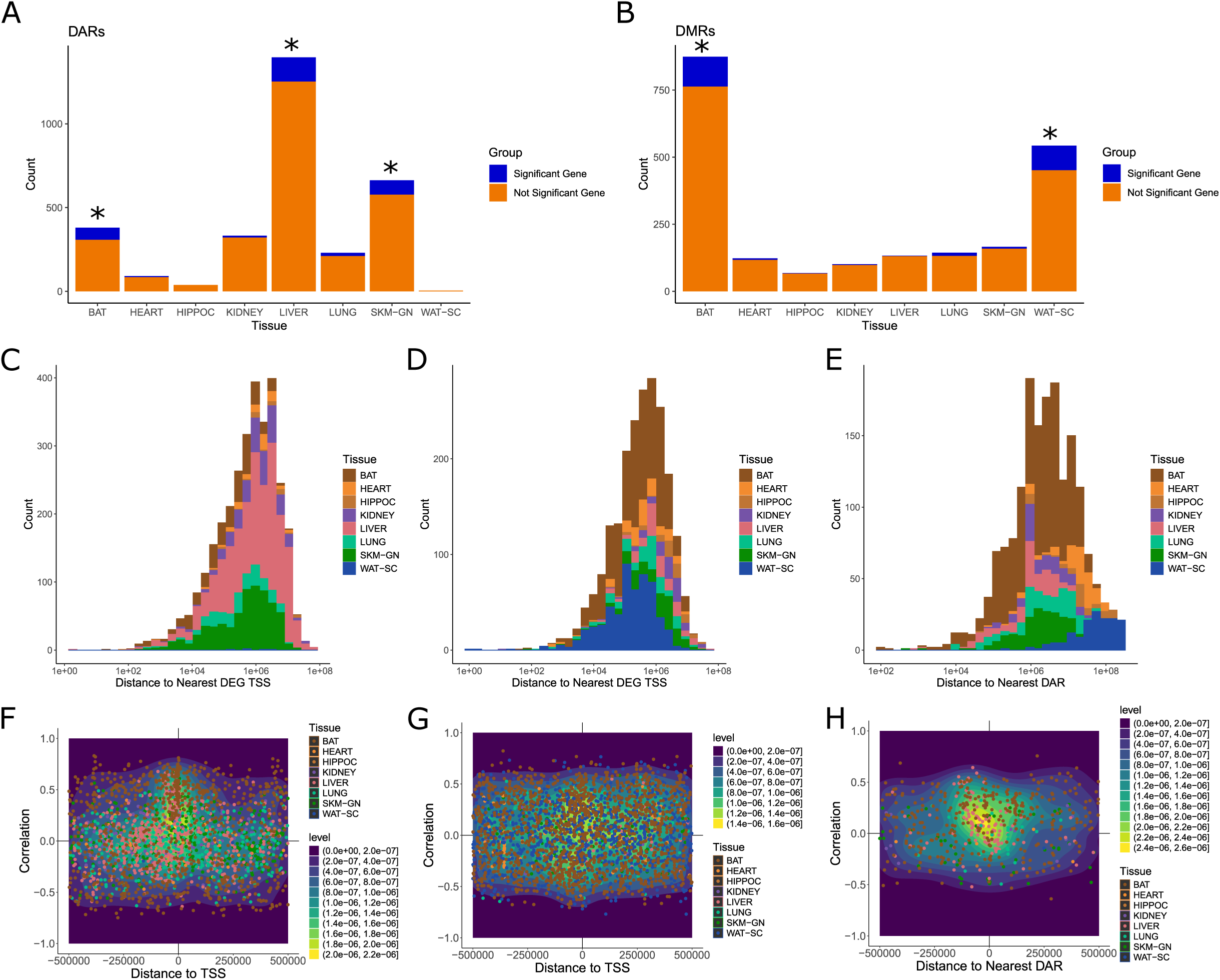
Responses in chromatin accessibility and DNA methylation may not directly link with the expression of the closest genes. (a) Count of DARs annotated to a DEG or a non-DEG. The closest gene to most DARs is not a DEG. * reflects p < 0.05 for hypergeometric test measuring the significance of the DAR-DEG overlap. (b) Count of DMRs annotated to a DEG or a non-DEG. Similar to DARs, the closest gene to most DMRs is not a DEG. * reflects p < 0.05 for hypergeometric test measuring the significance of the DMR-DEG overlap (c-e) Distributions of distance between DARs and nearest DEG TSS (c), DMRs and nearest DEG TSS (d), and DMRs and nearest DAR (e). DARs (c) and DMRs (d,e) are colored by tissue. (f-h) Density scatter plots of DAR-DEG training response correlation vs. distance (f), DMR-DEG training response correlation vs distance (g), and DMR-DAR training response correlation vs distance (h). DARs with high positive correlation to gene expression enriched for TSS-proximal regions in most tissues while DMRs with high positive and negative correlation to gene expression enriched for TSS-proximal regions. DAR-DMR correlations tended more positive when DMRs were upstream of DARs, and tended more negative when DMRs were downstream of DARs.

**Figure 4:**
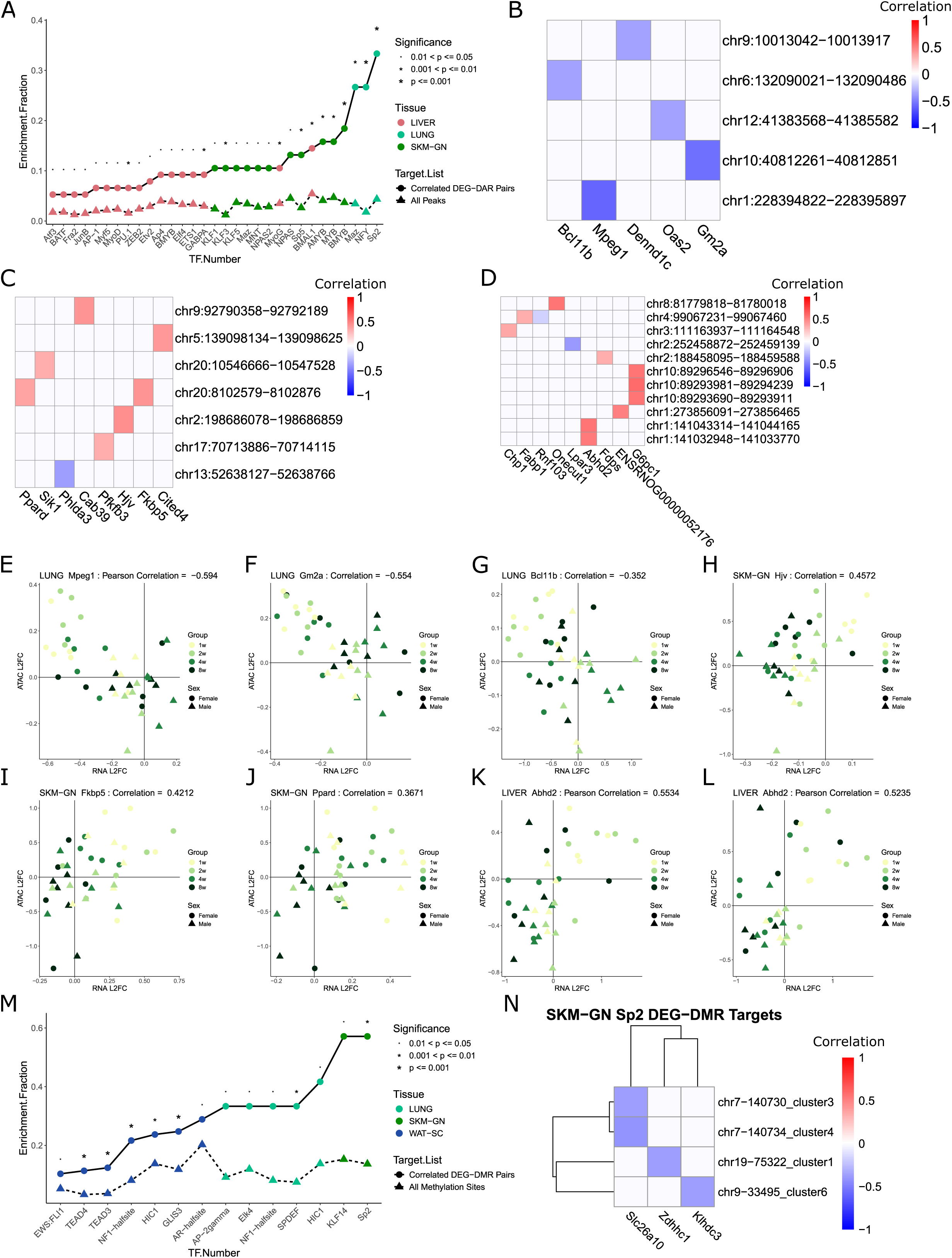
Responsive transcription factors in correlated DAR-DEG pairs. (a) TFs enriched for DAR-DEG pairs in a given tissue. * represents p < 0.05 in binomial test of frequency of TF motif presence in DAR-DEG pairs vs frequency in active peaks in tissue. (b-d) DAR-DEG pairs containing the binding motif of SP2 in lung (b), BMYB in SKM-GN (c), and BMAL1 in liver (d). (e-l) Individual DAR-DEG pair targets of enriched TFs and their cross-ome EET response correlation, including *Mpeg1* (e), *Gm2a* (f), *Bcl11b* (g), *Hjv* (h), *Fkbp5* (i), *Ppard* (j), and two separate DAR-DEG pairs involving *Abhd2* (k-l). (m) TFs enriched for DMR-DEG pairs in a given tissue. * represents p < 0.05 in binomial test of frequency of TF motif presence in DMR-DEG pairs vs frequency in methylation regions in tissue. (n) DMR-DEG pairs containing the binding motif of SP2 in SKM-GN.

Sex differences were the main driver for sample segregation in most tissues along top PC axes in RNA-seq (Fig. S6), ATAC-seq (Fig. S7), and RRBS (Fig. S8). We observed sex differences among DEGs in HIPPOC, BAT, and WAT-SC. Sex differences were also prevalent among DARs and DMRs in a majority of tissues. Overall, the proportion of DEGs showing concordant changes (L2FC) across time points and sexes was higher than that of DARs and DMRs (Fig. 1d). Notably, heart, SKM-GN, and kidney exhibited the most consistency in expression changes across all groups, whereas WAT-SC showed the least. In BAT, we detected pronounced variations in DEG as well as in DAR profiles between earlier and late time points in both sexes (Fig. S2g and S3g). Gene set enrichment analysis among week 1 or week 8 DEGs in males and females highlighted varying pathway enrichment patterns that were, in most cases, consistent across sexes and time points (Fig S9). SKM-GN and heart shared enrichment for oxidative phosphorylation and cardiac muscle contraction pathways, as well as markers for Parkinson’s, Huntington’s, and Alzheimer’s diseases. WAT-SC DEGs were enriched for the chemokine signaling pathway and immune-related diseases, including systemic lupus erythematosus, asthma, and primary immunodeficiency.

Next, we investigated whether alterations in cell type proportions contributed to the expression changes observed. Cell type deconvolution analysis (See Methods) identified large-scale changes in immune cell type proportions that were attributable to training duration in BAT (Fig. 1e and Fig. S10a) and to sex independent of training in WAT (Fig. 1f and Fig. S10b). Changes in dendritic cell proportion in heart, fibro/adipogenic progenitor (FAP) cell proportion in kidney, and monocyte and endothelial cell proportions in lung were also notable. Significant percentages of DEGs in many tissues were either correlated or anti-correlated with changes in immune cell type composition in response to EET (Fig. S11), with the most consistent correlations occurring in BAT or WAT-SC. Large percentages of DEGs were also correlated with changes in natural killer (NK) and granulocyte/monocyte progenitor (GMP) cells in liver and monocytes in lung. A full list of correlations between DEGs and cell type compositions can be found in Supplemental Table S1. Because the immune cell type composition differences observed between female and male WAT-SC were independent of exercise training, and because differential analysis was conducted in male and female samples separately, the DEGs identified in WAT-SC in each sex do not reflect any cell type composition changes occurring during training.

To further characterize the epigenomic changes induced by EET across tissues, we examined the genomic distribution of DARs vs. that of all open chromatin regions detected (Fig. 1g,h).

Compared to all accessible regions, DARs were significantly enriched at proximal promoters across all tissues except WAT, which was excluded due to a scarcity of DARs (Fig 1h,k, Methods). Consistent with previous studies^12,13^, open chromatin peaks across tissues were predominantly located in intronic and distal intergenic regions. Given the importance of the proximal promoter in the regulation of gene transcription^14^, the enrichment of DARs in this region suggested that EET results in the transcriptional activation of target genes. DMRs were significantly enriched at downstream regions across all tissues except WAT, relative to all methylation sites in each tissue (Fig 1i,j,l). Unlike DARs and DMRs, open chromatin regions that mapped to DEGs, which we refer to as DEG-associated peaks (DEGaPs), shared a similar genomic distribution as the peaks associated with all expressed genes (Fig S12); however, ATAC-seq peaks that mapped to either expressed genes or DEGs contained a higher proportion of intronic peaks and a lower proportion of distal intergenic peaks compared to all open chromatin peaks in the dataset. This was in part explained by a subset of unexpressed genes only containing distal intergenic peaks, suggesting that some gene-body chromatin accessibility was important for gene expression.

Differential analytes in a tissue also tend to have higher baseline enrichment in that tissue (Figure 2). Upon calculating the z scores of baseline expression of DEGs, accessibility of DARs, and methylation of DMRs across all 8 tissues, we found, in the case of both RNA-seq and ATAC-seq data, DEGs and DARs in a given tissue were, on average, more highly expressed (Figure 2a,d) and more highly accessible (Figure 2b,e), respectively, at baseline in the tissue with the significant training response. Conversely, DMRs in a given tissue were more hypomethylated at baseline in the tissue with a significant training response (Figure 2c,f). This pattern of tissue specificity was mostly maintained when subdividing differential analytes into up- and down-regulated subsets (Fig S13); however, down-regulated DMRs were less likely to follow this pattern, most notably seen with BAT. While there is some overlap in significant training responses across tissues, the majority of training response analytes represent tissue-enriched analytes, even if they are not tissue-specific.

### Identification of distal correlated epigenetic regulation despite few DARs and DMRs mapping to adjacent DEGs

We next sought out DAR-DEG associations by assessing the concordance between chromatin accessibility and gene expression changes. We assigned each DAR to the nearest gene and determined the fraction of DARs that were annotated to DEGs. Applying a hypergeometric test, we found that BAT, SKM-GN, and liver showed a higher proportion of overlap between DARs and DEGs (Fig 3a). The substantial overlap between DARs and DEGs in BAT may be related to the EET-induced increase in immune cell populations (see Fig. 1e). SKM-GN and liver showed the highest count of DARs among all tissues (Fig. 1b). Despite hundreds of DARs in both kidney and lung, only a few of their nearest genes were DEGs (Fig 3a). Similarly, we investigated DMR-DEG associations by assigning each DMR to the nearest gene and determining the fraction that was annotated to DEGs. Only BAT and WAT-SC have significant overlaps between DMRs and DEGs (Fig 3b). BAT and WAT-SC are the two tissues with the largest number of DMRs in the dataset by a considerable margin (Fig 1b).

The binding of TFs to distal open chromatin regions can regulate gene transcription^15,16^. Given the modest proportion of DARs mapped to adjacent DEGs, we extended the search window and sought relationships outside the closest gene for a given DAR. With respect to the location of DARs relative to the TSS of the nearest DEG, in all tissues, the majority of nearest DAR-DEG pairs reflected a normal distribution with a median centered approximately 1 Mb away from the nearest DEG, and a substantial left tail representing closer pairs (Fig 3c). BAT, SKM-GN, and liver contributed most of the DARs proximal to a DEG, confirming our earlier observations (Fig 3a). A similar pattern is seen when measuring the distance between a DMR and the nearest DEG (Fig 3d) and the distance between a DMR and the nearest DAR (Fig 3e). BAT and WAT-SC contain the majority of closest DMR-DEG pairs likely because of their greater DMR-DEG overlap (Fig 3b) and BAT contains the majority of closest DMR-DAR relationships. WAT-SC DMR-DAR relationships tend to be the most distant, largely because of the lack of DARs, reducing the likelihood of neighboring interactions.

DARs that were adjacent to DEGs tended to be more highly correlated with gene expression changes (Fig 3f, Fig S14), a pattern predominantly seen in SKM-GN and liver. In contrast, we found less of a connection between distance to TSS and correlation in training response for DMRs adjacent to DEGs, although there were still a large number of highly correlated adjacent pairs, especially seen in BAT and WAT-(Fig 3g, Fig S15). Interestingly, while instances of adjacent DMRs and DARs are more limited in this study, in tissues more populated by DAR-DMR pairs, positive correlation tended to occur when the DMR was upstream of the DAR, and correlation was decreased or negative when the DMR was downstream of the DAR (Fig 3h, Fig S16), most evidently seen in liver.

### DAR-DEG pairs are associated with distinct pathways in each tissue and SP2, BMYB, and BMAL1 represent key regulatory TFs

We focused on DARs and DEGs located within 500 kb from each other and isolated those that were either positively or negatively correlated across samples at all time points and sexes (t test of Pearson correlation p-value < 0.05; Supplementary Table S2). Pathway enrichment analysis among DAR-DEG pairs identified largely tissue-specific enrichment patterns for each tissue (Fig S17). In agreement with the training-associated increase in immune cell types inferred from cell type deconvolution analysis (see Fig. 1e), DAR-DEG pairs in BAT showed enrichment for several immune pathways. Lung also showed considerable enrichment for immune-associated pathways, suggesting activating roles in antigen defense by exercise. By contrast, liver DAR-DEG pairs were primarily enriched for lipid biosynthesis and metabolic processes, while heart DAR-DEG pairs were enriched for muscle movement and filament sliding, and SKM-GN DAR-DEG pairs were enriched for myofiber synthesis and muscle contraction (Fig S17). These results suggested that the correlated epigenetic and transcriptional changes induced by training affected tissue-specific functions.

To identify key regulators of the training response in each tissue, we analyzed TF motif enrichment at the DARs of DAR-DEG pairs (Fig. 4a). Notably, we identified SP2 in lung as the most significantly enriched TF with 33.33% of DAR-DEG pairs in lung containing an SP2 binding site vs. a general enrichment of 4.35% (p-value = 0.00032). The DAR-DEG pair targets of SP2 exhibited all negative correlations in training response (Fig. 4b) between gene expression and chromatin accessibility and included the genes *Mpeg1* (Fig. 4e), *Gm2a* (Fig. 4f), *Bcl11b* (Fig. 4g), *Dennd1c* (Fig. S18a), and *Oas2* (Fig. S18b). Each gene and associated peak shared a similar pattern of training response as well, reflecting a decrease in gene expression and an increase in chromatin accessibility at early time points followed by a return to baseline. In SKM-GN, BMYB was the most significantly enriched TF, with 18.42% enrichment among DAR-DEG pairs vs. 3.67% general enrichment (p-value = 0.00042). Most DAR-DEG pair targets of BMYB exhibited a positive correlation in training response (Fig. 4c), including *Hjv* (Fig. 4h), *Fkbp5* (Fig. 4i), *Ppard* (Fig. 4j), *Cited4* (Fig. S18c), *Cab39* (Fig. S18d), *Sik1* (Fig. S18e). *Phlda3* was the only target that exhibited a negative correlation between gene and peak (Fig. S18f). A female-specific response early in EET is shared among *Hjv*, *Fkbp5,* and *Ppard*.

The circadian clock factor BMAL1 was the most enriched TF in liver with binding sites present in 14.47% of DAR-DEG pairs vs 5.42% general enrichment (p-value = 0.0026). Most DAR-DEG pair targets of BMAL1 were positively correlated in training response (Fig. 4d), of which *Abhd2* (Fig. 4k,l) and *G6pc1* (Fig. S18g,h,i) were correlated to multiple neighboring DARs. Other positively correlated targets include *Chp1* and *Onecut1* (Fig. S18j,k), while *Lpar3* was negatively correlated with a neighboring DAR (Fig. S18l). MAZ was identified as significantly enriched for DAR-DEG pairs in both SKM-GN (10.52% vs 3.43%, p-value = 0.0398) (Fig. S18m) and lung (26.67% vs 3.43%, p-value = 0.0014) (Fig. S18n). MAZ targets in SKM-GN included both positively and negatively correlated DAR-DEG pairs, including targets *Kcna7* and *Eif4g3,* which were positively correlated with neighboring DARs (Fig. S18o,p) and *Srsf2,* which was negatively correlated (Fig. S18q). MAZ targets in lung were all negatively correlated between gene and peak. They also overlapped with the gene targets of SP2 in lung, except for *Hspb6* (Fig. S18r), likely because of similarities in MAZ and SP2 binding sites.

### Correlated DMR-DEG pairs enriched for key TFs in adipose and lung tissue

We identified DMR-DEG pairs located within 500kb of each other and whose L2FC training responses were either positively or negatively correlated across time points and sexes (t-test of Pearson correlation < 0.05, Supplementary Table S3). The majority of DMR-DEG pairs were found in WAT-SC, 194 of 245 in total, while 26 were found in lung, 9 in heart, 7 in SKM-GN, 4 in liver, 3 in kidney, and 2 in hippocampus. The heavy skew towards WAT-SC is accounted for by the increased numbers of both DEGs and DMRs found in the tissue. We identified 14 TFs whose motifs were significantly enriched among the DMRs in correlated neighboring DMR-DEG pairs in each tissue (Fig. 4M). Seven enriched TFs were found in WAT-SC, while 5 were found in lung, and two in SKM-GN. SP2 was the most enriched TF in SKM-GN with motifs found in 57.14% of DMR-DEG pairs vs 13.65% general enrichment (p-value = 0.00863). As was seen with DAR-DEG pairs in lung, SP2 targets were all negatively correlated between gene expression and DNA methylation in their training response (Fig. 4N). Zinc finger *Zdhhc1* was negatively correlated with a DMR that included an SP2 binding site (Fig. S19d), while *Slc26a10* was negatively correlated with two DMRs containing SP2 binding sites.

NF1-halfsite was the most significantly enriched TF in WAT-SC with motifs found in 21.64% of DMR-DEG pairs vs. 8.08% general enrichment (p-value = 0.000027) (Fig. S19a). Targets included *Erg* (positive correlation, Fig. S19e), and *Kcnj15* (negative correlation, Fig. S19f). The DMR chr2-375476_cluster1 was correlated with five DEGs, one positively correlated: *Syt11* (Fig. S18g), and four negatively correlated: *Crabp2* (Fig. S19h), *Paqr6* (Fig. S19i), *Tmem79* (Fig. S19j) and *Rab25*. NF1-halfsite was also significantly enriched in lung with motifs found in 33.33% of DMR-DEG pairs vs 8.08% general enrichment (p-value = 0.0124). The TF AP-2gamma was also identified as significantly enriched in lung with motifs found in 33.33% of DMR-DEG pairs vs 9.13% general enrichment (p-value = 0.0189). There was considerable overlap in DMR-DEG targets for both NF1-halfsite and AP-2gamma including the DMR chr7-140730_cluster4 which is correlated with 4 DEGs, three positive correlations: *Agap2* (Fig. S19k), *B4galnt1* (Fig. S19l), *Arhgap9* (Fig. S19m), and one negative correlation: *Avil* (Fig. S19n). DMR chr4-156530_cluster11 was identified as a target of AP-2 gamma, but not NF1-halfsite. This DMR was negatively correlated with two transmembrane protein encoding DEGs: *Tmem176a* (Fig. S19o), and *Tmem176b* (Fig. S19p).

### Characterization of TF expression responses to EET

As putative transcriptional regulators were inferred from DAR-DEG and DMR-DEG correlations in a restricted number of tissues, we sought to independently characterize TF expression responses to EET per tissue over the 8-week training period. We measured the RNA levels of all TF-encoding genes and assessed their protein abundances and phosphorylation levels based on mass spectrometry data from a subset of 6 tissues. Various subsets of TFs exhibited significant changes at the transcriptome (Fig 5a), proteome (Fig 5b), and phosphoproteome (Fig 5c) levels in each tissue. Most significant TF changes were tissue-specific (Fig. S20); however, overlaps at the transcriptomic level were identified between SKM-GN and heart, and between BAT and WAT-SC, and at the phosphoproteomics level between heart, liver and lung. BAT and WAT-SC had the largest and most significant changes in TF gene expression (Fig 5a), which included immune-related TF genes such as *Irf8* and *Pou2f2* responding in both tissues and *Irf1* and *Spi1* specifically in BAT. Transcript levels of *Egr1* decreased significantly across multiple tissues, including SKM-GN, heart, kidney, and lung. Fellow early response gene *Fos* also decreased across the same tissues, reaching significance in kidney. *Myb* expression was significantly altered both in HIPPOC, where transcript levels increased in response to training in both sexes, and in kidney, where transcript levels increased in females but decreased in males.

**Figure 5:**
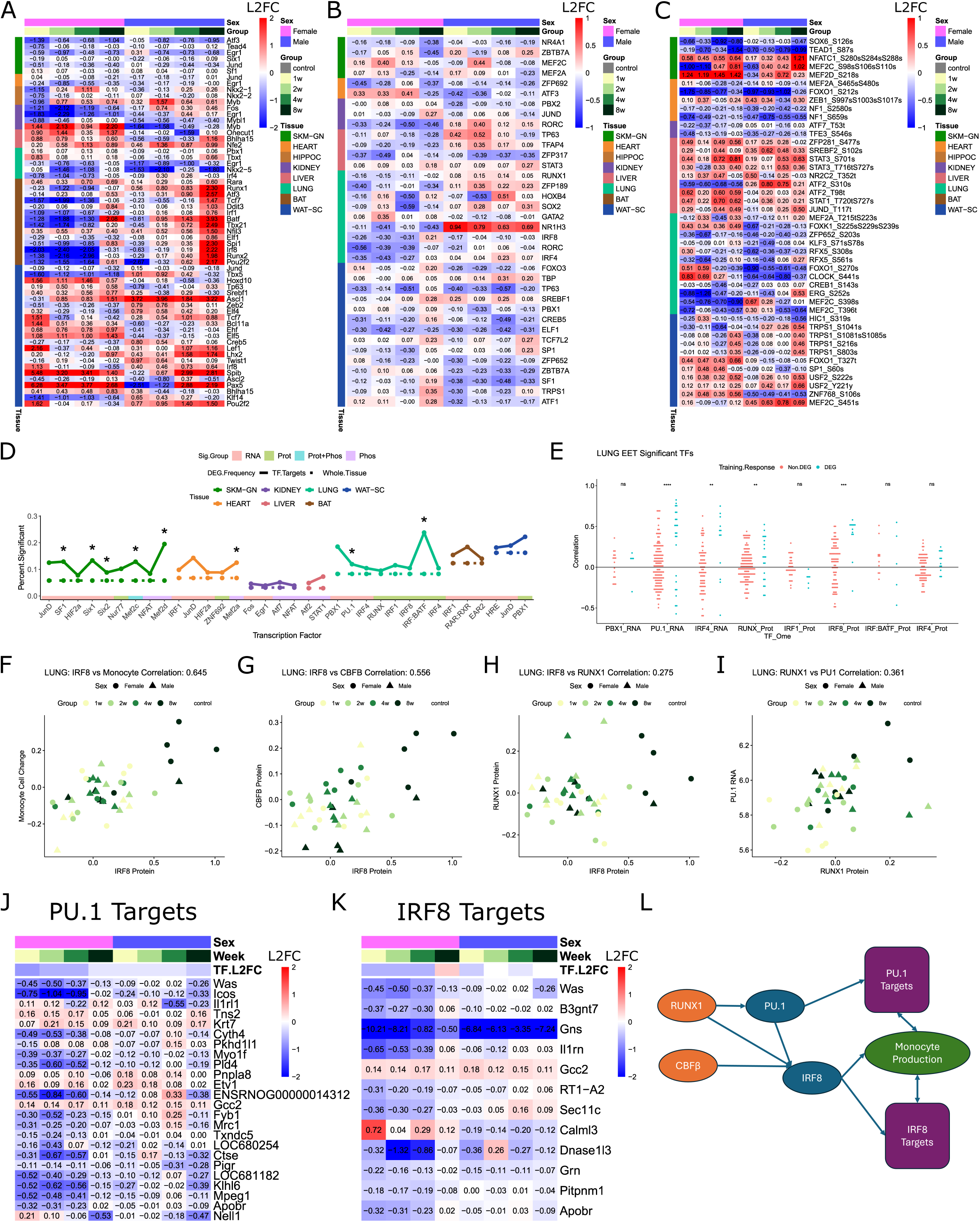
TFs showed significant EET responses at multiple omic levels. (a-c) Significant TF training responses at the transcriptomic (a), proteomic (b) and phosphoproteomic (c) levels. All TFs satisfy p-value < 0.1 in the specified ome and L2FC values are displayed at each time point and sex. Tissue of significance is color-annotated on the left side of each row. (d) TFs with significant changes at the transcriptomic, proteomic and phosphoproteomic levels and whose proximal promoter motif targets are enriched for DEGs are shown. Significant enrichments (p < 0.05) are represented with a *. Solid lines reflect the frequency of TF proximal promoter motif targets to be DEGs while the dashed lines show the frequency of DEGs among expressed genes in each tissue. Lines are colored by the tissue of TF training response and target DEG enrichment and TFs are individually colored by the source of their TF training response (RNA, Protein, Protein and Phosphoprotein, and Phosphoprotein). (e) Correlation of TFs with DEG and non-DEG targets in lung. * represents p < 0.05 of t test for difference between correlation of TF with DEG targets and non-DEG targets. PU.1 and IRF8 showed the most significant correlation with their DEG targets. (f-i) Correlated EET responses in lung of IRF8 and monocyte composition (f), IRF8 and CBFB (g), IRF8 and RUNX1 (h), and RUNX1 and PU.1 (*Spi1*) RNA (i). (j-k) Lung EET L2FC responses of DEG targets of PU.1 (j) and IRF8 (k). TF L2FCs in lung are included as column annotations. (l) Schematic of RUNX1 and CBFB-induced monocyte production pathway.

WAT-SC and lung exhibited the most significant changes in TF protein abundance (Fig. 5b). Select TFs showed significant protein level changes in multiple tissues, including RORC which decreased in female subjects in both lung and kidney. PBX1 decreased in WAT-SC while PBX2 decreased in kidney, and ATF1 and ATF3 increased in male subjects in WAT-SC and heart, respectively. NR4A1 (NUR77) and MEF2A levels decreased in SKM-GN at week 8 of training, while MEF2C levels increased at week 2 of training.

MEF2C showed significant protein phosphorylation changes in SKM-GN, lung, and WAT-SC (Fig 5c). Other MEF family members, MEF2A and MEF2D, also exhibited significant changes in phosphorylation in lung and SKM-GN, respectively. A number of TFs had multiple significant phosphosite changes in a single tissue, including two NF1 phosphosites in heart, two STAT3 sites and two ATF2 sites in liver, two RFX5 and two MEF2C sites in lung, and two USF2 sites and four TRPS1 sites in WAT-SC.

The majority of significant EET responses in TFs were ome-specific, with a few exceptions primarily in WAT-SC (Fig. S21). SREBF1, IRF8, FOXO3, FOXK1, and TCF7L2 all had significant EET responses in WAT-SC and were positively correlated in their transcriptomic and proteomic responses. We also identified positive correlations in RNA-Protein EET response in STAT4 in lung, RXRA and STAT3 in liver, and a positive correlation in RNA-Phosphoprotein EET response in CEBP6 in heart.

Now that we determined which TFs were found to have significant training responses at either a transcriptome, proteome or phosphoproteome level, we wanted to find which of their motifs were enriched among the promoter regions of DEGs, identifying potential training-induced regulatory relationships. We found a number of TFs whose motifs were enriched among DEGs (Fig 5d, Fig S22-S23), including eight statistically significant motif enrichments.

SF1 (Fig. S24a), SIX1 (Fig. S24b) and SIX2 all exhibited a significant training response at the transcriptome level in SKM-GN and their motifs were significantly enriched in target DEGs. The FOS motif was not significantly enriched in target DEGs in kidney; however, four of its target DEGs exhibited a similar EET response to FOS itself (Fig. S24c). A JUND motif was enriched in target DEGs in SKM-GN (Fig. S24d), heart (Fig. S24e), and WAT-SC (Fig. S24f), a PBX1 element was enriched in lung (Fig. S24g) and RAR:RXR was enriched in BAT (Fig. S24h).

MEF2C had significant EET responses in SKM-GN at both the proteome and phosphoproteome level and its motif was significantly enriched in target DEGs (Fig. S25a,e). IRF:BATF was significantly enriched in DEG targets in lung while the heterodimer also exhibited a significant EET response at the proteome level (Fig. S25b). Other TFs with significant EET responses in protein abundance and motif enrichment in DEG targets included NR4A1 (Nur77) in SKM-GN (Fig. S25c), and PBX1 in WAT-SC (Fig. S25d). *Dll4* was found to be a DEG target of PBX1 in both WAT-SC and lung. MEF2A responded significantly to EET at the phosphoproteome level and its motif was significantly enriched in target DEGs in heart (Fig. S25f), while binding sites for ATF7 in kidney (Fig. S25g) and STAT1 in liver (Fig. S25h) were also enriched in DEG targets.

SIX1 DEG targets in SKM-GN exhibit a range of functions and training responses, including collagen gene *Col3a1*, and muscle-contraction associated gene *Lmod1*, both of which decreased in response to training. Protein modification-associated genes *Golga4* and *Art1* increased over training, while ubiquitin gene *Usp2* decreased. Malic enzyme *Me3*, which is involved in the oxidative decarboxylation of malate to pyruvate, is consistently expressed at higher levels during training. MEF2C protein levels significantly increased at week 2 of training and then returned to baseline, while phosphorylation increased significantly throughout the eight weeks of training. Among MEF2C target genes, clock gene *Per1* demonstrated the highest increase in expression at the onset of training, while *Sema6c*, *Ankh*, *Ptpn1*, and *Phkg1* exhibited decreased expression following training. *Dystrophin* (*Dmd*), a critical protein for muscle fiber integrity^24^, was the most negatively correlated with MEF2C protein level changes, but positively correlated with MEF2C phosphorylation changes. IRF:BATF targets in lung predominantly exhibited decreased expression following training, including tubulin gene *Tuba1c*, mitochondrial biogenesis-associated gene *Perm1,* and actin cytoskeleton organizational gene *Cfl1*. NR4A1 (NUR77) significantly decreased in protein level in SKM-GN, as did the majority of its DEG targets including heat shock protein *Hspa1l* and dual specificity phosphatase *Dupd1*.

We calculated the sample-level correlation in EET response between significant TFs and their target genes, and identified TFs whose DEG targets were most likely to be correlated to TF response (Fig. S26). NR4A1 and SIX2 in SKM-GN and JUND in heart were significantly correlated with DEG targets (Fig. S26a,b); however, the strongest correlations were found in lung (Fig. 5e). TFs PU.1 and IRF8 were found to be the most significantly correlated with DEG targets, with IRF4 and RUNX also achieving significance. IRF8, RUNX, and *Spi1*, the gene encoding the PU.1 TF, all share similar patterns of EET response, namely a decrease in expression most pronounced in female subjects and early training time points. Indeed, this pattern is positively correlated with the decreased proportion of monocytes identified in lung by cell type decomposition analysis (Fig. 5f). IRF8 is known to act on monocyte differentiation, and CBFB and RUNX have been identified as upstream regulators of IRF8 activity^25^. Indeed, CBFB knockouts in mice blocked IRF8 expression and monocyte differentiation, with monocyte activity being restored upon reintroduction of IRF8. Previous mouse work showed that RUNX1 also acts as an upstream regulator of PU.1^26^. These regulatory relationships have not been previously characterized in EET response. However, our data reveal positive sample-level correlations in EET response between IRF8 and CBFB protein amounts (Fig. 5g), IRF8 and RUNX1 protein amounts (Fig. 5h), and between RUNX1 protein and PU.1 *(Spi1*) transcript levels (Fig. 5i). The majority of PU.1 DEG targets (Fig. 5j) and IRF8 DEG targets (Fig. 5k) show similar correlations with their respective TFs. In light of prior knowledge and our current findings, we propose a pathway involving the activation of IRF8 by RUNX1 and CBFB, the activation of PU.1 by RUNX1, and, via their gene targets, the induction of differentiation of myeloid progenitors into monocytes (Fig. 5l). In our study of EET response, we found this pathway inhibited at the level of RUNX1 and CBFB (Fig. 5g,h), potentially producing a cascade of inhibition that leads to a decrease in monocyte abundance in the lung in females following exercise training.

### DARs vs. DMRs vs. DEGaPs show distinct TF motif enrichment patterns that differentially correlate with TF gene expression

The lack of nearby DARs or DMRs for the majority of DEGs within each tissue (see Fig. 3a,b) led us to hypothesize that DARs, DMRs, and DEGaPs may mediate different paths of transcriptional regulation: i) DARs and DMRs coordinating a combination of gene-proximal and gene-distal cis-regulatory interactions, ii) a combination of statically open cis-regulatory elements (DEGaPs) and changes in TF behavior influencing differential gene expression. To address this, we analyzed TF binding site enrichment at either DARs or DEGaPs relative to all open chromatin peaks in each tissue, or at DMRs relative to all methylated CpG sites in each tissue (Fig. 6a,b,c; see Methods).

**Figure 6:**
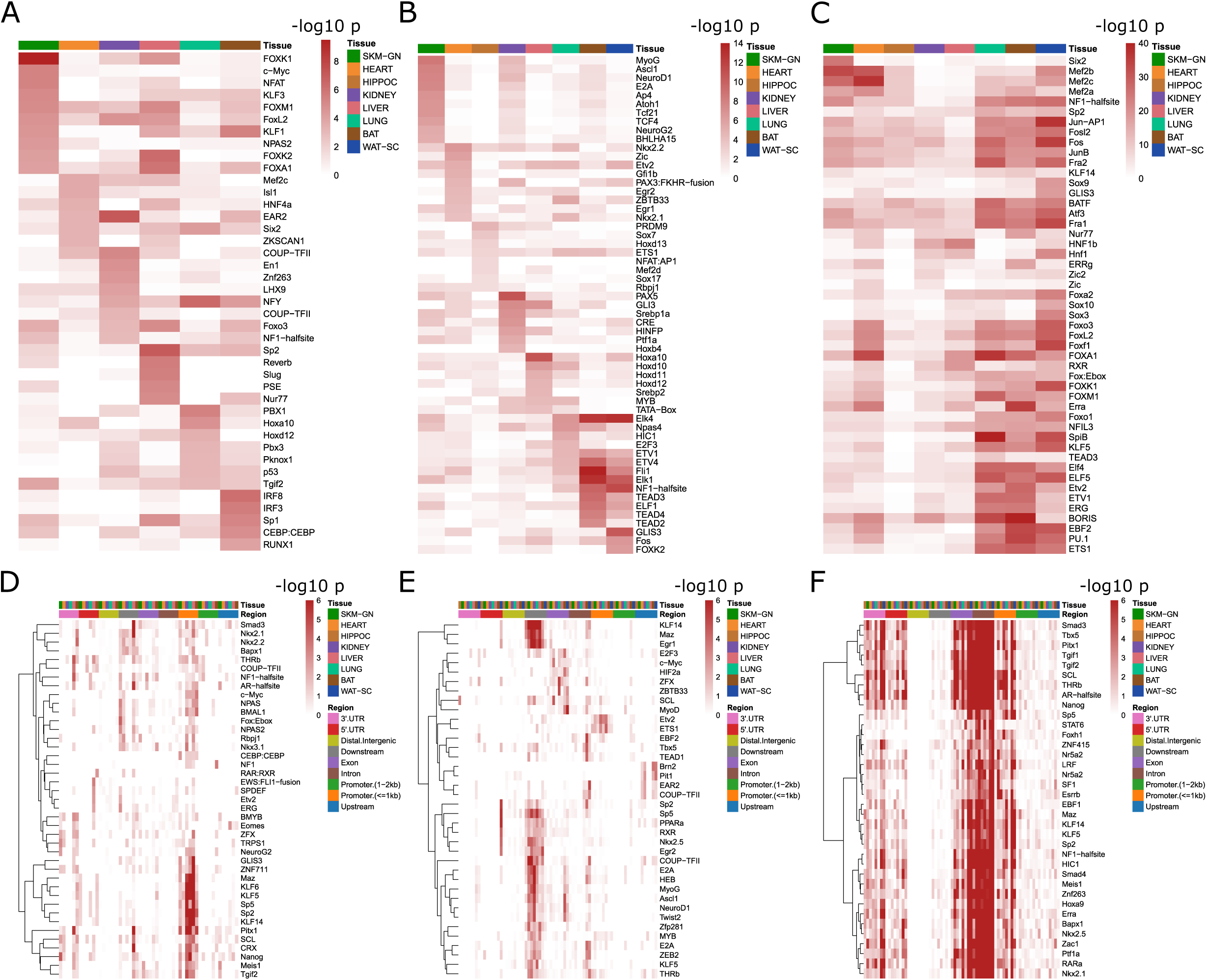
TF motif enrichment at the tissue level and at the genomic feature level in each tissue. (a-c) -log10 p-value of TF motif enrichment in tissue DARs (a), DMRs (b), and DEGaPs (c). Top enriched TFs were selected for each tissue in (a-c). (d-f) -log10 p-value of TF motif enrichment among sets of DARs (d), DMRs (e), and DEGaPs (f) split by genomic feature. Distinct sets of TF motifs are significantly enriched in proximal promoter regions (<=1kb from TSS) and downstream regions, shared by multiple tissues for DARs and DMRs, while TFs are most heavily enriched among intronic DEGaPs in most tissues.

Due to their very low number of DARs, WAT and HIPPOC were removed from the analysis (Fig. 6a). Motif enrichment patterns varied greatly across the six remaining tissues, reflecting a high degree of tissue-specificity. Motifs for both FOX and KLF families of TFs were over-represented among SKM-GN DARs. KLFs are zinc-finger TFs that have been associated with myogenesis and muscle fusion via their recruitment to Muscle Creatine Kinase (MCK) promoters^27^. Motifs enriched in DARs included SIX2 and MEF2C in heart, COUP-TFII in kidney, and SP2 in liver.

HOXA10 and HOXD12 motifs were enriched in lung, while both IRF8 and IRF3 motifs were enriched in BAT. Tissue-specific patterns of motif enrichment were maintained when measuring the frequency of motif presence in DARs across tissues (Fig S27a).

A similar pattern of tissue-specific motif enrichment was observed among DMRs for each tissue (Fig 6b), albeit with different TF motifs than in DARs. A MYOG motif was the most significantly enriched among DMRs in SKM-GN, along with binding motifs for two neuronal TFs NEUROD1 and NEUROG2. Another motif for nervous system-associated TF NKX2.2 was the most significantly enriched in heart. A PRDM9 motif was the most enriched in HIPPOC, as was PAX5 in kidney, and multiple HOX TF motifs in liver including HOXA10 and HOXD12, which were also enriched among DARs in lung. TF motif enrichment significance among DMRs overlapped more between lung, BAT and WAT-SC with ELK4, ELK1, ETV1, and ETV4 being enriched across the three tissues. GLIS3 motif enrichment was restricted to WAT-SC. As with DARs, the frequency of motif presence in DMRs across tissues maintained tissue specificity and even more sharply distinguished lung, BAT and WAT-SC tissues (Fig S27b).

Motif enrichment patterns among DEGaPs differed considerably from those in DARs and DMRs (Fig. 6c, S27c). Indeed, pairwise motif enrichment comparisons between DARs, DMRs, and DEGaPs per tissue were weakly correlated (<0.39; Fig S28, Fig S29, Fig S30). Motif enrichment significance in DEGaPs was greater in lung, BAT, and WAT-SC, presumably due to their higher proportions of DEGs (see Fig. 1b). MEF2 TF motifs were enriched across lung, BAT, and WAT-SC, as well as in SKM and heart, forming one cluster; on the other hand, ETS and ELF TF motifs were more exclusively enriched in lung, BAT, and WAT-SC and formed another cluster. MEF2 TFs are typically involved in muscle tissue regeneration^28^. ETS and ELF TFs are associated with the regulation of immunity^29,30^, suggesting that they may be related to the immune cell type composition changes occurring in adipose tissues. With respect to tissue-specific motif enrichment, sites for immediate early gene-encoded JUN and FOS TFs were enriched in HIPPOC (Fig S27c), HNF1 and ZIC families of TFs in kidney, and FOX and SOX families of TFs in liver.

To establish correlations between the motifs enriched at either DARs, DMRs, or DEGaPs and the relative expression levels of the corresponding TFs, we examined TF gene expression patterns in control tissues (Fig S31a, Fig S31b, Fig S31c, respectively) and L2FC in TF gene expression following EET (Fig S32a, Fig S32b, Fig S32c, respectively). We found a stronger correlation between motif enrichment and TF control gene expression levels among DEGaPs (Fig S33g-i) than among DARs (Fig S33a-c), and consistently found a negative correlation consistently among DMRs (Fig S33d-f). Conversely, there was no correlation between motif enrichment and L2FC in TF gene expression, be it among DARs (Fig. S27a,S32a), DMRs (Fig. S27b,S32b), or DEGaPs (Fig. S27c,S32c). These findings suggest that the influence of DEGaPs on DEG expression is more dependent upon the expression level of the associated TFs, whereas DARs and DMRs can influence DEGs directly, supporting the concept that DEGaPs vs. DARs and DMRs mediate distinct paths of transcriptional regulation.

### DARs, DMRs, and DEGaPs show cross-tissue motif enrichment conservation at specific genomic regions

The different genomic distributions of DARs and DMRs (see Fig. 1h,j) also suggested that some regions may contribute more to the regulation of EET gene responses than others. Thus, following the mapping of tissue DARs, DMRs, and DEGaPs to various genomic regions (i.e. promoter, upstream, downstream, 3’UTR, 5’UTR, intron, exon, distal intergenic), we examined motif enrichment within each subcategory of DARs, DMRs, and DEGaPs. We found that tissues shared conserved patterns of TF motif enrichment at the DARs mapped to the proximal promoter, downstream, and 3’UTR regions (Fig 6d). Proximal promoter DARs showed strong enrichment for SP and KLF TF motifs, MAZ and PITX1. Downstream DARs were enriched for binding sites for NPAS and BMAL1, two core circadian clock TFs, as well as NKX TFs. The NF1 subcluster also showed enrichment among 3’ UTR DARs, while THRB was most highly enriched among 3’ UTR DARs.

In contrast, TF motifs were most highly enriched among downstream DMRs relative to other genomic regions (Fig 6e), including multiple KLF TFs, MYOG, MAZ, and EGR1. ETV2 and ETS1 motifs were enriched among proximal promoter DMRs, while BM2 and PIT1 were enriched among upstream DMRs. Another cluster of TF motifss were enriched among exon DMRs including c-MYC, HIF2A, and MYOD. TF motif enrichment among DEGaPs was most significant in intronic regions, one of the regions most populated with DEGaPs (Fig 6f).

However, there was no enrichment among distal intergenic DEGaPs despite their large proportion of total DEGaPs. TF motifs enriched among intronic DEGaPs included SCL, SMAD3, THRB and ERRA. Significant enrichment was also found among exonic DEGaPs, which was highly correlated to intronic DEGaP enrichment. While also highly enriched among intronic DEGaPs, the most enriched TF motifs among promoter DEGaPs included MAZ and multiple KLF family members. Overall, genomic features appeared to be a driving force in clustering the enriched binding motifs among DEGaPs, DARs, and DMRs.

### Differential TF motif enrichment at DEGaPs of upregulated vs. downregulated DEGs

An associated MoTrPAC paper^6^, found that the biological pathways that were over-represented among upregulated vs. downregulated DEGs in each tissue after 8 weeks of training were distinct, suggesting that different sets of TFs were involved in their regulation. We sought to predict the sets of TFs that preferably bind to up-versus down-regulated genes. After identifying the DEGaPs related to up-regulated DEGs and those linked to downregulated DEGs in each tissue, we refined each DEGaP category into distal intergenic, intron, and promoter peak subsets (the most prevalent genomic features among peaks, as shown in Fig. 1g), and determined their respective patterns of TF motif enrichment. We then generated z scores for each TF enrichment across the peak sets, and applied hierarchical clustering to the TFs. We found that in most tissues, clustering was mainly driven by differences in TF motif enrichment between the promoter peaks associated with up-regulated DEGs vs. those associated with down-regulated DEGs (Fig S34). Adipose tissues had the greatest motif enrichment overlap between promoter peaks associated with up-regulation and those associated with down-regulation.

Restricting the TF motifs enriched within each subset of promoter peaks (i.e. promoter peaks associated with either up- or down-regulation) to those conserved across all four time points (Fig. S35), revealed some overlap in motif enrichment between the tissues (Fig. 7a,b). For instance, CLOCK, BHLHE41, and MYC (c-Myc) were among the TF motifs enriched across WAT-SC, heart, lung, SKM-GN, and kidney at the promoter peaks associated with upregulation; another set of TF motifs enriched primarily in heart and lung included ZFX and EBF2, while ATF2 was enriched in SKM-GN and kidney. Among the promoter peaks associated with downregulation, there was less consistent enrichment across tissues. However, the NKX2 motif was enriched across SKM-GN, kidney, liver, and lung, and the SOX9 binding site was enriched among 6 of the 8 tissues. Twenty seven TFs were shared between the 50 TF motifs associated with upregulation (Fig. 7a) and the 39 associated with downregulation (Fig. 7b), suggesting their involvement in both positive and negative regulation depending on the tissue. Heart, SKM-GN, and kidney shared the most enriched TF motifs associated with upregulation and lung, liver, and SKM-GN shared the most enriched motifs associated with downregulation (Fig 7c,d). Overall, while gene responses and TF expression responses were largely tissue-specific, tissues showed an overlap in the sets of TF motifs associated with either upregulated or downregulated DEGs, suggesting that specific patterns of EET-induced regulation could be conserved across tissues.

**Figure 7:**
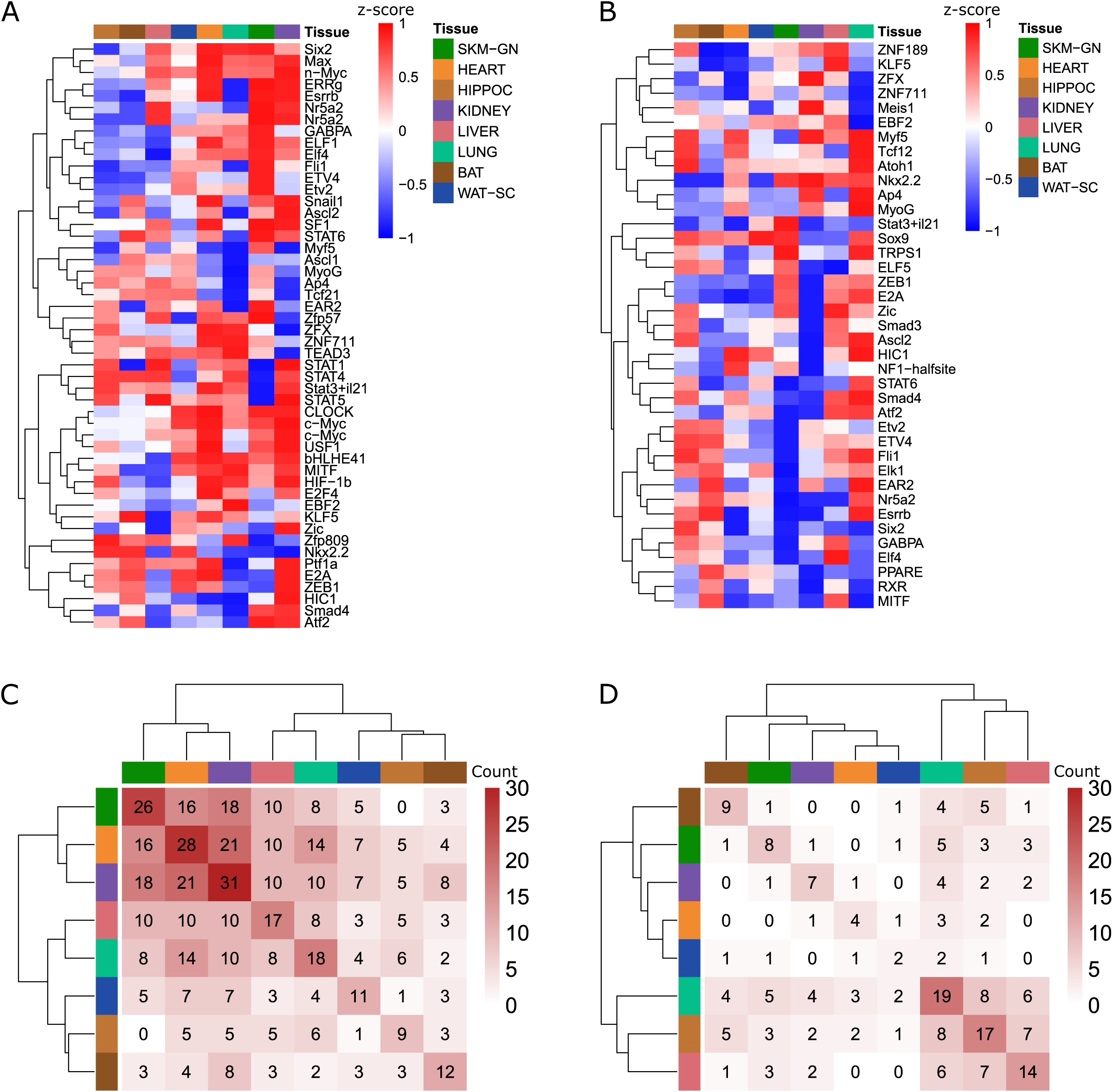
Tissues shared sets of TFs enriched in the promoter regions of genes that were up- or down-regulated after 8 weeks of training. (a-b) Heatmaps of relative enrichment of TFs among the promoter peaks of up- (a) or down-regulated genes (b) across tissues. (c-d) Number of enriched TFs shared between pairs of tissues in the promoter peaks of up- (c) or down-regulated genes (d).

### Connecting TFs to DEGs associated with EET-induced phenotypic changes

Exercise elicits various phenotypic changes such as increased aerobic capacity (VO2max) and reduced body fat mass, all of which reflect physiological adaptations. As several parameters were clinically measured throughout the EET period, we sought to: i) assess possible correlations between phenotypic alterations and gene expression responses to training, ii) infer key transcriptional regulators based on their motif enrichment at the accessible promoters of DEGs that correlated with phenotypic changes.

Parameters measured included weight, body fat mass, body lean mass, body water, lactate levels, and VO2 max (Fig. S36). Body weight and fat mass were significantly lower in 8-week trained rats than in controls (Fig S36a,b), with the greatest difference observed in males. VO2 max significantly increased in both females and males in response to EET (Fig S36c). Body lean mass also increased in both sexes, though significance was only reached in females (Fig S36d). Lactate (Fig S36e) and body water (Fig S36f) showed no significant changes relative to control, though a significant decrease in lactate levels was observed between week 4 and week 8 in males. Positively correlated parameters included VO2 max vs. body lean mass, and body weight vs. body fat mass (Fig 8a).

**Figure 8:**
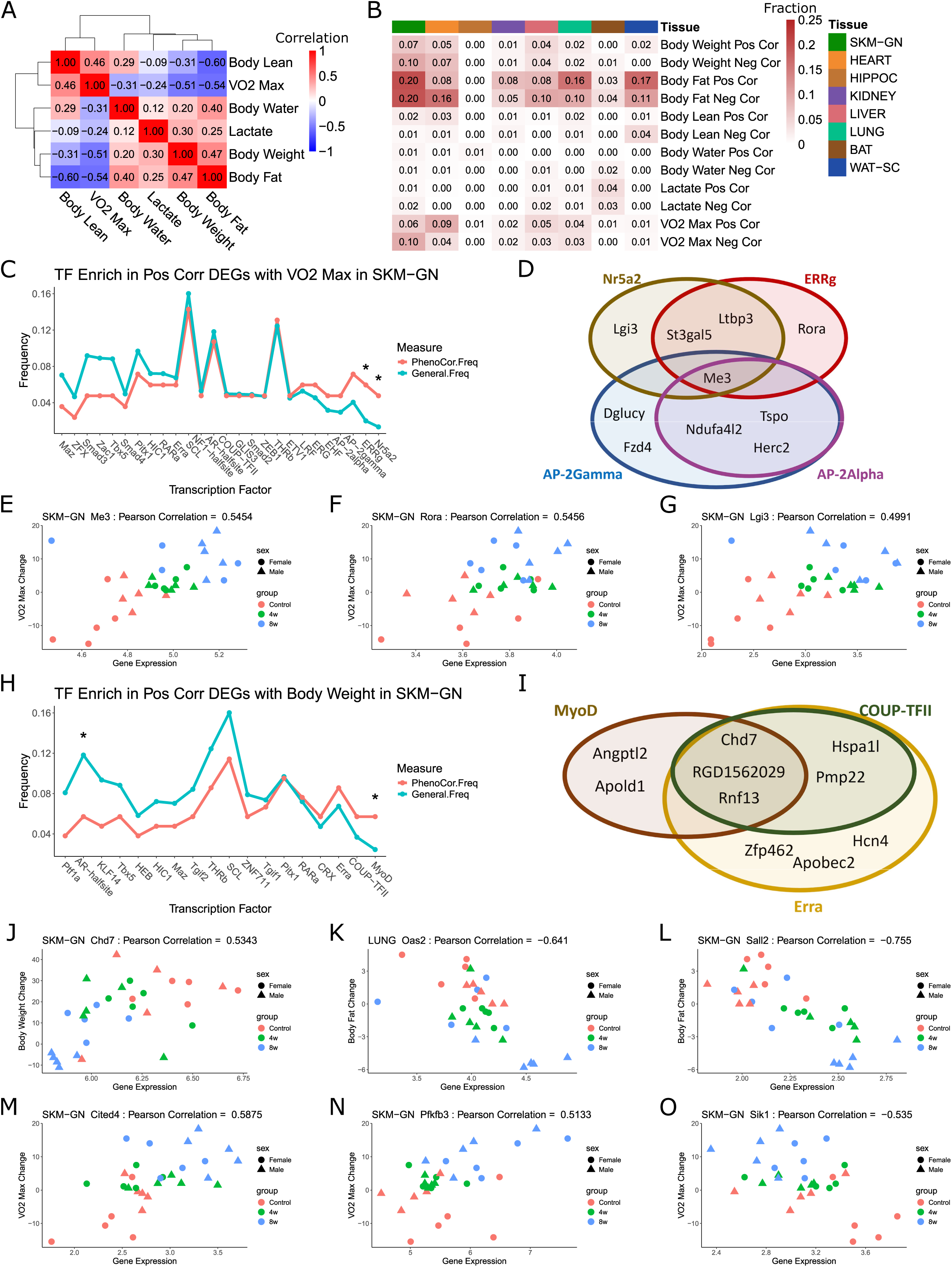
TF enrichment among phenotype-correlated DEGs. (a) Heatmap of Pearson correlation between phenotypic measures. Strong relationships between body lean and VO2 max and body fat and weight gain. (b) Frequency of tissue DEGs positively (> 0.5) or negatively (< -0.5) correlated with each phenotypic measure. (c) Comparison of TF enrichment among the active promoter peaks of VO2 max positively correlated DEGs vs TF enrichment among the total active promoter peak set in SKM-GN. * represents binomial test significance (p < 0.05) for difference in phenotype correlated DEG frequency and general frequency. (d) Overlap of target DEGs for most enriched TFs among VO2 max positively correlated DEG promoter peaks. (e-g) Scatter plots of gene expressed vs correlated phenotypic measure. In SKM-GN, VO2-max change is positively correlated with Me3 (e), Rora (f), and Lgi3 (g). (h) Comparison of TF enrichment among the active promoter peaks of body weight positively correlated DEGs vs TF enrichment among the total active promoter peak set in SKM-GN. * represents binomial test significance (p < 0.05) for difference in phenotype correlated DEG frequency and general frequency. (i) Overlap of target DEGs for most enriched TFs among body weight positively correlated DEG promoter peaks. (j-o) Scatter plots of gene expression vs correlated phenotypic measure. In SKM-GN, weight gain is positively correlated with Chd7 (j), and body fat change is negatively correlated with Oas2 (k), and Sall2 (l). In SKM-GN, VO2 max change is positively correlated with Cited4 (m) and Pfkfb3 (n), and negatively correlated with Sik1(o).

In most tissues, DEGs formed 2 separate clusters based on their either positive or negative correlation with a measured parameter change (Fig S37). Correlation with body fat involved the highest proportion of DEGs in most tissues (Fig. 8b), except for BAT and HIPPOC.

To identify key regulators of the DEGs that were associated with phenotypic changes, we analyzed motif enrichment at the corresponding promoter DEGaPs. In SKM-GN, the promoter DEGaPs of genes that were positively correlated with changes in VO2 max were significantly enriched for NR5A2 and ERRG motifs (Fig. 8c). The VO2 max-correlated DEG targets of those two TFs and of AP-2GAMMA and AP-2ALPHA, whose motif enrichment did not reach significance, showed considerable overlap (Fig 8d). The positive correlation between three of the DEGs targeted by either NR5A2, ERRG, or/and AP-2GAMMA (*Me3*, *Rora*, and *Lgi3*) and VO2 max change is depicted in Fig. 8e-g.

In SKM-GN, we also found a positive correlation between DEGs and body weight changes and identified the MyoD motif as significantly enriched at the corresponding promoter DEGaPs (Fig. 8h). While an enrichment for the COUP-TFII and ERRA motifs was also detected, it did not reach significance (Fig 8h). The *Chd7, Rnf13,* and *RGD1562029* DEGs showed concurrent enrichment for the MyoD, COUP-TFII, and ERRA motifs (Fig 8i). The positive correlation between *Chd7* expression and body weight change is illustrated in Fig. 8j.

As SP2, MAZ and BMYB have been identified as major regulators of the DARs-associated DEGs (DAR-DEG pairs; see Fig. 4), we asked whether those DEGs correlated with phenotypic changes. *Oas2* expression in lung, a target of both SP2 and MAZ, was negatively correlated with body fat change (Fig. 8k), as was *Sall2* in SKM-GN, another identified MAZ target (Fig. 8l). Three BMYB targets in SKM-GN were significantly correlated with VO2 max change, including *Cited4* (Fig. 8m) and *Pfkfb3* (Fig. 8n) positively correlated, and *Sik1* (Fig. 8o) negatively correlated. In the significant body fat and body weight correlations, we noted sex differences both in EET-dependent RNA levels and in phenotypic changes, as week 8-trained males deviated more from controls than week 8-trained females (Fig. 8j-l). In view of the characterization of TF expression responses to EET, we next evaluated correlations between TF expression changes, at either the RNA or protein level (see Fig. 5) and phenotypic changes. We observed the strongest correlations between TF RNAs and body fat changes (Fig S38a), with *Pknox1* in SKM-GN being the most positively correlated, and *Srebf1* in the kidney being the most negatively correlated. *Srebf1* in the kidney was also the most positively correlated with body lean mass changes. *Rora* was the most positively correlated with VO2 max changes. *Zeb1* and *Zeb2* shared similar correlation patterns, namely a positive correlation with body fat in WAT-SC. *Gata2* had diverging correlation patterns depending on the tissue; it was positively correlated with body weight and body fat in WAT-SC, and negatively correlated with both measures in the lung. Similar to the correlations with TF RNA level-changes, the strongest correlations at the protein level were with body fat changes. Remarkably, liver showed the highest frequency of strong correlations (Fig S38b). In WAT-SC, JUN was the most positively correlated with body fat, while TCFL2 and EHF were the most negatively correlated. Similar to their respective transcripts, ZEB1 and ZEB2 proteins in WAT-SC both exhibited a positive correlation with body fat, while ZEB2 in lung was also positively correlated with body fat. IRF3 was the most positively correlated with body lean changes in SKM-GN, and CLOCK was negatively correlated with body fat in WAT-SC. Connecting EET-modulated genes and proteins, and specifically TFs, with correlated phenotypic changes suggest potential mechanistic roles. In particular, MAZ and SMAD3 target gene correlation with body fat changes reinforces their functional role in the training response.

## Discussion

Regular exercise has a variety of physiological benefits affecting many organ systems. This comprehensive study using the rat model allowed investigating the molecular mechanisms of EET in important tissues that otherwise are inaccessible in human participants. We integrated chromatin accessibility, DNA methylation, and transcriptome data from 8 rat tissues to infer the TFs underlying EET responses in each tissue. We found multiple layers of regulation characterizing EET adaptations, including utilization of the tissue-specific and genomic region-specific TF machinery, TF expression changes at the RNA and protein levels, and post-translational phosphorylation of TFs. We also identified direct interactions between proximal promoter DARs or DMRs and assigned DEGs, long-range interactions between highly correlated DAR-DEG or DMR-DEG pairs enriched for known TFs, and specific patterns of training response with distinct TF motif enrichments, some conserved across tissues (Fig 9). We identified SP2, BMYB, and BMAL1 as key regulatory TFs in lung, SKM-GN, and liver, and found the down-regulation of immediate-early response genes over EET and enrichment of KLF and SP TF motifs among proximal promoter DEGaPs. In addition, we identified MEF2A, MEF2C, and MEF2D TFs with significant changes in protein level or phosphorylation whose motifs were enriched in DEGs in SKM-GN and heart. We also uncovered a pathway of correlated TFs in lung, involving down-regulation of CBFB and RUNX1 that, via PU.1 and IRF8, could be decreasing monocyte populations following EET. We found that the DEGaPs of upregulated vs. downregulated DEGs show differential motif enrichment, which predominantly occurs at promoter regions. Finally, we identified TFs associated with EET-induced phenotypic changes, including VO2 max and body weight.

**Figure 9:**
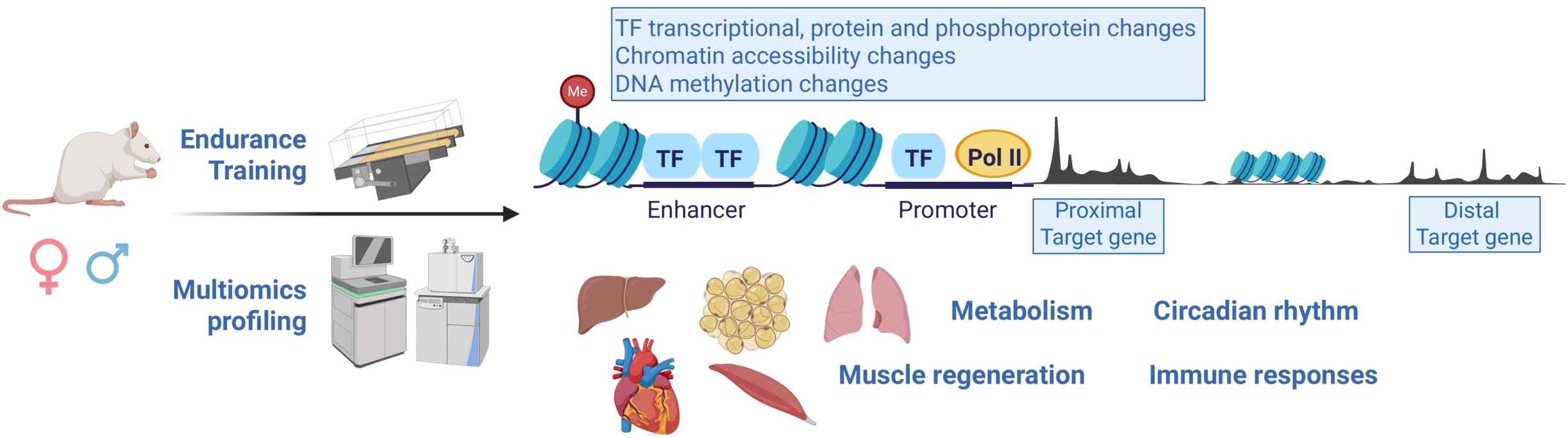
Transcription factors regulate exercise training-induced gene expression via multiple methods. Following eight weeks of endurance training, multiomic analysis across eight tissues have noted gene regulation through direct proximal promoter DAR and DMR to DEG relationships, DAR and DMR to distant correlated DEG relationships, tissue-specific or gene-region-specific TF machinery, and through changes in TF expression and phosphorylation within a specific tissue. Enriched TF gene targets are associated with metabolism, muscle regeneration, immune responses and circadian rhythm pathways.

The role of IRF8 in monocyte differentiation has been previously studied^25^; however, it has not been previously connected to exercise response. Here, we identify the same regulatory connections between RUNX1, CBFB, IRF8, and PU.1 within this novel context in females. The female-specific nature of the downregulation of these factors and associated decrease in monocyte proportion is quite striking and highlights inherent sex differences in EET response. Furthermore, through our work, we have identified a number of downstream correlated targets of both IRF8 and PU.1, many of which have known immune or monocyte functions, that could potentially be involved in monocyte differentiation.

Sex differences in baseline tissue behavior and EET response arise throughout our analysis. WAT-SC exhibits different cell type compositions in male and female rats, while the cell type composition changes in response to EET in BAT and lung are also sex-specific. This is mirrored in the differential analytes identified in this study. Sex differences in EET response for DARs and DMRs were very common and, even in RNA-seq where certain tissues like SKM-GN, heart, and kidney were more consistent across sexes in EET response, the remaining five tissues exhibited considerable sex differences. Sex differences are also prevalent in the phenotypic data with body fat decreasing more significantly in male rats than female rats over training. This highlights the necessity of designing experiments with a large sample size of diverse subjects to ensure important sex-specific distinctions are identified.

As seen with sex differences in EET response, most responses to EET are tissue-specific. The majority of differential responses to EET are unique to a tissue; indeed, the majority of DEGs and DARs are more highly expressed or accessible in their tissue of significant response than other tissues. This includes TFs where the majority of significant EET responses in TFs at a transcriptome or proteome level are tissue-specific. Beyond this, the majority of TF motif enrichments among DEGs are tissue-specific, with some interesting exceptions. JUND had significant consistent transcriptomic responses to EET in SKM-GN, heart, and WAT-SC, down-regulated in each tissue, but its DEG targets in each tissue are largely distinct. MAZ and NF1-halfsite were enriched for DAR-DEG and DMR-DEG pairs, respectively, in multiple tissues, while SP2 was enriched for DAR-DEG pairs in lung and DMR-DEG pairs in SKM-GN. In each case, the target pairs were distinct in the different tissues. These tissue-specific patterns are carried over to the pathway enrichment analysis, which was also mostly tissue-specific. This tissue specificity suggests that EET triggers unique signaling cascades within each tissue.

Normally, these responses use machinery unique to a given tissue; however, some more universal TFs function over multiple tissues, but through mechanisms unique to each tissue. This is a reminder of the importance of multi-tissue analysis to gain a fuller picture of a subject’s response to exercise.

The SP2 binding motif was enriched among both correlated DAR-DEG pairs in lung and correlated DMR-DEG pairs in SKM-GN. Across both comparisons, each target pair exhibited a negative correlation between pair members. SP2 has been shown to be essential for development in mice and fibroblast proliferation in mouse embryos^31^, and to negatively regulate cholesterol biosynthesis^32^. Here, DAR targets of SP2 include immune-associated factors: *Mpeg1* expresses a transmembrane protein in macrophages in an antimicrobial capacity^33^; *Oas2* is a type I-interferon response gene^34^; *Gm2a* affects T cell function through lipid exchange with CD1D^35^; *Bcl11b* regulates the early T cell differentiation stages^36^. Among the DMR targets of SP2, *Zdhhc1* has been connected with activation of the p53 tumor suppressor gene^37^. In fact, *Zdhhc1* has been found to be downregulated through DNA hypermethylation in tumors^38^. In response to EET, *Zdhhc1* is initially downregulated in early time points, but expression increases by week 8, coupled with hypomethylation of its negatively correlated DMR pair. In lung, DAR-DEG targets are enriched for the MAZ binding site. MAZ can act both as a transcriptional activator and a repressor^14,17^. MAZ is also over-represented in DAR-DEG targets in SKM-GN, with more immune-linked DEG targets, including *Ppp1r15a* (*Gadd34*), which has been found necessary for interferon production^39^. The MAZ-specific DEG target in lung, *Hspb6*, acts as a mediator of platelet aggregation along with smooth muscle relaxation^40^. This suggests that SP2 and MAZ could both be involved in the inflammatory response following exercise^41^.

The TF BMYB (MYBL2) is activated via phosphorylation during S phase of the cell cycle by CDK2^42^. BMYB is known to promote cell cycle progression through its downstream targets. The identified DEG target of BMYB *Pfkfb3* is known to promote cell cycle progression via regulating *Cdk1*^43^. *Pfkfb3* along with another BMYB target, *Sik1*, are associated with circadian rhythm factors^44,45^. *Hjv* is involved in iron metabolism and mutations in the gene can result in hemochromatosis^46^. *Cab39*, an oncogene, is also tied to cellular proliferation^47^. The majority of the BMYB targets showed increased expression in the early weeks of exercise training, followed by a return to baseline by week 8, suggesting initial stress responses from the shift into an active exercise lifestyle to which, over the course of EET, the body adapts. *Cited4,* an outlier to the BMYB target trend with expression continuing to increase over EET, can promote cardiomyocyte growth^48^.

The motif of AP2-gamma, also known as TFAP2C, was enriched among correlated DMR-DEG pairs in lung. The AP2-gamma motif was also significantly enriched among DEGs in SKM-GN that are correlated with a change in VO2max over EET. AP2-gamma’s primarily studied function is in early development morphogenesis^49^; however, it has also been shown to have an inflammatory role by activating Th17 and Th1 cells^50^. The NF1-halfsite motif was also enriched among correlated DMR-DEG pairs in lung, as well as WAT-SC. NF1 is categorized as a tumor suppressor protein and negatively regulates the RAS/MAPK pathway^51^. Mutations in the protein lead to neurofibromatosis type 1 which can have numerous negative effects including musculoskeletal defects and impaired exercise capabilities^52^. The regulatory relationships between target DMR-DEGs and both NF1 and AP2-gamma can be either positively or negatively correlated; however, we do identify multiple instances where single DMRs are correlated with multiple DEGs, including chr16-108674_cluster2 in WAT-SC negatively correlated with *Asah1*, *Pdgfrl*, and *Slc7a2*; and chr2-375476_cluster1 in WAT-SC negatively correlated with *Rab25*, *Crabp2, Tmem79*, *Paqr6,* and positively correlated with *Syt11*. This suggests that both binding motifs could act as regional regulators for multiple genes in response to EET.

Several TFs that were downregulated by EET in SKM-GN either at the RNA level (*Fos, Jun, Egr1,* and *Atf3*) or at the protein level (NR4A1) represent immediate-early response genes (IEGs), which were previously reported to be induced by acute exercise^53–56^. Stress-inducible ATF3 was shown to reduce the RNA expression of inflammatory chemokines and cytokines in mouse SKM following acute exercise, and ATF3 knockout resulted in impairment of some of the molecular adaptations to exercise training^53,57^. In SKM-GN, NR4A1 regulates genes associated with glucose uptake, glycogen synthesis, muscle growth,^58^ and inflammatory responses^59,60^. As exercise represents a physical stress, modulation of IEGs may help tissues like SKM-GN to recover their homeostasis and thus facilitate their adaptation to exercise training. The two gene targets of NR4A1 with the largest correlated decreases in EET-response gene expression in our study were the heat shock protein *Hspa1l*, a critical element of the cellular stress response^61^, and *Dupd1* (*Dusp29*), which has been linked to the regulation of muscle cell differentiation, development, and atrophy^62^.

We found that binding sites for circadian clock TFs were over-represented at the promoter DEGaPs of DEGs that were associated with VO2 max changes in SKM-GN. Notably, there were enriched motifs for the nuclear receptors NR5A2 and ERRG^63^ which bind to the core clock proteins CLOCK^64^ and BMAL1^65^, respectively. BMAL1 itself was enriched for correlated DAR-DEG pairs in liver. Moreover, we found an over-representation of binding sites for circadian clock-related TFs at the promoter and downstream DARs (NPAS and BMAL1 motifs), and at the DEGaPs of upregulated DEGs (CLOCK motif) in multiple tissues, suggesting a cross-tissue, exercise training effect on circadian clock factors. It is noteworthy that the EET program in rats was conducted during an active, dark phase, thus excluding that the enrichment for circadian clock TF motifs would be related to exercise training in the inactive phase. Recent studies revealed interactions between exercise and circadian rhythms. Some demonstrated that time of exercise can modify the transcriptional response to acute exercise, while others showed that exercise can modify the muscle clock phase and expand the circadian transcriptome in SKM^66–71^.

Limitations of our study include the use of inbred rats, which eliminates effects of genetic diversity, and high variance in some tissues, thus reducing statistical power. Multiomic power analysis identifies that these limitations are especially present in the phosphoproteomics data. However, differences in cell type composition within testing groups of same strain, same age, same sex rats suggest greater interindividual variation than initially anticipated and stronger applicability to the diversity expected in human responses to EET. Exercise effects on cell type composition could only be addressed computationally in assays of bulk tissue. Many integrated multiomic EET responses that we identified, such as muscle development in SKM-GN, innate immunity in lung, increased immune cell type proportions in adipose tissues, and metabolic processes in liver may contribute to known health effects of exercise. The goal of the MoTrPAC project is to provide the first systematic compendium of exercise effects. Follow-up studies of the multi-organ roadmap of genomic regulatory responses that we have identified are warranted.

Our multi-omic analysis across rat tissues allowed us to map the epigenomic changes to the transcriptional changes occurring during EET and infer TFs driving training responses. By providing a view of the complex interplay between chromatin structure modifications, DNA methylation, gene transcription, and TF abundance and activity throughout EET, this work provides a comprehensive survey of the multi-organ gene control mechanisms underlying the effects of regular exercise. This insight helps further the goal of maximizing the benefits of exercise in healthy individuals and developing targeted exercise therapies for patients with disease or disability.

## Methods

### Animal study design

#### Animal care

Male and female Fischer 344 (F344) inbred rats from the National Institute on Aging were transported to the University of Iowa a minimum of 4 weeks prior to starting exercise training. Rats were housed with the same sex, 2 per cage (146.4 square inches of floor space) in ventilated racks (Thoren Maxi-Miser IVC Caging System) on Tekland 7093 Shredded Aspen bedding and fed the Lab Diet 5L79 pelleted diet.

Rats were acclimated to a reverse dark-light cycle with lights off at 9:00am and lights on at 9:00pm, with temperature controlled at 68-77°F and humidity of 25-55%. For these studies we will use Zeitgeber Time (ZT) to refer to time of day relative to the time of lights off/lights on with lights off at ZT12. All experimental interventions and husbandry were performed during the active, dark phase of the rats under red light. All animal procedures were approved by the Institutional Animal Care and Use Committee at the University of Iowa.

#### Treadmill familiarization

Prior to exercise training, rats were acclimated to treadmill exercise on a Panlab 5-lane rat treadmill (Harvard Instruments, Model LE8710RTS). Day 1-2 consisted of static treadmill exploration for 10 minutes. Days 3-5 consisted of running at 6m/min at 0° incline for 10 minutes, speed was increased to 10m/min between Days 6-12. On Day 12, rat running behavior was scored for compliance with running at 12m/min for 5 minutes at a 10° incline. Based on running behavior, rats received a score from 1-4 with 4 being the highest score. Rats that were assigned a score of 1 were removed from the study. 25 male and 25 female compliant rats were randomized to a control or one of 4 training groups, resulting in 5 male and 5 female rats per group.

#### Progressive exercise training protocol

At 6-months of age, 1, 2, 4 or 8 weeks of exercise training began. Control rats were placed on a static treadmill for 15 min per day, 5 times per week. Exercise training consisted of a progressive training protocol 5 times per week at ZT13-20, to target 70% VO2max (see below). Training times each day for each rat subject are included in Supplemental Table S4. Week 1 sessions started at 13m/min for males and 16m/min for females at 5° for 20 minutes, with duration increased by one minute each day until reaching 50 min on day 31 of training. The treadmill grade was increased from 5° to 10° at the start of week 3. The treadmill speed increased at the start of week 2 (15m/min males, 18m/min females), 4 (18m/min males, 21m/min females), 5 (20m/min males, 23m/min females), 6 (23m/min males, 26m/min females), and 7 (25m/min males, 28m/min females) and was fixed for the final 10 days of training. Rats performing less than 4 days of training per week were removed from the study and euthanized as described below.

#### Body composition measurements

Body composition (lean tissue%, fat tissue% and body fluid) was measured for all rats 13 days prior to the start of training and 5 days prior to euthanasia in the 4 and 8-week training groups using the minispec LF90II Body Composition Rat and Mice Analyzer (Bruker, 6.2 MHz Time Domain Nuclear Magnetic Resonance (TD-NMR) system). VO2max was determined prior to commencing training in all rats, and during the last week of training for rats in the 4- and 8- week exercise groups, in a single-lane enclosed treadmill (Columbus Instruments Metabolic Modular Treadmill), with rats acclimated two days prior to testing. For testing, the rat was placed in the treadmill and testing began once oxygen consumption stabilized. The testing protocol consisted of a 15 min warm up at 9 m/min and 0° incline. The incline was increased to 10° and treadmill speed was increased by 1.8 m/min every 2 minutes^72^. During the test, electric shocks were used if the rat stopped running and sat on the shock area. Testing stopped when the rat sat on the shock area 3 consecutive times and did not respond to increased shock. Blood was then taken from the tail to measure lactate. VO2max was determined as a leveling off of oxygen uptake, despite increased workload, a respiratory exchange ratio above 1.05, and an unhaemolyzed blood lactate concentration ≥6 mM.

#### Euthanasia and tissue collection

Rats in the 1 week training group were euthanized and had tissues collected for omic analysis after 1 week of training, while rats in the 2 week training group were euthanized and had tissues collected for omic analysis after 2 weeks of training. A similar pattern is found for the 4 week and 8 week training groups. The 8 week control group of rats were euthanized and had tissues collected after eight weeks, similar to the 8 week training group. On the day of euthanasia, food was removed at ZT11.5, 3 hours before tissue collections which took place between ZT14.5-17.5, a minimum of 48 hours post their last exercise bout. Rats were deeply anesthetized with approximately 1-2% isoflurane in oxygen, and gastrocnemius, white adipose, liver, lung, and heart were removed under anesthesia. Following removal of the heart, the rat was decapitated using a guillotine. The brain was removed from the skull and hippocampus dissected. The remaining tissues ( kidney, brown adipose, and hippocampus) were dissected post death. All tissues were cleaned of excess connective/fat tissue and immediately flash-frozen in liquid nitrogen, placed in cryovials and stored at -80°C. Rat tissues were archived and cryopulverized at the MoTrPAC Biospecimens Repository, until distributed to Chemical Analysis Sites for respective assays^5^. RNA-seq, ATAC-seq and RRBS analyses were conducted in all eight tissues and time points. Proteomics and phosphoproteomics were conducted in all time points for six of the eight tissues: SKM-GN, heart, kidney, liver, lung, and WAT-SC.

### Data production and quantification

#### ATAC-seq data generation

Nuclei from 30 mg white adipose, 15 mg brown adipose, and 10 mg of other tissue samples were extracted using the Omni-ATAC protocol with modifications^73^. The white adipose, brown adipose, and hippocampus tissues were processed using no-douncing nuclei extraction.

Cryopulverized tissue powder was incubated in the homogenization buffer for 10 min at 4°C, tubes inverted every 2-3 minutes. Other tissue powder was incubated in the homogenization buffer for 5 minutes on ice and dounced 10 times using pestle A and 20 times with pestle B. Nuclei were stained with DAPI and counted using an automated cell counter. 50,000 nuclei (or max. 500 µl nuclei) were added to 1 ml ATAC-RSB buffer and spun at 1000 g for 10 minutes, and the supernatant was removed.

The nuclei pellet was resuspended in 50 µl of transposition mixture and incubated at 37°C for 30 minutes with 1000 rpm shaking. The transposed DNA was purified using Qiagen MinElute Purification kits (Qiagen # 28006), and amplified using NEBnext High-Fidelity 2x PCR Master Mix (NEB, M0541L) and custom indexed primers^73^. 1.8x SPRIselect beads were used to clean the PCR reaction and remove primer dimers. The ATAC-seq libraries were sequenced on a NovaSeq 6000 using 2x50bp with 35 million pairs of reads per sample.

#### ATAC-seq data processing and normalization

Reads were demultiplexed with bcl2fastq2 (v2.20.0) and processed with the ENCODE ATAC-seq pipeline (v1.7.0) (https://github.com/ENCODE-DCC/atac-seq-pipeline)^74^. Samples from the same sex and training group were analyzed as biological replicates. Reads were aligned to genome rn6.0.96^75^ with Bowtie 2 (v2.3.4.3)^76^. Duplicate and mitochondrial chromosome mapped reads were removed. Peaks were called using MACS2 (v2.2.4)^77^, both from reads from each sample and pooled reads from all biological replicates. Pooled peaks were compared with the peaks called for each replicate individually using irreproducible discovery rate^78^ and thresholded to generate an optimal set of peaks. Optimal peaks from all workflows were concatenated, trimmed to 200 base pairs around the summit, sorted and merged with bedtools (v2.29.0)^79^ to generate a master peak list. This peak list was intersected with the filtered alignments from each sample using bedtools coverage to generate a peak by sample matrix of raw counts. Peaks from non-autosomal chromosomes were removed. Peaks that did not have at least 10 read counts in four samples in a tissue were removed. Filtered raw counts were then quantile-normalized with limma-voom^80^. This version of the normalized data was used for downstream analyses.

#### ATAC-seq peak assignment to genomic features

Accessible regions identified using ATAC-seq were assigned to one of the nine terms of genomic features using Ensembl Rn6 GTF (gene annotation file) and function annotatePeak of package ChIPseeker^81^ (v1.8.6).

Nine genomic features are defined as:

Promoter (<=1 kb) (proximal promoter): within +/-1 kb from the transcription start site (TSS); Promoter (1-2 kb): +/-1 to 2 kb from the TSS;

Upstream (<5kb) 2-5 kb upstream from the TSS;

Downstream (< 5 kb): within 5kb downstream of the transcription end site (TES);

5’ UTR (5’ untranslated region); Exon; Intron; 3’ UTR (3’ untranslated region);

Distal Intergenic: regions > 5kb downstream of TES or > 5kb upstream from next TSS; Overlaps Gene: overlaps with gene annotation, but not in any terms above.

All ATAC-seq identified accessible regions were assigned to the closest genomic feature of a genome. Differentially expressed gene associated peaks (DEGaPs) are defined as all accessible regions assigned to the differentially expressed gene.

#### DNA methylation data generation

Rat tissues were disrupted in GenFind v2 lysis buffer (Beckman Coulter, Indianapolis, IN) with a tissue bead ruptor (Omni International, Kennesaw, GA). Genomic DNA was extracted in a Biomek FxP automation workstation (Beckman Coulter, Chaska, MN). DNA sample quantification was done by Qubit HS dsDNA assay (Thermo Fisher Scientific) and quality was determined by Nanodrop A260/280 and A260/230 ratios. Reduced representation bisulfite sequencing (RRBS)^82^ libraries were generated with the Ovation® RRBS Methyl-Seq kit from Tecan Genomics (Baldwin Park, CA). Quantity of the libraries was determined by the Qubit HS ds DNA assay (Thermo Fisher Scientific) and quality evaluation was done by either Bioanalyzer High Sensitivity DNA Chip or Fragment Analyzer High Sensitivity NGS Assay (Agilent Technologies, Santa Clara, CA). Libraries were sequenced on a NovaSeq 6000 platform (Illumina, San Diego, CA) using paired-end 100 base-pair run configuration.

#### RRBS data processing and normalization

bcl2fastq (version 2.20) was used to demultiplex reads with options --use-bases-mask Y*,I8Y*,I*,Y* --mask-short-adapter-reads 0 --minimum-trimmed-read-length 0 (Illumina, San Diego, CA). FastQC (v0.11.8) was used to calculate pre-alignment QC metrics^83^ and reads were indexed and aligned to the Ensembl *Rattus norvegicus* (rn6) genome using Bismark (v0.20.0)^84^. Bowtie 2 (v2.3.4.3) was used to quantify the percent of reads that mapped to globin, rRNA and phix sequence contaminants and spike-ins^76^. Chromosome mapping percentages were calculated with SAMtools (v1.3.1)^85^ and NuGEN’s “nodup.py” script quantified PCR duplicates.

CpG sites were selected for downstream analysis if they exhibited methylation coverage >= 10 in all samples. Individual CpG sites were divided into 500 base-pair windows and clustered with the Markov clustering algorithm R package *MCL*^86^. Quantile normalization of sites/clusters was conducted separately on each tissue using R package *preprocessCore*^87^.

#### RNA-seq data generation

Rat tissue powders were further disrupted using Agencourt RNAdvance tissue lysis buffer (Beckman Coulter, Brea, CA) with a tissue bead ruptor (Omni International, Kennesaw, GA, # 19-040E). The total RNA was quantified using NanoDrop (ThermoFisher Scientific, # ND-ONE-W) and Ribogreen assay (ThermoFisher Scientific). Total RNA quality was determined by Fragment Analyzer Standard Sensitivity RNA analysis (Agilent Technologies, Santa Clara, CA).

200-300 ng total RNA was used for library generation. The Universal Plus mRNA-Seq kit from NuGEN/Tecan (# 9133) was used to select polyadenylated RNA. The generated sequencing libraries contain dual barcodes (i7 and i5) and UMIs (unique molecular identifiers) to accurately quantify the transcript levels. The RNA-seq libraries were sequenced on a NovaSeq 6000 using 2x100 bp with 35 million pairs of reads per sample.

#### RNA-seq data processing and normalization

Reads were demultiplexed with bcl2fastq2 (v2.20.0). Adapters were trimmed with cutadapt (v1.18). STAR (v2.7.0d) was used to index and align reads to genome rn6.0.96 and gene annotations^75^. Bowtie 2 (v2.3.4.3) was used to index and align reads to globin, rRNA, and phix sequences in order to quantify the percent of reads that mapped to these contaminants and spike-ins^76^. UMIs were used to accurately quantify PCR duplicates with NuGEN’s “nudup.py” script (https://github.com/tecangenomics/nudup). QC metrics from every stage of the quantification pipeline were compiled, in part with multiQC (v1.6)^88^. Lowly expressed genes (having 0.5 or fewer counts per million in all but one sample) were removed and normalization was performed separately in each tissue. These filtered raw counts were used as input for differential analysis with DESeq2^89^. To generate normalized sample-level data, filtered gene counts were TMM-normalized using edgeR::calcNormFactors, followed by conversion to log counts per million with edgeR::cpm^90^. The same normalization technique was used on the 8 week control samples of each tissue for cross-tissue comparisons.

#### Proteomics data generation

Liquid chromatography tandem mass spectrometry (LC-MS/MS) was conducted on six tissues: heart and liver at the Broad Institute and skeletal muscle, kidney, lung, and white adipose at Pacific Northwest National Laboratory (PNNL). Sample processing followed a modified version of a previous protocol^91^. Peptides were labeled using tandem mass tag (TMT)^92^ and samples were grouped into sex- and training time point-based TMT11 multiplexes. Multiplex samples were fractionated by high pH reversed phase separation. Heart and liver samples underwent online separation with a nanoflow Proxeon EASY-nLC 1200 UHPLC system (Thermo Fisher Scientific), and then analyzed with a Q-Exactive Plus mass spectrometer (Thermo Fisher Scientific). The remaining tissues’ samples underwent online separation with a nanoAcquity M-Class UHPLC system (Waters), and analyzed with a Q Exactive HF mass spectrometer (Thermo Fisher Scientific).

#### Phosphoproteomics data generation

Phosphopeptide enrichment was performed through immobilized metal affinity chromatography (IMAC)^5^. Phosphopeptides were eluted off IMAC beads in 3x70 µl agarose bead elution buffer, desalted with C18 stage tips, eluted with 50% ACN, and then lyophilized. Samples were reconstituted in 3% ACN / 0.1% FA for LC-MS/MS analysis. Heart and liver samples were separated by a nanoflow Proxeon EASY-nLC 1200 UHPLC system (Thermo Fisher Scientific) and analyzed with a Q-Exactive HFX mass spectrometer (Thermo Fisher Scientific). SKM-GN, WAT-SC, kidney and lung samples were separated by a Dionex Ultimate 3000 UHPLC direct-inject system (Thermo Fisher Scientific) then analyzed with a Q-Exactive HFX mass spectrometer.

#### Proteomics and phosphoproteomics data processing and normalization

For heart and liver, raw MS/MS data samples were processed by a Spectrum Mill (v.7.09.215) (Agilent Technologies). For the remaining tissues, sample processing was implemented by an in-house cloud-based proteomics pipeline executed in the Google Cloud Platform^5^. In all tissues, MS2 spectra were processed and searched against the rat RefSeq protein database (downloaded November 2018). Log2 TMT ratios to the common reference were used as quantitative values for all proteins and phosphosites. Principal component analysis and median protein abundance across samples were used to find sample outliers. Proteomics features that were not fully quantified in at least two plexes within a tissue and non-rat contaminants were removed. Median-centering and mean absolute deviation scaling of Log2 TMT ratios were done for sample normalization. Plex batch effects were removed using *limma::removeBatchEffect* function in R (v 3.48.0). Phosphoproteome data was not normalized to the total proteome due to the lack of complete overlap of phosphosites and total proteome features (80.5% -89.7%).

## Statistical analysis

### Differential analysis

Differential analyses were performed in each tissue of each ome. Males and females in one dataset were analyzed separately. Limma with empirical Bayes variance shrinkage was used for ATAC-seq, proteomics, and phosphoproteomics data^93^; the edgeR pipeline for methylation analysis was used for RRBS data^94^; DESeq2 was used for RNA-Seq^83^. For all proteomics and ATAC-seq data, the input for differential analysis was normalized as described above. For RNA-Seq, the input was filtered raw counts, in accordance with the DESeq2 workflow.

To select analytes that changed over the training time course, we performed F-tests (limma, *edgeR::glmQLFTest*) or likelihood ratio tests (DESeq2::nbinomLRT, lrtest) to compare a full model with ome-specific technical covariates and training group as a factor variable (i.e. sedentary control, 1 week, 2 weeks, 4 weeks, 8 weeks) against a reduced model with only technical covariates. For each analyte, male- and female-specific p-values were combined using Fisher’s sum of logs meta-analysis to provide a single p-value, referred to as the training p-value. To account for false discovery rate across all statistical tests, the training p-values were adjusted across all datasets within each ome using Independent Hypothesis Weighting (IHW) with tissue as a covariate^95^. Training-differential features were selected at 10% IHW FDR.

We used the contrasts of each training time point versus the sex-matched sedentary controls to calculate time- and sex-specific effect sizes, their variance, and their p-values (e.g., using linear F-tests), referred to as the timewise summary statistics. Specifically, for limma models we used limma::contrasts.fit and limma::eBayes, for DESeq2 models we used DESeq2::DESeq, for edgeR models we used *edgeR::glmQLFTest*. Covariates were selected from assay-specific technical metrics that explained variance in the data and were not correlated with exercise training: RNA integrity number (RIN), median 5’-3’ bias, percent of reads mapping to globin, and percent of PCR duplicates as quantified with Unique Molecular Identifiers (UMIs) for RNA-Seq; fraction of reads in peaks and library preparation batch for ATAC-seq.

### Genomic Distribution of Peaks and Methylation Sites

Gene and gene-region annotations for ATAC-seq peaks and methylation sites were generated through the *annotatePeak* function in the R package ChIPseeker^81^. Identified gene regions included proximal promoter (<= 1kb from TSS), distal promoter (1-2 kb from TSS), upstream (< 5kb), exon, intron, 5’ UTR, 3’ UTR, downstream (< 5kb), overlaps gene, and distal intergenic (outside the gene body or near vicinity). We then measured the frequency of peaks and methylation sites to be present in each gene region, as well as the frequency of each gene region annotation in DARs and DMRs. This was also conducted on the subsets of active peaks (we define peaks with a median normalized accessibility value > -1) and DEGaPs for each tissue. Significant changes in gene region distribution were identified by the *fisher.test* R function.

### Tissue enrichment of differential analytes

For each tissue and ome (RNA-seq, ATAC-seq, RRBS), the differential analytes were collected. For each differential analyte in a given ome, we measured the mean baseline (8 week control samples) expression. Z-scores were generated measuring the differences in baseline expression across all 8 tissues for each differential analyte. Differential analytes were ordered and annotated by their tissue of differential behavior.

### Identification of transcription factor motifs using HOMER

Transcription factor motif enrichment analysis was performed on sets of DARs, DMRs, and DEGaPs for each tissue. DARs and DMRs for motif enrichment analysis were selected for each tissue by satisfying an adjusted p-value threshold of 0.1. Similarly, DEGaPs for each tissue were selected by isolating the DEGs that satisfied an adjusted p-value threshold of 0.1, and selecting peaks annotated to the DEGs that contained a median normalized accessibility of -1. For genomic feature-specific analysis, DARs, DMRs, and DEGaPs were divided based upon their gene region annotation. The analysis was carried out by findMotifsGenome.pl (HOMER v4.11.1)^96^. It was performed on the ±50 bp flanking regions of the peak summits. The search lengths of the motifs were 8, 10, and 12 bp. We applied the -find flag to generate a list of all known rat motifs contained within the ±50 bp flanking regions of the summits for each peak in the ATAC-seq dataset and within the ±50 bp flanking regions of the center of each methylation site in the RRBS dataset, using the same settings as above. HOMER-identified TF motif enrichment for input lists of DARs, DMRs, and DEGaPs are calculated using a cumulative binomial distribution comparing TF motif representation among a target input list and a background list from which the target list is a subset.

### DAR genomic feature TF motif enrichment analysis

Applying the output from HOMER, the top ten significantly enriched TF motifs among DARs, DMRs and DEGaPs in each tissue were selected for further downstream analysis and cross-tissue comparisons. TFs were removed from further analysis if their gene was not expressed in the tissue in which their motifs were enriched. TF motif enrichments for differentially accessible regions (DARs) divided into gene features were calculated using the Fisher test. The test compared the ratio of DARs containing the motif for a specific TF/non-DARs containing this motif in one genomic feature, and the ratio of DARs containing this motif / non-DARs containing this motif in other genomic features. p-values were adjusted and FDR cutoff = 0.1 to select significant motifs in specific genomic features. This analysis was repeated for DMRs and DEGaPs. For the top TFs enriched among the DARs, DMRs, and DEGaPs of each tissue, we calculate their baseline expression (in control samples) and generated z scores of baseline expression across all tissues. Pearson correlation was calculated for TF enrichment z score versus TF baseline expression across all tissues.

### Correlations between DARs, DMRs, and DEGs

We selected DARs whose centers were within 500kb of a DEG TSS in each tissue. We then calculated the Pearson correlation of the L2FC of the DAR and the DEG for each EET sample relative to their sex control values using the *cor.test* function in R. We considered a DAR-DEG pair for further analysis if their training response Pearson correlation achieved a p-value less than 0.05 and the DAR contained a known motif for a TF expressed within the tissue. We identified TFs that were statistically enriched for containing binding sites within the DARs of DAR-DEG pairs using a binomial test comparing correlated targets:total correlated versus total targets: total active peaks in each tissue. We followed the same protocol to identify correlated DMR-DEG pairs and their associated enriched TFs.

### TFs significantly responding to EET and enriched for DEG targets

TFs were identified that showed significant response to EET at the RNA-seq, proteomic or phosphoproteomic level. Among TFs with significant EET responses, we calculated their enrichment for the promoter regions of DEGs. Enrichment significance was determined with a binomial test comparing the number of TF promoter targets in DEGs : total TF promoter targets versus the number of DEGs : total gene count in a given tissue.

### TF Enrichment among active promoter regions of DEGs with sex-consistent training responses

In each tissue, we identified subsets of DEGs that were up-regulated (L2FC > 0) in both sexes or down-regulated (L2FC < 0) in both sexes for each time point and then identified the DEGaPs associated with each set of up-regulated or down-regulated DEGs. We divided the DEGaPs by gene region, isolating the most predominant regions of distal intergenic, promoter (here a merge of the proximal and distal promoter assignments), and intron. For each TF expressed in a given tissue, we calculated the frequency of TF binding sites within each DEGaP population and then calculated z scores of binding site frequency across the DEGaP populations. As up-regulated and down-regulated promoter DEGaPs divided TF enrichment, we isolated the most enriched TFs for these two classes of DEGaPs, selecting the TFs most discerning between up-regulation and down-regulation within each tissue.

### Cell-type deconvolution

Cell type deconvolution was conducted by the R package CellCODE^97^ using the getallSPVs function. Marker sets were generated using the IRIS (Immune Response *In Silico*^98^ and DMAP (Differentiation Map) reference datasets^99^. The Kruskal-wallis test was implemented with the R function *kruskal.test* to determine if the variability in cell type proportion across samples in a given tissue would suggest a significant training response or sex difference.

### Pathway enrichment analyses

Gene set enrichment analysis (GSEA) was performed using ssGSEA2.0 with differential enrichment analysis t-scores (trained versus control) used as input. The gene set database used included canonical pathways available through the Human Matrisome database available through MatrisomeDB (http://matrisome.org/), and the MitoPathways database available through MitoCarta 3.082–84. Human gene symbols were mapped to rat orthologs before running the analysis. The ssGSEA2 function was run using parameters that avoid normalization, require at least 5 overlapping features with the gene set, and use the area under the curve as the enrichment metric (sample.norm.type = “none”, weight=0.75, correl.type = “rank”, statistic =“area.under.RES”, output.score.type = “NES”, min.overlap=5). (cite the landscape paper )

Overrepresentation analysis of significant DEGs was performed for each tissue separately using the R package gprofiler2:gost using Gene Ontology Biological Process, Reactome, WikiPathways and KEGG databases against background gene sets defined as a set of all genes expressed in each tissue, taken from the rna-seq data^100^. Top 10 non-redundant pathway enrichments based on Benjamini-Hochberg (BH) corrected false discovery rate for each tissue are displayed as bubble plots with sizes indicating the number of significant genes relative to the pathway size (number of genes in that pathway) and colors indicating the significance (BH-corrected p-value). At least 10 genes were required to be differential in a pathway with a maximum of 200 genes.

### Correlations between Phenotypic Measures and DEGs

Phenotypic measures were calculated at weeks 4 and 8 of EET and in week 8 controls. Measures were presented as changes between time point and original baseline measurements in each rat. For each phenotypic measure and DEG combination, we calculated the Pearson correlation between the change in phenotypic measure between baseline and a given time point, and the gene expression of the DEG at the time point for each animal subject. We isolated the DEGs that exhibited > 0.5 or < -0.5 correlations with each phenotypic measure in each tissue and selected the DEGaPs annotated to the promoter region of the DEGs. TF motif enrichment significance among a set of positively or negatively correlated DEG’s promoter DEGaPs in a tissue were determined by an exact binomial test comparing the frequency of enrichment among phenotype-correlated DEGs versus general enrichment among the promoter DEGaPs in the tissue.

## Data Availability

MoTrPAC data are publicly available via motrpac-data.org/data-access. Data access inquiries should be sent to. Additional resources can be found at motrpac.org and motrpac-data.org. Raw and processed data were deposited in the following public repositories: RNA-seq, ATAC-seq and RRBS data were deposited at the Sequence Read Archive under accession PRJNA908279 and at the Gene Expression Omnibus under accession GSE242358; proteomics data were deposited at MassIVE under accessions MSV000092911, MSV000092922, MSV000092923, MSV000092924, MSV0000929 25 and MSV000092931.

## Code Availability

MoTrPAC data processing pipelines for RNA-Seq, ATAC-seq, RRBS, and proteomics will be made public at the time of publication: https://github.com/MoTrPAC/motrpac-rna-seq-pipeline, https://github.com/MoTrPAC/motrpac-atac-seq-pipeline, https://github.com/MoTrPAC/motrpacrrbs-pipeline, https://github.com/MoTrPAC/motrpac-proteomics-pipeline. Normalization and QC scripts will be available at https://github.com/MoTrPAC/motrpac-bic-norm-qc. Code for the underlying differential analysis for the manuscript will be provided in the MotrpacRatTraining6mo R package (motrpac.github.io/MotrpacRatTraining6mo). Code for conducting the analysis and generating the figures contained within this paper will be available at https://github.com/gsmith990306/MoTrPAC_PASS1B_Transcription_Factor_Paper.

## Inclusion and Ethics Statement

Research responsibilities and authorship were agreed amongst the collaborators during the research and writing processes. As this was a rat-based study of exercise, there were no international local collaborators to include in this study; however, we endeavored to include all participating researchers in the author list, either individually named or as part of the MoTrPAC consortium. Animal welfare regulations were strictly followed and animal procedures were approved by the institutional animal care and use committee at the University of Iowa.

## Acknowledgements

**Funding:** The MoTrPAC Study is supported by NIH grants U24OD026629 (Bioinformatics Center), U24DK112349, U24DK112342, U24DK112340, U24DK112341, U24DK112326, U24DK112331, U24DK112348 (Chemical Analysis Sites), U01AR071133, U01AR071130, U01AR071124–01, U01AR071128, U01AR071150, U01AR071160, U01AR071158 (Clinical Centers), U24AR071113 (Consortium Coordinating Center), U01AG055133, U01AG055137 and U01AG055135 (PASS/Animal Sites).

Parts of this work were performed in the Environmental Molecular Science Laboratory, a U.S. Department of Energy national scientific user facility at Pacific Northwest National Laboratory in Richland, Washington.

## Author Contributions

Gregory R. Smith, supervised the investigation, performed data and bioinformatics analysis, prepared figures, drafted and revised the original manuscript

Bingqing Zhao, supervised the investigation, performed data and bioinformatics analysis, prepared figures, drafted and revised the original manuscript

Malene E. Lindholm, performed data and bioinformatics analysis, prepared figures, drafted and revised the original manuscript

Archana Raja, performed data and bioinformatics analysis, prepared figures, revised the original manuscript

Mark Viggars, contributed to data interpretation, drafted and revised the original manuscript Hanna Pincas, contributed to data interpretation, revised the original manuscript

Nicole R. Gay, performed data and bioinformatics analysis, revised the original manuscript Yifei Sun, performed and analyzed experiments

Sindhu Vangeti, contributed to data interpretation Yongchao Ge, performed data and bioinformatics analysis

Venugopalan D. Nair, supervised data generation, contributed to data interpretation James A. Sanford, contributed to data interpretation, revised the original manuscript

Mary Anne S. Amper, performed and analyzed experiments Mital Vasoya, performed and analyzed experiments

Kevin S. Smith, contributed to data interpretation Irene Ramos, contributed to data interpretation

Stephen Montgomery, contributed to data interpretation, revised the original manuscript Elena Zaslavsky, contributed to data interpretation

Sue C. Bodine, contributed to data interpretation

Karyn A. Esser, contributed to data interpretation, drafted and revised the original manuscript Martin J. Walsh, contributed to data interpretation, revised the original manuscript

Michael P. Snyder, provided overall leadership and guidance to the investigation Stuart C. Sealfon, provided overall leadership and guidance to the investigation

## Abbreviations

Abbreviation Definition

BAT: Brown adipose tissue
DAR: Differentially accessible regions
DEG: Differentially expressed genes
DEGaP: Differentially expressed gene associated peak
EET: Endurance Exercise Training
FAP: cell Fibro/adipogenic progenitor cell
GMP: cell Granulocyte/monocyte progenitor cell
HEART: Heart
HIPPOC: Hippocampus
KIDNEY: Kidney
LIVER: Liver
LUNG: Lung
NK: cell Natural Killer cell
SKM-GN: Skeletal muscle (Gastrocnemius)
TES: Transcription end site
TF: Transcription factor
TSS: Transcription start site
WAT-SC: Subcutaneous white adipose tissue

## Supplementary Information

Supplementary Figures. Supplementary Figures S1-S38 and corresponding legends.

Supplementary Table S1. Pearson correlations of DEGs in each tissue with estimated immune cell type composition changes. Each row is a DEG in a tissue and each column an immune cell type included in the deconvolution analysis.

Supplementary Table S2. DAR-DEG pairs with Pearson correlation > 0.5 or < -0.5 and annotated with an expressed TF motif. For each pair, we listed the tissue, DEG ensembl id and symbol, DAR id, distance between DAR midpoint and DEG TSS, Pearson correlation, known TF motifs within the DAR, and the L2FC of the DEG and the DAR for each time point relative to eight week control in both female and male samples. Brown adipose DAR-DEG pairs are excluded because of confounding cell type composition effects.

Supplementary Table S3. DMR-DEG pairs with Pearson correlation > 0.5 or < -0.5 and annotated with an expressed TF motif. For each pair, we listed the tissue, DEG ensembl id and symbol, DMR id, distance between DMR midpoint and DEG TSS, Pearson correlation, known TF motifs within the DMR, and the L2FC of the DEG and the DMR for each time point relative to eight week control in both female and male samples. Brown adipose DMR-DEG pairs are excluded because of confounding cell type composition effects.

Supplementary Table S4. Time of day for daily training for each animal subject. Rows are labeled by a unique id representing each animal and columns represent each training day.

## Supplemental Figures

Supplemental Figure S1: Overlap in DEGs, DARs, and DMRs across tissues. (a) Proportion of intersection of DEG lists divided by the union of DEG lists for each pair of tissues. Tissues clustered by overlap. (b) Proportion of intersection of DAR lists divided by the union of DAR lists for each pair of tissues. Tissues clustered by overlap. (c) Proportion of intersection of DMR lists divided by the union of DMR lists for each pair of tissues. Tissues clustered by overlap.

Supplemental Figure S2: Distribution of training responses for DEGs in eight tissues. (a-h) A histogram of L2FC values across sexes and time points for all DEGs, a separate histogram of L2FC values across sexes and time points for all genes, and a heatmap of L2FC values for DEGs ordered by hierarchical clustering for skeletal muscle (a), heart (b), hippocampus (c), kidney (d), liver (e), lung (f), brown adipose (e), and white adipose (f).

Supplemental Figure S3: Distribution of training responses for DARs in eight tissues. (a-h) A histogram of L2FC values across sexes and time points for all DARs, a separate histogram of L2FC values across sexes and time points for all peaks, and a heatmap of L2FC values for DARs ordered by hierarchical clustering for skeletal muscle (a), heart (b), hippocampus (c), kidney (d), liver (e), lung (f), brown adipose (e), and white adipose (f).

Supplemental Figure S4: Distribution of training responses for DMRs in eight tissues. (a-h) A histogram of L2FC values across sexes and time points for all DMRs, a separate histogram of L2FC values across sexes and time points for all methylation regions, and a heatmap of L2FC values for DMRs ordered by hierarchical clustering for skeletal muscle (a), heart (b), hippocampus (c), kidney (d), liver (e), lung (f), brown adipose (e), and white adipose (f).

Supplemental Figure S5: Multiomic power analysis in eight tissues for RNAseq, ATACseq, RRBS, protein abundance (Prot_PR) and protein phosphorylation (Prot_PH) data. For each tissue, plots of statistical power for each ome at different sample sizes (a,c,e,g,i,k,m,o) and with different cutoffs for effect size (b,d,f,h,j,l,n,p). The square represents the sample size or cutoff needed to achieve 0.25 power in all omes. Tissues include skeletal muscle (SKM-GN) (a,b), heart (c,d), hippocampus (HIPPOC) (e,f), kidney (g,h), liver (i,j), lung (k,l), brown adipose (BAT) (m,n), white adipose (WAT-SC) (o,p).

Supplemental Figure S6: PCA plots of top two PCs for RNA-seq data in eight tissues: skeletal muscle (a), heart (b), hippocampus (c), kidney (d), liver (e), lung (f), brown adipose (e), and white adipose (f).

Supplemental Figure S7: PCA plots of top two PCs for ATAC-seq data in eight tissues: skeletal muscle (a), heart (b), hippocampus (c), kidney (d), liver (e), lung (f), brown adipose (e), and white adipose (f).

Supplemental Figure S8: PCA plots of top two PCs for RRBS data in eight tissues: skeletal muscle (a), heart (b), hippocampus (c), kidney (d), liver (e), lung (f), brown adipose (e), and white adipose (f).

Supplemental Figure S9: Variability in GSEA enrichment across tissue, sex and time point. Gene set enrichment analysis (GSEA)-generated normalized enrichment score (NES) for significantly enriched pathways among week 1 and week 8 training response genes in both sexes and eight tissues.

Supplemental Figure S10: Cell type deconvolution of RNA-seq data for eight tissues. (a-h) z-scores of predicted shifts in cell type proportion for each sample in brown adipose (a), white adipose (b), skeletal muscle (c), heart (d), hippocampus (e), kidney (f), liver (g), and lung (h).

Supplemental Figure S11: Heatmaps of correlation between the EET response of DEGs and cell type composition changes in eight tissues: BAT (a), WAT-SC (b), SKM-GN (c), heart (d), HIPPOC (e), kidney (f), liver (g), lung (h).

Supplemental Figure S12: Comparison of accessible peak distribution between expressed genes and DEGs. (A) Distribution of genomic regions among the accessible peaks for expressed genes in each tissue. (B) Distribution of genomic regions among the accessible peaks for DEGs (DEGaPs) in each tissue. (c) DEGaPs are enriched in exons for a majority of tissues and exhibit a decreased proportion in distal intergenic regions for half of tissues.

Supplemental Figure S13: (a-c) Heatmaps of all up-regulated differential analytes across eight tissues in RNA-seq (a), ATAC-seq (b), and RRBS (c). Columns reflect z scores of baseline expression (a), accessibility (b), or methylation (c), across the eight tissues. Rows are annotated by the tissues in which each analyte exhibits a differential training response. (d-f) Heatmaps of all down-regulated differential analytes across eight tissues in RNA-seq (d), ATAC-seq (e), and RRBS (f). Columns reflect z scores of baseline expression (d), accessibility (e), or methylation (f), across the eight tissues. Rows are annotated by the tissues in which each analyte exhibits a differential training response.

Supplemental Figure S14: Density scatter plots of DAR-DEG distance and training response correlation for eight tissues: skeletal muscle (a), heart (b), hippocampus (c), kidney (d), liver (e), lung (f), brown adipose (e), and white adipose (f).

Supplemental Figure S15: Density scatter plots of DMR-DEG distance and training response correlation for eight tissues: skeletal muscle (a), heart (b), hippocampus (c), kidney (d), liver (e), lung (f), brown adipose (e), and white adipose (f).

Supplemental Figure S16: Density scatter plots of DMR-DAR distance and training response correlation for six tissues with non-zero measurements of neighboring DMR-DAR pairs: skeletal muscle (a), heart (b), kidney (c), liver (d), lung (e), and brown adipose (f).

Supplemental Figure S17: Per tissue pathway enrichment analysis of DEGs withDARs located within 500 kb. Top 10 enriched, non-redundant pathways are shown for each tissue.

Supplemental Figure S18: (a-l) Training response L2FC scatter plots of correlated DAR-DEG examples. (a-b) SP2-target DARs in lung correlated with *Dennd1c* (a), *Oas2* (b); (c-f) BMYB- target DARs in SKM-GN correlated with *Cited4* (c), *Cab39* (d), *Sik1* (e), *Phlda3* (f); (g-l) BMAL1- target DARs in liver correlated with *G6pc1* (g-i), *Chp1* (j), *Onecut1* (k), *Lpar3* (l). (m-n) DAR-DEG pairs containing the binding motif of MAZ in SKM-GN (m), and lung (n). (o-q) MAZ-target DARs in SKM-GN correlated with *Kcna7* (o), *Eif4g3* (p), *Srsf2* (q); (r) MAZ-target DAR in lung correlated with *Hspb6*.

Supplemental Figure S19: (a-b) TF NF1-halfsite correlated DMR-DEG targets in WAT-SC (a), and lung (b). (c) TF AP-2gamma correlated DMR-DEG targets in lung. (d-p) Training response L2FC scatter plots of correlated DMR-DEG examples. (d) SP2-target DMR in SKM-GN correlated with *Zdhhc1*. (e-j) NF1-halfsite-target DMRs in WAT-SC correlated with *Erg* (e), *Kcnj15* (f), *Syt11* (g), *Crabp2* (h), *Paqr6* (i), *Tmem79* (j); (k-n) NF1-halfsite and AP-2gamma targeted DMRs in lung correlated with *Agap2* (k), *B4galnt1* (l), *Arhgap9* (m), *Avil* (n); (o-p) AP- 2gamma-target DMRs in lung correlated with *Tmem176a* (o), *Tmem176b* (p).

Supplemental Figure S20: Identifying cross-tissue conserved differential signal in TFs. (a-c) Heatmaps of TFs displaying significance at the RNA-seq (a), proteome (b), and phosphoproteome (c) level. In each heatmap, a grid square is red if the TF is significant in that ome in that tissue. (d-f) Quantifying the overlap of significant TFs at the RNA-seq (d), proteome (e), and phosphoproteome (f) level.

Supplemental Figure S21: Scatter plots of magnitude (x-axis) and significance (y-axis) of correlation in EET response for TFs across omes and tissues. Correlation is measured for EET response between protein abundance and protein phosphorylation, gene expression and protein phosphorylation and gene expression and protein abundance for each tissue: skeletal muscle (a), heart (b), kidney (c), liver (d), lung (e), white adipose (f). Names of TFs on the scatter plots are colored orange if the TF has a significant (adjusted p.val < 0.1) EET response in one of the two compared omes, and colored red if the TF has a significant EET response in both compared omes.

Supplemental Figure S22: Frequency of DEGs among gene targets of TFs with significant proteomic or phosphoproteomic training response in (a) skeletal muscle, (b) heart, (c) kidney, (d) liver, (e) lung, (f) white adipose. TFs are color-annotated by the ome of their training response: protein (red), phosphoprotein (blue) or both (green). The dashed line represents the frequency of DEGs among active genes in each tissue. Gene targets are divided by presence of TF motif within active peaks annotated to the proximal promoter (orange), intron (brown) or distal intergenic (gold) features.

Supplemental Figure S23: Frequency of DEGs among gene targets of TFs with significant transcriptomic training response in (a) skeletal muscle, (b) heart, (c) hippocampus, (d) kidney, (e) liver, (f) lung, (g) brown adipose, and (f) white adipose. The dashed line represents the frequency of DEGs among active genes in each tissue. Gene targets are divided by presence of TF motif within active peaks annotated to the proximal promoter (orange), intron (brown) or distal intergenic (gold) features.

Supplemental Figure S24: Heatmaps of EET response for DEGs witih proximal promoter motifs of TFs with significant changes at the RNA-seq level including (a) SF1, (b) Six1 in SKM-GN, (c) Fos in kidney, (d) JunD in SKM-GN, (e) JunD in heart, (f) JunD in WAT-SC, (g) PBX1 in lung, (h) RAR:RXR in BAT.

Supplemental Figure S25: Heatmaps of EET response for DEGs with proximal promoter motifs of TFs with significant changes at the proteomic (a-d) or phosphoproteomic (e-h) level including (a) MEF2C in SKM-GN, (b) IRF:BATF in lung, (c) NR4A1 in SKM-GN, (d) PBX1 in WAT-SC among proteomic TF changes, and (e) MEF2C in SKM-GN, (f) MEF2A in heart, (g) ATF7 in kidney, and (h) STAT1 in liver among phosphoproteomic TF changes.

Supplemental Figure S26: Dotplots measuring correlation between TF response to EET and proximal promoter targets’ response to EET for select TFs with significant training responses at the RNA, proteome or phosphoproteome levels in skeletal muscle (a), heart (b), kidney (c), liver (d), lung (e), white adipose (f). Target genes are divided into non-DEGs (red) and DEGs (blue). * represents p < 0.05 of t test for difference between correlation of TF with DEG targets and non-DEG targets.

Supplemental Figure S27: (a) z-scores of TF motif frequency among DARs across tissues for top DAR-enriched TFs. (b) z-scores of TF motif frequency among DMRs across tissues for top DMR-enriched TFs. (c) z-scores of TF motif frequency among DEGaPs across tissues for top DEGaP-enriched TFs.

Supplemental Figure S28: Comparing TF enrichment among DARs and DEGaPs. Scatter plots of -log10 p-value of TF motif enrichment among DARs and DEGaPs in six tissues: (a) skeletal muscle, (b) heart, (c) kidney, (d) liver, (e) lung, (f) brown adipose. Pearson correlation is calculated for each tissue.

Supplemental Figure S29: Comparing TF enrichment among DMRs and DEGaPs. Scatter plots of -log10 p-value of TF motif enrichment among DARs and DEGaPs in six tissues: (a) skeletal muscle, (b) heart, (c) hippocampus, (d) kidney, (e) liver, (f) lung, (g) brown adipose, and (h) white adipose. Pearson correlation is calculated for each tissue.

Supplemental Figure S30: Comparing TF enrichment among DARs and DMRs. Scatter plots of - log10 p-value of TF motif enrichment among DARs and DEGaPs in six tissues: (a) skeletal muscle, (b) heart, (c) kidney, (d) liver, (e) lung, (f) brown adipose. Pearson correlation is calculated for each tissue.

Supplemental Figure S31: (a) z-scores of TF gene expression in control samples across tissues for top DAR-enriched TFs. (b) z-scores of TF gene expression in control samples across tissues for top DMR-enriched TFs. (c) z-scores of TF gene expression in control samples across tissues for top DEGaP-enriched TFs.

Supplemental Figure S32: (a) Training response L2FC across sexes, time points and tissues for top DAR-enriched TFs. (b) Training response L2FC across sexes, time points and tissues for top DMR-enriched TFs. (c) Training response L2FC across sexes, time points and tissues for top DEGaP-enriched TFs.

Supplemental Figure S33: (a) Correlation, for each tissue, of z-scores of TF motif frequency among DARs across tissues (S27a) and z-scores of TF gene expression in control samples across tissues (S28a). (b,c) Scatter plots of enrichment z score vs expression z score for top DAR-enriched TFs in heart (b) and lung (c). (d) Correlation, for each tissue, of z-scores of TF motif frequency among DMRs across tissues (S27b) and z-scores of TF gene expression in control samples across tissues (S28b). (e,f) Scatter plots of enrichment z score vs expression z score for top DAR-enriched TFs in hippocampus (e) and brown adipose (f). (g) Correlation, for each tissue, of z-scores of TF motif frequency among DEGaPs across tissues (S27c) and z-scores of TF gene expression in control samples across tissues (S28c). (h,i) Scatter plots of enrichment z score vs expression z score for top DEGaP-enriched TFs in liver (h) and lung (i). Pearson correlation is calculated for each tissue and a linear model is fit to the data. Shaded regions in the scatter plots in panels b,c,e,f,h,i reflected a 95% confidence interval for predictions from a linear model.

Supplemental Figure S34: z-scores of TF motif frequency among DEGaPs divided into up- regulated and down-regulated genes at each training time point and split by genomic feature in: (a) skeletal muscle, (b) heart, (c) hippocampus, (d) kidney, (e) liver, (f) lung, (g) brown adipose, (h) white adipose. For comparison, each panel includes DEGaP sets not divided by genomic feature or training response. Distinct patterns of TF motif frequency among the up-regulated and down-regulated DEG promoter peaks.

Supplemental Figure S35: Conserved TF motif frequency of up-regulated and down-regulated promoter peaks in each tissue. z-scores of TF motif frequency among DEG promoter peaks divided by up-regulated and down-regulated training responses at each training time point in: (a) skeletal muscle, (b) heart, (c) hippocampus, (d) kidney, (e) liver, (f) lung, (g) brown adipose, (h) white adipose. TFs were selected in each tissue for enrichment in up-regulated or down- regulated DEG promoter peaks and differentiation of enrichment between up-regulated and down-regulated DEG promoter peaks.

Supplemental Figure S36: Boxplots of phenotypic changes over training for body weight (a), body fat (b), vo2 max (c), body lean (d), lactate (e), and body water(f). Statistical measurements were generated with a t test. ns (not significant) represents *p* >= 0.05, \**p <* 0.05, \*\**p <* 0.01, \*\*\**p <* 0.001, \*\*\*\**p <* 0.0001. In each boxplot generated by the function *ggboxplot*, the center line, represents median, the bottom and top of the box are the first and third quartile with whiskers extending to the smallest and largest non-outliers in the data measurement. Outliers are defined as at least 1.5 times outside the interquartile range.

Supplemental Figure S37: Heatmaps of correlation between DEGs and phenotypic measures for eight tissues: skeletal muscle (a), heart (b), hippocampus (c), kidney (d), liver (e), lung (f), brown adipose (e), and white adipose (f). Most tissues show distinct groups of DEGs positively correlated with vo2 max and body lean vs body fat and weight gain, with the exception of brown adipose (DEGs driven by cell type changes).

Supplemental Figure S38: Significant correlations of TF training response with phenotypic measures in RNA-seq (a) and proteomic (b) data. All TFs listed have at least one phenotypic correlation > 0.5 or < -0.5 in the annotated tissue (annotated at the left of each row).

## References

1. Neufer, P. D. et al. Understanding the Cellular and Molecular Mechanisms of Physical Activity- Induced Health Benefits. Cell Metab. 22, 4–11 (2015).

2. Mansueto, G. et al. Transcription Factor EB Controls Metabolic Flexibility during Exercise. Cell Metab. 25, 182–196 (2017).

3. Bléher, M. et al. Egr1 loss-of-function promotes beige adipocyte differentiation and activation specifically in inguinal subcutaneous white adipose tissue. Sci. Rep. 10, 15842 (2020).

4. Popov, D. V. et al. Contractile activity-specific transcriptome response to acute endurance exercise and training in human skeletal muscle. Am. J. Physiol.-Endocrinol. Metab. 316, E605– E614 (2019).

5. MoTrPAC Study Group et al. Temporal dynamics of the multi-omic response to endurance exercise training across tissues. 2022.09.21.508770 Preprint at 10.1101/2022.09.21.508770 (2022).

6. Nair, V. D., et al. Molecular adaptations in response to exercise training are associated with tissue-specific transcriptomic and epigenomic signatures. Cell Genom. 2024 Jun 12;4(6):100421. doi: 10.1016/j.xgen.2023.100421.

7. Oliveira, A. N. & Hood, D. A. Exercise is mitochondrial medicine for muscle. Sports Med. Health Sci. 1, 11–18 (2019).

8. Gregoire, F. M., Smas, C. M. & Sul, H. S. Understanding Adipocyte Differentiation. Physiol. Rev. 78, 783–809 (1998).

9. Moore, D. & Loprinzi, P. D. Exercise influences episodic memory via changes in hippocampal neurocircuitry and long-term potentiation. Eur. J. Neurosci. 54, 6960–6971 (2021).

10. Sonawane, A. R. et al. Understanding Tissue-Specific Gene Regulation. Cell Rep. 21, 1077– 1088 (2017).

11. Ramachandran, K. et al. Dynamic enhancers control skeletal muscle identity and reprogramming. PLOS Biol. 17, e3000467 (2019).

12. Miao, Z. et al. Single cell regulatory landscape of the mouse kidney highlights cellular differentiation programs and disease targets. Nat. Commun. 12, 2277 (2021).

13. Yan, F., Powell, D. R., Curtis, D. J. & Wong, N. C. From reads to insight: a hitchhiker’s guide to ATAC-seq data analysis. Genome Biol. 21, 22 (2020).

14. Izzo, M. W., Strachan, G. D., Stubbs, M. C. & Hall, D. J. Transcriptional Repression from the c- myc P2 Promoter by the Zinc Finger Protein ZF87/MAZ *. J. Biol. Chem. 274, 19498–19506 (1999).

15. Nair, V. D. et al. Differential analysis of chromatin accessibility and gene expression profiles identifies cis-regulatory elements in rat adipose and muscle. Genomics 113, 3827–3841 (2021).

16. Panigrahi, A. & O’Malley, B. W. Mechanisms of enhancer action: the known and the unknown. Genome Biol. 22, 108 (2021).

17. Maity, G. et al. The MAZ transcription factor is a downstream target of the oncoprotein Cyr61/CCN1 and promotes pancreatic cancer cell invasion via CRAF–ERK signaling. J. Biol. Chem. 293, 4334–4349 (2018).

18. Bang, P. et al. Exercise-induced changes in insulin-like growth factors and their low molecular weight binding protein in healthy subjects and patients with growth hormone deficiency. Eur. J. Clin. Invest. 20, 285–292 (1990).

19. Stein, A. M. et al. Physical exercise, IGF-1 and cognition A systematic review of experimental studies in the elderly. Dement. Neuropsychol. 12, 114–122 (2018).

20. Jiao, S. et al. Differential regulation of IGF-I and IGF-II gene expression in skeletal muscle cells. Mol. Cell. Biochem. 373, 107–113 (2013).

21. Clavarino, G. et al. Protein phosphatase 1 subunit Ppp1r15a/GADD34 regulates cytokine production in polyinosinic:polycytidylic acid-stimulated dendritic cells. Proc. Natl. Acad. Sci. U. S. A. 109, 3006–3011 (2012).

22. Farkas, C. et al. Characterization of SALL2 Gene Isoforms and Targets Across Cell Types Reveals Highly Conserved Networks. Front. Genet. 12, 613808 (2021).

23. Amar, D. et al. Time trajectories in the transcriptomic response to exercise - a meta-analysis. Nat. Commun. 12, 3471 (2021).

24. García-Pelagio, K. P., Bloch, R. J., Ortega, A. & González-Serratos, H. Biomechanics of the sarcolemma and costameres in single skeletal muscle fibers from normal and dystrophin-null mice. J. Muscle Res. Cell Motil. 31, 323–336 (2011).

25. Satpathy, A.T., et al. Runx1 and Cbfβ regulate the development of Flt3+ dendritic cell progenitors and restrict myeloproliferative disorder. Blood. 123(**19**), 2968–2977 (2014).

26. Huang, G., et al. PU.1 is a major downstream target of AML1 (RUNX1) in adult mouse hematopoiesis. Nat. Genet. 40, 51–60 (2008).

27. Himeda, C. L., Ranish, J. A., Pearson, R. C. M., Crossley, M. & Hauschka, S. D. KLF3 regulates muscle-specific gene expression and synergizes with serum response factor on KLF binding sites. Mol. Cell. Biol. 30, 3430–3443 (2010).

28. Taylor, M. V. & Hughes, S. M. Mef2 and the skeletal muscle differentiation program. Semin. Cell Dev. Biol. 72, 33–44 (2017).

29. Gallant, S. & Gilkeson, G. ETS transcription factors and regulation of immunity. Arch. Immunol. Ther. Exp. (Warsz.) 54, 149–163 (2006).

30. Seifert, L. L. et al. The ETS transcription factor ELF1 regulates a broadly antiviral program distinct from the type I interferon response. PLoS Pathog. 15, e1007634 (2019).

31. Bauer, F., et al. Specificity protein 2 (Sp2) is essential for mouse development and autonomous proliferation of mouse embryonic fibroblasts. PLoS ONE, 5(**3**), e9587 (2010).

32. Terrados, G., et al. Genome-wide localization and expression profiling establish Sp2 as sequence-specific transcription factor regulating vitally important genes. Nuc. Acids Res., 40(**16**), 7844–7857 (2012).

33. Bayly-Jones, C., Pang, S. S., Spicer, B. A., Whisstock, J. C. & Dunstone, M. A. Ancient but Not Forgotten: New Insights Into MPEG1, a Macrophage Perforin-Like Immune Effector. Front. Immunol. 11, (2020).

34. Liao, X. et al. 2′, 5′-Oligoadenylate Synthetase 2 (OAS2) Inhibits Zika Virus Replication through Activation of Type Ι IFN Signaling Pathway. Viruses 12, 418 (2020).

35. Zhou, D., et al. Editing of CD1d-bound lipid antigens by endosomal lipid transfer proteins. Science 303(5657), 523–527 (2003).

36. Ha, J., et al. Identification of a novel PML-RARG fusion in acute promyelocytic leukemia. Leukemia 31, 1992–1995 (2017).

37. Tang, J., et al. Cancer cells escape p53’s tumor suppression through ablation of ZDHHC1- mediated p53 palmitoylation. Oncogene 40, 5416–5426 (2021).

38. Le, X., et al. DNA methylation downregulated ZDHHC1 suppresses tumor growth by altering cellular metabolism and inducing oxidative/ER stress-mediated apoptosis and pyroptosis. Theranostics 10(**21**), 9495–9511 (2020).

39. Induction of GADD34 Is Necessary for dsRNA-Dependent Interferon-β Production and Participates in the Control of Chikungunya Virus Infection | PLOS Pathogens. https://journals.plos.org/plospathogens/article?id=10.1371/journal.ppat.1002708.

40. Li, F., Xiao, H., Zhou, F., Hu, Z. & Yang, B. Study of HSPB6: Insights into the Properties of the Multifunctional Protective Agent. Cell. Physiol. Biochem. 44, 314–332 (2017).

41. Rogeri, P. S. et al. Crosstalk Between Skeletal Muscle and Immune System: Which Roles Do IL-6 and Glutamine Play? Front. Physiol. 11, (2020).

42. Ziebold, U., et al. Phosphorylation and activation of B-Myb by cyclin A-Cdk2. Curr. Biol. 7(**4**), 253–260 (1997).

43. Yalcin, A., et al. Nuclear targeting of 6-phosphofructo-2-kinase (PFKFB3) increases proliferation via cyclin-dependent kinases. J. Biol. Chem. 284(**36**), 24223–24232 (2009).

44. Boyd, S., et al. Structure-based design of potent and selective inhibitors of the metabolic kinase PFKFB3. J. Med. Chem. 58(**8**), 3611–3625 (2015).

45. Jagganath, A., et al. The CRTC1-SIK1 pathway regulates entrainment of the circadian clock. Cell 154(**5**), 1100–1111 (2013).

46. Moreno-Risco, M., et al. Juvenile hemochromatosis due to a homozygous variant in the HJV gene. Case Rep. Pediatr. 2022, 7743748 (2022).

47. Peng, L., Yan, H., Qi, S., and Deng, L. CAB39 promotes the proliferation of nasopharyngeal carcinoma CNE-1 cells via up-regulating p-JNK. Cancer Manag. Res. 12, 11203–11209 (2020).

48. Lerchenmuller, C., et al. CITED4 protects against adverse remodeling in response to physiological and pathological stress. Circ. Res. 127(**5**), 631–646 (2020).

49. Auman, H. J., et al. Transcription factor AP-2γ is essential in the extra-embryonic lineages for early postimplantation development. Development. 129, 2733–2747 (2002).

50. Zhang, H., et al. TFAP2C exacerbates psoriasis-like inflammation by promoting Th17 and Th1 cells activation through regulating TEAD4 transcription. Allergol. Immunopathol. 51, 124–134 (2023).

51. Trovó-Marqui, A. B., et al. Neurofibromin: a general outlook. Clin. Genet. 70, 1–13 (2006).

52. Ferreira de Souza, J., et al. Exercise capacity impairment in individuals with neurofibromatosis type 1. Am. J. Med. Genet. Part A. **161A**, 393–395 (2013).

53. Fernández-Verdejo, R. et al. Activating transcription factor 3 attenuates chemokine and cytokine expression in mouse skeletal muscle after exercise and facilitates molecular adaptation to endurance training. FASEB J. Off. Publ. Fed. Am. Soc. Exp. Biol. 31, 840–851 (2017).

54. Kawasaki, E. et al. Role of local muscle contractile activity in the exercise-induced increase in NR4A receptor mRNA expression. J. Appl. Physiol. Bethesda Md 1985 106, 1826–1831 (2009).

55. Rundqvist, H. C. et al. Acute sprint exercise transcriptome in human skeletal muscle. PLOS ONE 14, e0223024 (2019).

56. Puntschart, A. et al. Expression of fos andjun genes in human skeletal muscle after exercise. Am. J. Physiol.-Cell Physiol. 274, C129–C137 (1998).

57. Ku, H.-C. & Cheng, C.-F. Master Regulator Activating Transcription Factor 3 (ATF3) in Metabolic Homeostasis and Cancer. Front. Endocrinol. 11, (2020).

58. Zhang, L. et al. The Orphan Nuclear Receptor 4A1: A Potential New Therapeutic Target for Metabolic Diseases. J. Diabetes Res. 2018, 9363461 (2018).

59. Pei, L., Castrillo, A. & Tontonoz, P. Regulation of Macrophage Inflammatory Gene Expression by the Orphan Nuclear Receptor Nur77. Mol. Endocrinol. 20, 786–794 (2006).

60. Liebmann, M., et al. Nur77 serves as a molecular brake of the metabolic switch during T cell activation to restrict autoimmunity. Proc. Natl. Acad. Sci. 115, E8017–E8026 (2018).

61. Mayer, M. P. & Bukau, B. Hsp70 chaperones: Cellular functions and molecular mechanism. Cell. Mol. Life Sci. 62, 670–684 (2005).

62. Cooper, L. M., West, R. C., Hayes, C. S. & Waddell, D. S. Dual-specificity phosphatase 29 is induced during neurogenic skeletal muscle atrophy and attenuates glucocorticoid receptor activity in muscle cell culture. Am. J. Physiol.-Cell Physiol. 319, C441–C454 (2020).

63. Zhao, X. et al. Nuclear receptors rock around the clock. EMBO Rep. 15, 518–528 (2014).

64. Oiwa, A. et al. Synergistic regulation of the mouse orphan nuclear receptor SHP gene promoter by CLOCK–BMAL1 and LRH-1. Biochem. Biophys. Res. Commun. 353, 895–901 (2007).

65. Rey, G. et al. Genome-Wide and Phase-Specific DNA-Binding Rhythms of BMAL1 Control Circadian Output Functions in Mouse Liver. PLOS Biol. 9, e1000595 (2011).

66. Sato, S. et al. Atlas of exercise metabolism reveals time-dependent signatures of metabolic homeostasis. Cell Metab. 34, 329–345.e8 (2022).

67. Kemler, D., Wolff, C. A. & Esser, K. A. Time of Day Dependent Effects of Contractile Activity on the Phase of the Skeletal Muscle Clock. 2020.03.05.978759 Preprint at 10.1101/2020.03.05.978759 (2020).

68. Casanova-Vallve, N. et al. Daily running enhances molecular and physiological circadian rhythms in skeletal muscle. Mol. Metab. 61, 101504 (2022).

69. Silva, B. S. de A., et al. Exercise as a Peripheral Circadian Clock Resynchronizer in Vascular and Skeletal Muscle Aging. Int. J. Environ. Res. Public. Health 18, 12949 (2021).

70. Mansingh, S. & Handschin, C. Time to Train: The Involvement of the Molecular Clock in Exercise Adaptation of Skeletal Muscle. Front. Physiol. 13, (2022).

71. Duglan, D. & Lamia, K. A. Clocking In, Working Out: Circadian Regulation of Exercise Physiology. Trends Endocrinol. Metab. 30, 347–356 (2019).

72. Wisløff, U., Helgerud, J., Kemi, O. J. & Ellingsen, O. Intensity-controlled treadmill running in rats: VO(2 max) and cardiac hypertrophy. Am. J. Physiol. Heart Circ. Physiol. 280, H1301–1310 (2001).

73. Corces, M. R. et al. An improved ATAC-seq protocol reduces background and enables interrogation of frozen tissues. Nat. Methods 14, 959–962 (2017).

74. Moore, J. E. et al. Expanded encyclopaedias of DNA elements in the human and mouse genomes. Nature 583, 699–710 (2020).

75. Dobin, A. et al. STAR: ultrafast universal RNA-seq aligner. Bioinformatics 29, 15–21 (2013).

76. Langmead, B. & Salzberg, S. L. Fast gapped-read alignment with Bowtie 2. Nat. Methods 9, 357–359 (2012).

77. Gaspar, J. M. Improved peak-calling with MACS2. 496521 Preprint at 10.1101/496521 (2018).

78. Li, Q., Brown, J. B., Huang, H. & Bickel, P. J. Measuring reproducibility of high-throughput experiments. Ann. Appl. Stat. 5, 1752–1779 (2011).

79. Quinlan, A. R. & Hall, I. M. BEDTools: a flexible suite of utilities for comparing genomic features. Bioinforma. Oxf. Engl. 26, 841–842 (2010).

80. Law, C. W., Chen, Y., Shi, W. & Smyth, G. K. voom: precision weights unlock linear model analysis tools for RNA-seq read counts. Genome Biol. 15, R29 (2014).

81. Yu, G., Wang, L.-G. & He, Q.-Y. ChIPseeker: an R/Bioconductor package for ChIP peak annotation, comparison and visualization. Bioinformatics 31, 2382–2383 (2015).

82. Meissner, A. et al. Reduced representation bisulfite sequencing for comparative high-resolution DNA methylation analysis. Nucleic Acids Res. 33, 5868–5877 (2005).

83. Andrews, S. FastQC: A Quality Control Tool for High Throughput Sequence Data. https://qubeshub.org/resources/fastqc (2015).

84. Krueger, F. & Andrews, S. R. Bismark: a flexible aligner and methylation caller for Bisulfite-Seq applications. Bioinformatics. 27, 1571–2 (2011).

85. Li, H. et al. The Sequence Alignment/Map format and SAMtools. Bioinformatics 25, 2078–2079 (2009).

86. Jäger, M. L. MCL: Markov Cluster Algorithm. https://CRAN.R-project.org/package=MCL (2015).

87. Bolstad, B. preprocessCore: A collection of pre-processing functions. GitHub https://github.com/bmbolstad/preprocessCore (2021).

88. Ewels, P., Magnusson, M., Lundin, S. & Käller, M. MultiQC: summarize analysis results for multiple tools and samples in a single report. Bioinforma. Oxf. Engl. 32, 3047–3048 (2016).

89. Love, M. I., Huber, W. & Anders, S. Moderated estimation of fold change and dispersion for RNA-seq data with DESeq2. Genome Biol. 15, 550 (2014).

90. Robinson, M. D., McCarthy, D. J. & Smyth, G. K. edgeR: a Bioconductor package for differential expression analysis of digital gene expression data. Bioinforma. Oxf. Engl. 26, 139– 140 (2010).

91. Mertins, P. et al. Reproducible workflow for multiplexed deep-scale proteome and phosphoproteome analysis of tumor tissues by liquid chromatography–mass spectrometry. Nat. Protoc. 13, 1632–1661 (2018).

92. Zecha, J. et al. TMT Labeling for the Masses: A Robust and Cost-efficient, In-solution Labeling Approach *[S]. Mol. Cell. Proteomics 18, 1468–1478 (2019).

93. Ritchie, M. E. et al. limma powers differential expression analyses for RNA-sequencing and microarray studies. Nucleic Acids Res. 43, e47 (2015).

94. Chen, Y., Pal, B., Visvader, J. E. & Smyth, G. K. Differential methylation analysis of reduced representation bisulfite sequencing experiments using edgeR. F1000Research 6, 2055 (2017).

95. Ignatiadis, N., Klaus, B., Zaugg, J. B. & Huber, W. Data-driven hypothesis weighting increases detection power in genome-scale multiple testing. Nat. Methods 13, 577–580 (2016).

96. Heinz, S. et al. Simple combinations of lineage-determining transcription factors prime cis- regulatory elements required for macrophage and B cell identities. Mol. Cell 38, 576–589 (2010).

97. Chikina, M., Zaslavsky, E. & Sealfon, S. C. CellCODE: a robust latent variable approach to differential expression analysis for heterogeneous cell populations. Bioinformatics 31, 1584– 1591 (2015).

98. Abbas, A. R., Wolslegel, K., Seshasayee, D., Modrusan, Z. & Clark, H. F. Deconvolution of Blood Microarray Data Identifies Cellular Activation Patterns in Systemic Lupus Erythematosus. PLOS ONE 4, e6098 (2009).

99. Novershtern, N. et al. Densely Interconnected Transcriptional Circuits Control Cell States in Human Hematopoiesis. Cell 144, 296–309 (2011).

100. Raudvere, U. et al. g:Profiler: a web server for functional enrichment analysis and conversions of gene lists (2019 update). Nucleic Acids Res. 47, W191–W198 (2019).

