## Supplemental Figures for "Multi-omic identification of key transcriptional regulatory programs during endurance exercise training in rats"

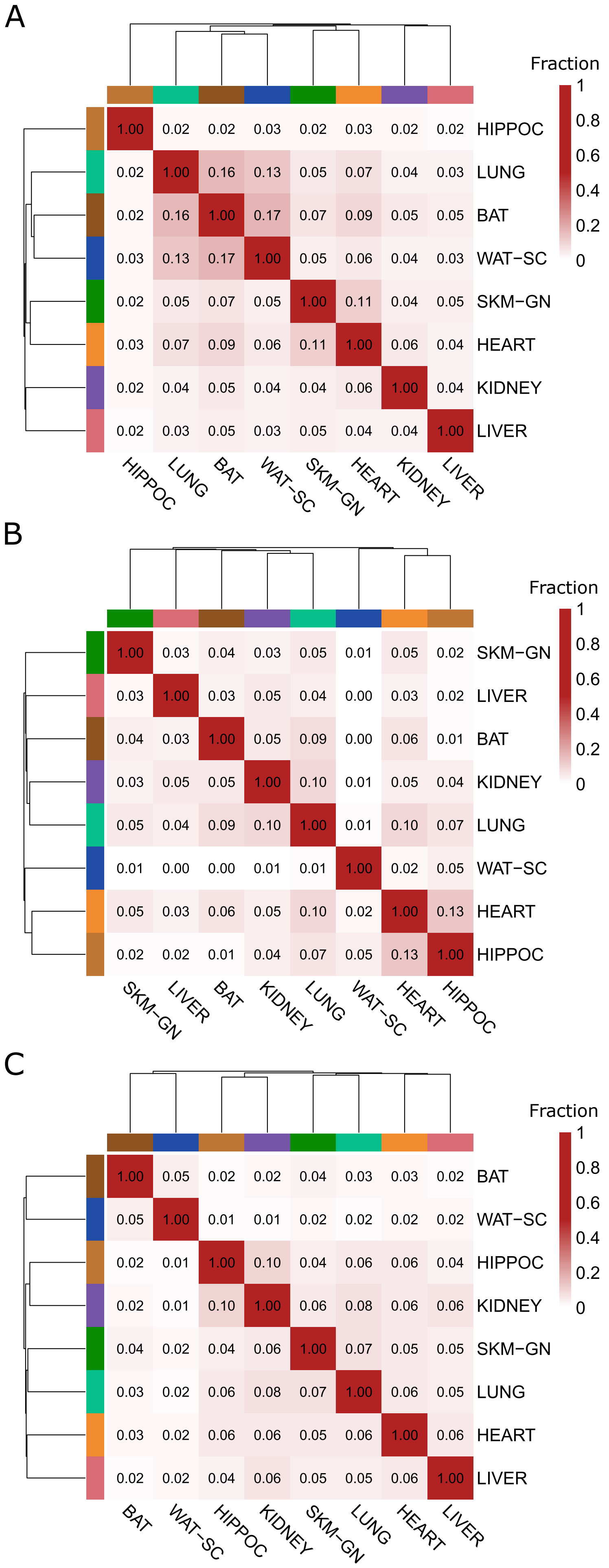
A
Fraction
1
0.8
0.6
0.4
0.2
0
HIPPOC 1.00 0.02 0.02 0.03 0.02 0.03 0.02 0.02
LUNG 0.02 1.00 0.16 0.13 0.05 0.07 0.04 0.03
BAT 0.02 0.16 1.00 0.17 0.07 0.09 0.05 0.05
WAT−SC 0.03 0.13 0.17 1.00 0.05 0.06 0.04 0.03
SKM−GN 0.02 0.05 0.07 0.05 1.00 0.11 0.04 0.05
HEART 0.03 0.07 0.09 0.06 0.11 1.00 0.06 0.04
KIDNEY 0.02 0.04 0.05 0.04 0.04 0.06 1.00 0.04
LIVER 0.02 0.03 0.05 0.03 0.05 0.04 0.04 1.00
HIPPOC LUNG BAT WAT−SC SKM−GN HEART KIDNEY LIVER
B
Fraction
1
0.8
0.6
0.4
0.2
0
SKM−GN 1.00 0.03 0.04 0.03 0.05 0.01 0.05 0.02
LIVER 0.03 1.00 0.03 0.05 0.04 0.00 0.03 0.02
BAT 0.04 0.03 1.00 0.05 0.09 0.00 0.06 0.01
KIDNEY 0.03 0.05 0.05 1.00 0.10 0.01 0.05 0.04
LUNG 0.05 0.04 0.09 0.10 1.00 0.01 0.10 0.07
WAT−SC 0.01 0.00 0.00 0.01 0.01 1.00 0.02 0.05
HEART 0.05 0.03 0.06 0.05 0.10 0.02 1.00 0.13
HIPPOC 0.02 0.02 0.01 0.04 0.07 0.05 0.13 1.00
SKM−GN LIVER BAT KIDNEY LUNG WAT−SC HEART HIPPOC
C
Fraction
1
0.8
0.6
0.4
0.2
0
BAT 1.00 0.05 0.02 0.02 0.04 0.03 0.03 0.02
WAT−SC 0.05 1.00 0.01 0.01 0.02 0.02 0.02 0.02
HIPPOC 0.02 0.01 1.00 0.10 0.04 0.06 0.06 0.04
KIDNEY 0.02 0.01 0.10 1.00 0.06 0.08 0.06 0.06
SKM−GN 0.04 0.02 0.04 0.06 1.00 0.07 0.05 0.05
LUNG 0.03 0.02 0.06 0.08 0.07 1.00 0.06 0.05
HEART 0.03 0.02 0.06 0.06 0.05 0.06 1.00 0.06
LIVER 0.02 0.02 0.04 0.06 0.05 0.05 0.06 1.00
BAT WAT−SC HIPPOC KIDNEY SKM−GN LUNG HEART LIVER

Supplemental Figure S1: Overlap in DEGs, DARs, and DMRs across tissues. (a) Proportion of intersection of DEG lists divided by the union of DEG lists for each pair of tissues. Tissues clustered by overlap. (b) Proportion of intersection of DAR lists divided by the union of DAR lists for each pair of tissues. Tissues clustered by overlap. (c) Proportion of intersection of DMR lists divided by the union of DMR lists for each pair of tissues. Tissues clustered by overlap.

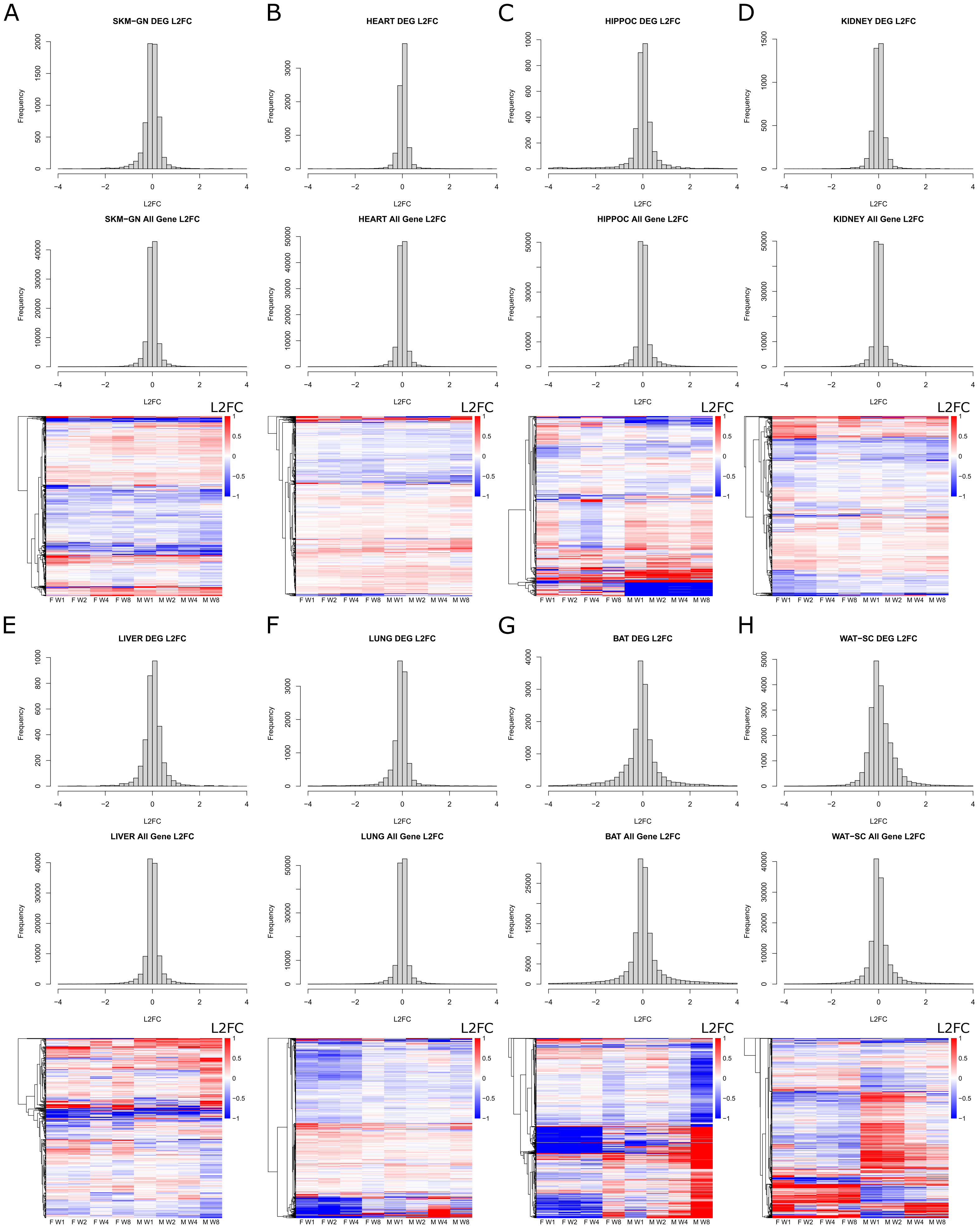

A
SKM−GN DEG L2FC
SKM−GN All Gene L2FC
B
HEART DEG L2FC
HEART All Gene L2FC
C
HIPPOC DEG L2FC
HIPPOC All Gene L2FC
D
KIDNEY DEG L2FC
KIDNEY All Gene L2FC
E
LIVER DEG L2FC
LIVER All Gene L2FC
F
LUNG DEG L2FC
LUNG All Gene L2FC
G
BAT DEG L2FC
BAT All Gene L2FC
H
WAT−SC DEG L2FC
WAT−SC All Gene L2FC
Frequency
L2FC
F W1 F W2 F W4 F W8 M W1 M W2 M W4 M W8

Supplemental Figure S2: Distribution of training responses for DEGs in eight tissues. (a-h) A histogram of L2FC values across sexes and time points for all DEGs, a separate histogram of L2FC values across sexes and time points for all genes, and a heatmap of L2FC values for DEGs ordered by hierarchical clustering for skeletal muscle (a), heart (b), hippocampus (c), kidney (d), liver (e), lung (f), brown adipose (e), and white adipose (f).

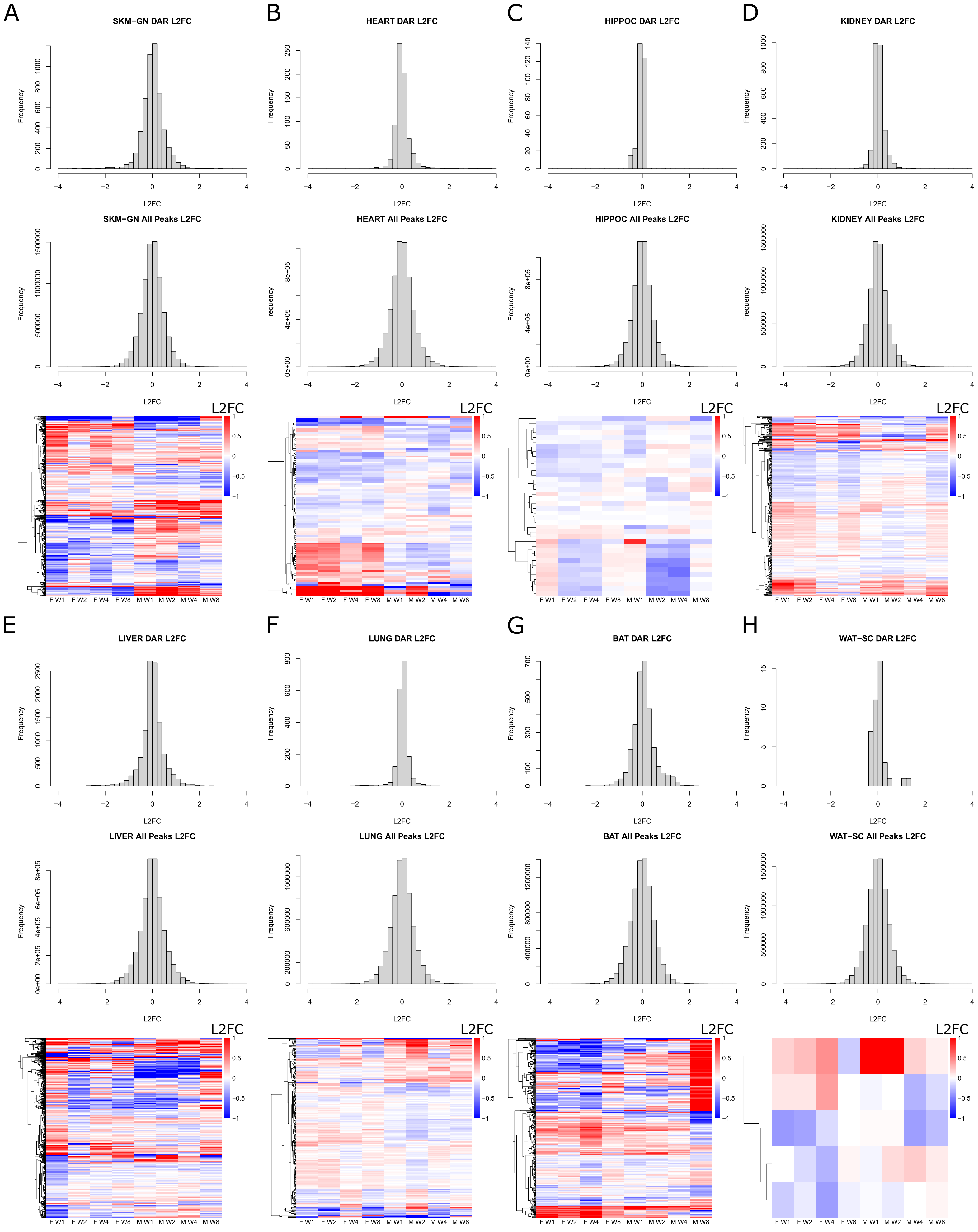
A
SKM−GN DAR L2FC
SKM−GN All Peaks L2FC
B
HEART DAR L2FC
HEART All Peaks L2FC
C
HIPPOC DAR L2FC
HIPPOC All Peaks L2FC
D
KIDNEY DAR L2FC
KIDNEY All Peaks L2FC
E
LIVER DAR L2FC
LIVER All Peaks L2FC
F
LUNG DAR L2FC
LUNG All Peaks L2FC
G
BAT DAR L2FC
BAT All Peaks L2FC
H
WAT−SC DAR L2FC
WAT−SC All Peaks L2FC
Frequency
L2FC
F W1
F W2
F W4
F W8
M W1
M W2
M W4
M W8

Supplemental Figure S3: Distribution of training responses for DARs in eight tissues. (a-h) A histogram of L2FC values across sexes and time points for all DARs, a separate histogram of L2FC values across sexes and time points for all peaks, and a heatmap of L2FC values for DARs ordered by hierarchical clustering for skeletal muscle (a), heart (b), hippocampus (c), kidney (d), liver (e), lung (f), brown adipose (e), and white adipose (f).

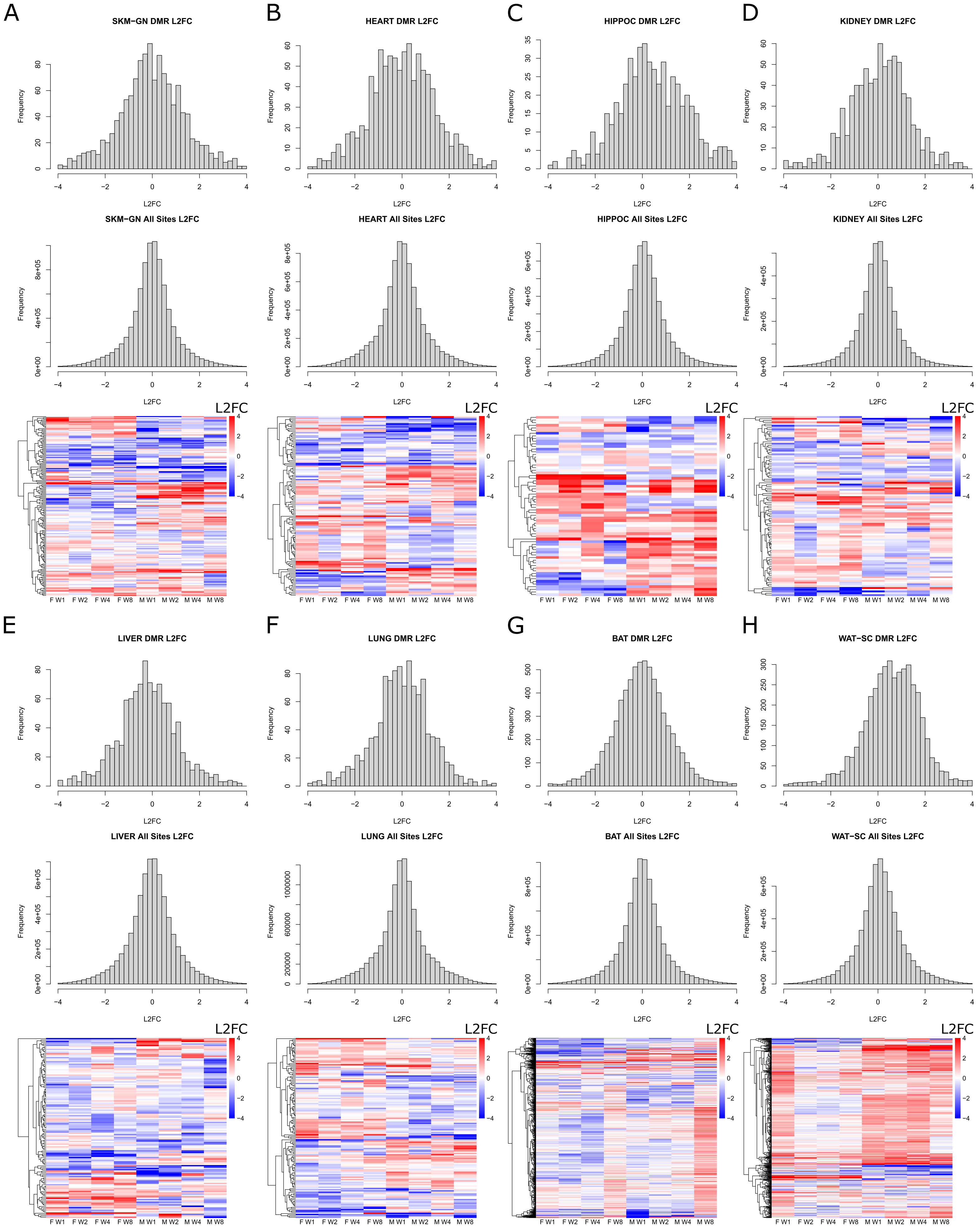
A
SKM−GN DMR L2FC
SKM−GN All Sites L2FC
B
HEART DMR L2FC
HEART All Sites L2FC
C
HIPPOC DMR L2FC
HIPPOC All Sites L2FC
D
KIDNEY DMR L2FC
KIDNEY All Sites L2FC
E
LIVER DMR L2FC
LIVER All Sites L2FC
F
LUNG DMR L2FC
LUNG All Sites L2FC
G
BAT DMR L2FC
BAT All Sites L2FC
H
WAT−SC DMR L2FC
WAT−SC All Sites L2FC
Frequency
L2FC
F W1
F W2
F W4
F W8
M W1
M W2
M W4
M W8

Supplemental Figure S4: Distribution of training responses for DMRs in eight tissues. (a-h) A histogram of L2FC values across sexes and time points for all DMRs, a separate histogram of L2FC values across sexes and time points for all methylation regions, and a heatmap of L2FC values for DMRs ordered by hierarchical clustering for skeletal muscle (a), heart (b), hippocampus (c), kidney (d), liver (e), lung (f), brown adipose (e), and white adipose (f).

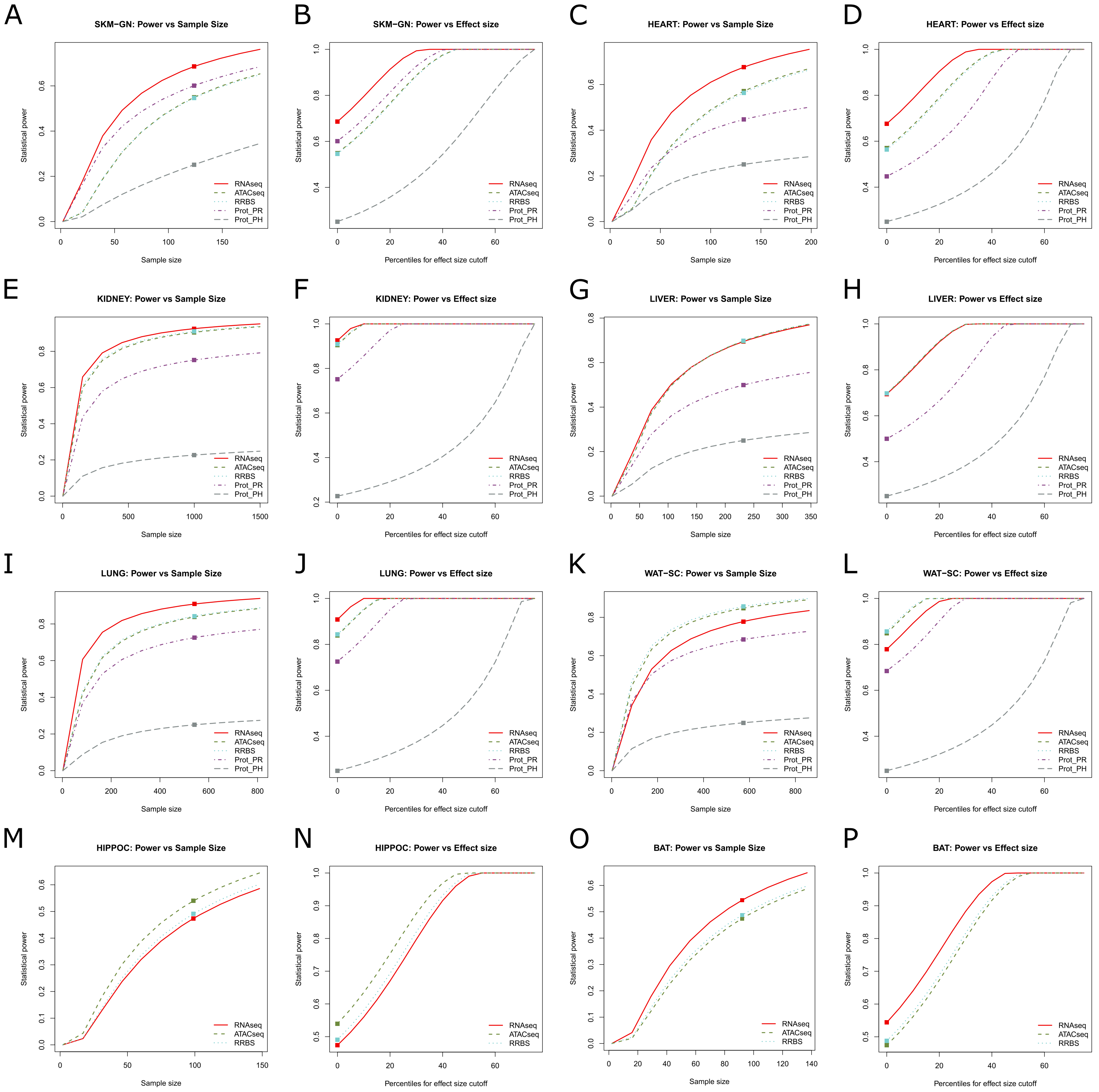
A
SKM−GN: Power vs Sample Size
B
SKM−GN: Power vs Effect size
C
HEART: Power vs Sample Size
D
HEART: Power vs Effect size
E
KIDNEY: Power vs Sample Size
F
KIDNEY: Power vs Effect size
G
LIVER: Power vs Sample Size
H
LIVER: Power vs Effect size
I
LUNG: Power vs Sample Size
J
LUNG: Power vs Effect size
K
WAT−SC: Power vs Sample Size
L
WAT−SC: Power vs Effect size
M
HIPPOC: Power vs Sample Size
N
HIPPOC: Power vs Effect size
O
BAT: Power vs Sample Size
P
BAT: Power vs Effect size
Statistical power
Sample size
Percentiles for effect size cutoff
RNAseq
ATACseq
RRBS
Prot_PR
Prot_PH

Supplemental Figure S5: Multiomic power analysis in eight tissues for RNAseq, ATACseq, RRBS, protein abundance (Prot_PR) and protein phosphorylation (Prot_PH) data. For each tissue, plots of statistical power for each ome at different sample sizes (a,c,e,g,i,k,m,o) and with different cutoffs for effect size (b,d,f,h,j,l,n,p). The square represents the sample size or cutoff needed to achieve 0.25 power in all omes. Tissues include skeletal muscle (SKM-GN) (a,b), heart (c,d), hippocampus (HIPPOC) (e,f), kidney (g,h), liver (i,j), lung (k,l), brown adipose (BAT) (m,n), white adipose (WAT-SC) (o,p).

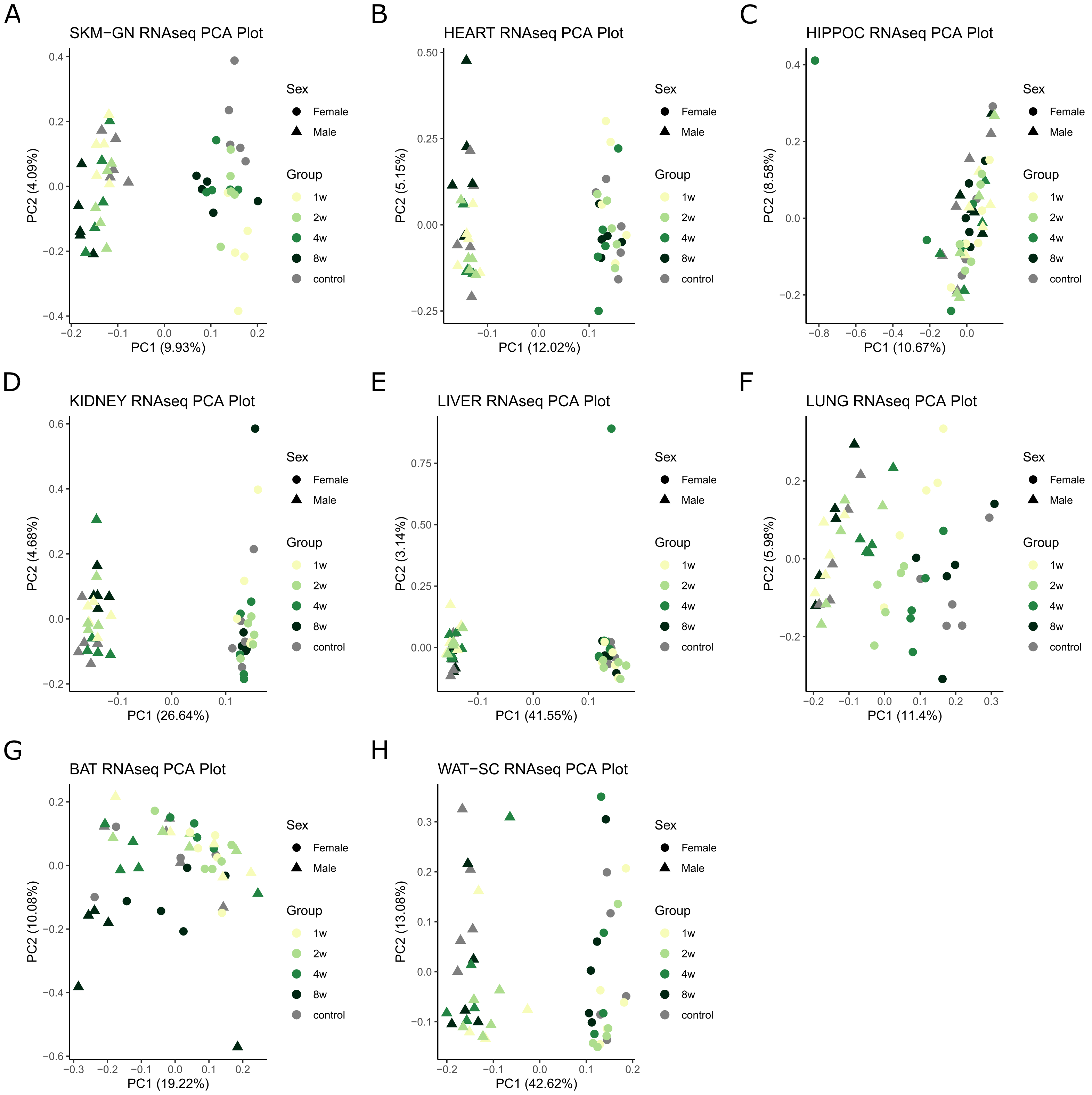
A
SKM−GN RNAseq PCA Plot
PC1 (9.93%)
PC2 (4.09%)
B
HEART RNAseq PCA Plot
PC1 (12.02%)
PC2 (5.15%)
C
HIPPOC RNAseq PCA Plot
PC1 (10.67%)
PC2 (8.58%)
D
KIDNEY RNAseq PCA Plot
PC1 (26.64%)
PC2 (4.68%)
E
LIVER RNAseq PCA Plot
PC1 (41.55%)
PC2 (3.14%)
F
LUNG RNAseq PCA Plot
PC1 (11.4%)
PC2 (5.98%)
G
BAT RNAseq PCA Plot
PC1 (19.22%)
PC2 (10.08%)
H
WAT−SC RNAseq PCA Plot
PC1 (42.62%)
PC2 (13.08%)
Sex
Female
Male
Group
1w
2w
4w
8w
control

Supplemental Figure S6: PCA plots of top two PCs for RNA-seq data in eight tissues: skeletal muscle (a), heart (b), hippocampus (c), kidney (d), liver (e), lung (f), brown adipose (e), and white adipose (f).

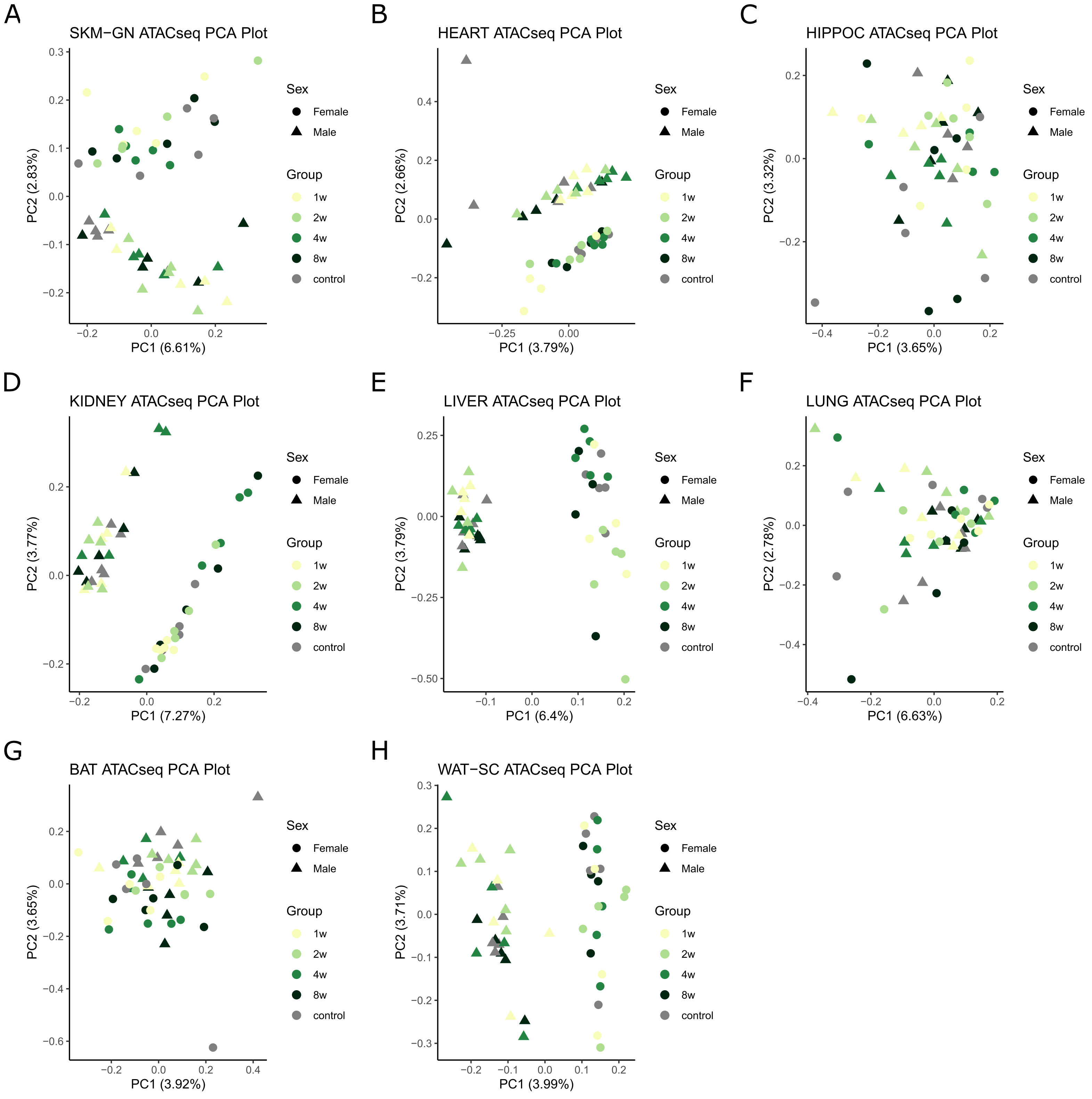
A
SKM−GN ATACseq PCA Plot
PC1 (6.61%)
PC2 (2.83%)
B
HEART ATACseq PCA Plot
PC1 (3.79%)
PC2 (2.66%)
C
HIPPOC ATACseq PCA Plot
PC1 (3.65%)
PC2 (3.32%)
D
KIDNEY ATACseq PCA Plot
PC1 (7.27%)
PC2 (3.77%)
E
LIVER ATACseq PCA Plot
PC1 (6.4%)
PC2 (3.79%)
F
LUNG ATACseq PCA Plot
PC1 (6.63%)
PC2 (2.78%)
G
BAT ATACseq PCA Plot
PC1 (3.92%)
PC2 (3.65%)
H
WAT−SC ATACseq PCA Plot
PC1 (3.99%)
PC2 (3.71%)
Sex
Female
Male
Group
1w
2w
4w
8w
control

Supplemental Figure S7: PCA plots of top two PCs for ATAC-seq data in eight tissues: skeletal muscle (a), heart (b), hippocampus (c), kidney (d), liver (e), lung (f), brown adipose (e), and white adipose (f).

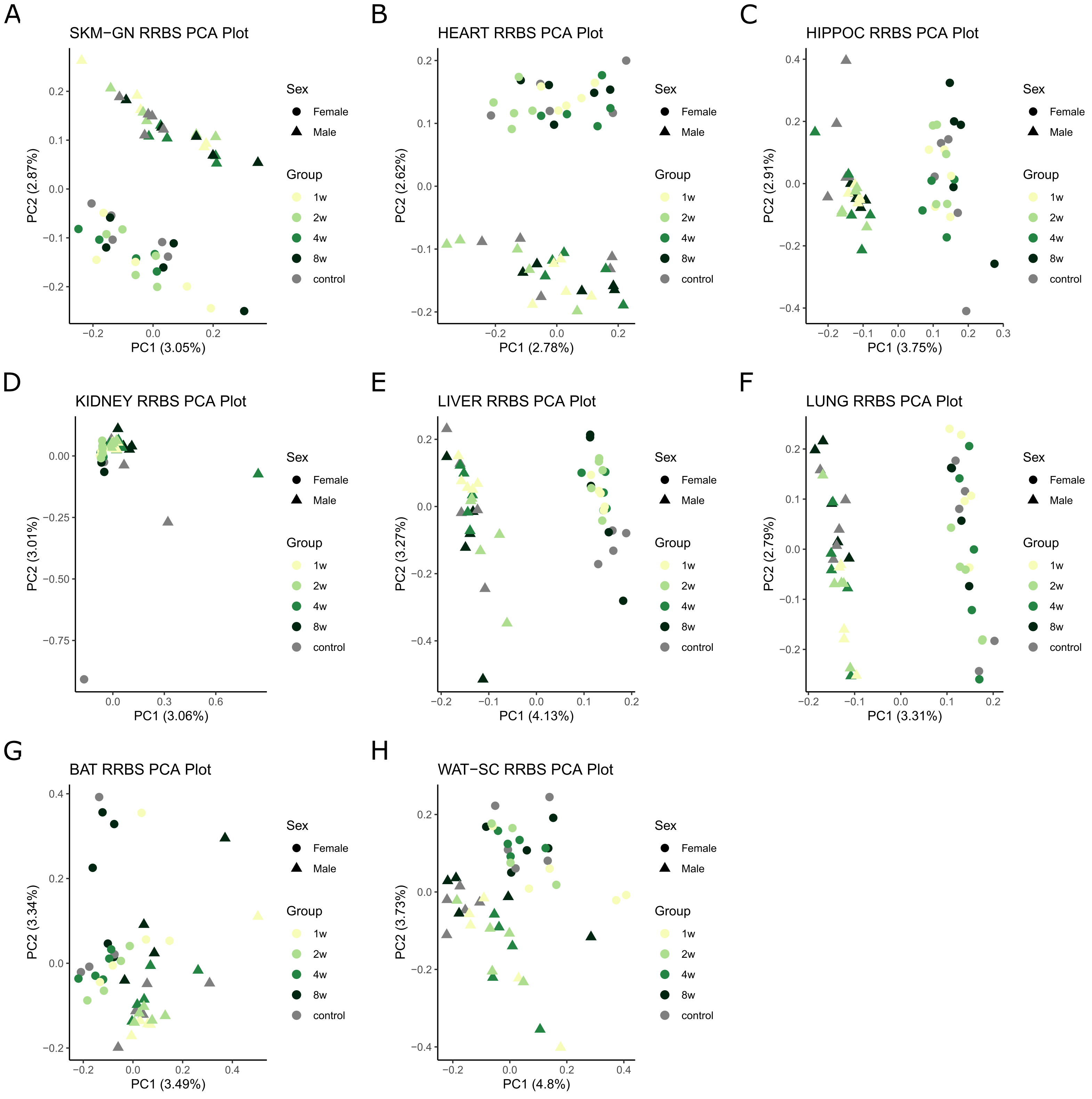
A
SKM−GN RRBS PCA Plot
PC1 (3.05%)
PC2 (2.87%)
B
HEART RRBS PCA Plot
PC1 (2.78%)
PC2 (2.62%)
C
HIPPOC RRBS PCA Plot
PC1 (3.75%)
PC2 (2.91%)
D
KIDNEY RRBS PCA Plot
PC1 (3.06%)
PC2 (3.01%)
E
LIVER RRBS PCA Plot
PC1 (4.13%)
PC2 (3.27%)
F
LUNG RRBS PCA Plot
PC1 (3.31%)
PC2 (2.79%)
G
BAT RRBS PCA Plot
PC1 (3.49%)
PC2 (3.34%)
H
WAT−SC RRBS PCA Plot
PC1 (4.8%)
PC2 (3.73%)
Sex
Female
Male
Group
1w
2w
4w
8w
control

Supplemental Figure S8: PCA plots of top two PCs for RRBS data in eight tissues: skeletal muscle (a), heart (b), hippocampus (c), kidney (d), liver (e), lung (f), brown adipose (e), and white adipose (f).

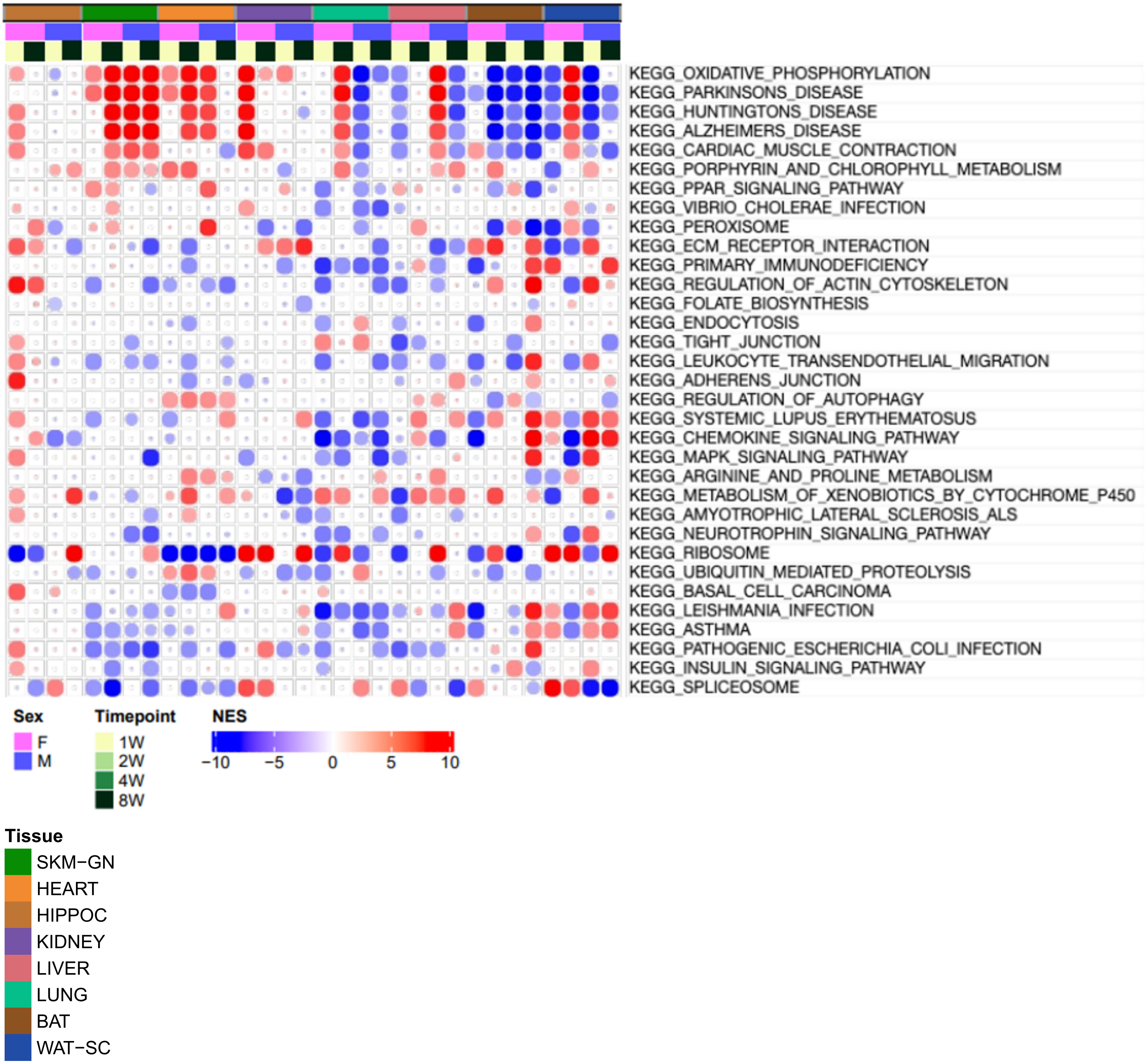
KEGG_OXIDATIVE_PHOSPHORYLATION
KEGG_PARKINSONS_DISEASE
KEGG_HUNTINGTONS_DISEASE
KEGG_ALZHEIMERS_DISEASE
KEGG_CARDIAC_MUSCLE_CONTRACTION
KEGG_PORPHYRIN_AND_CHLOROPHYLL_METABOLISM
KEGG_PPAR_SIGNALING_PATHWAY
KEGG_VIBRIO_CHOLERAE_INFECTION
KEGG_PEROXISOME
KEGG_ECM_RECEPTOR_INTERACTION
KEGG_PRIMARY_IMMUNODEFICIENCY
KEGG_REGULATION_OF_ACTIN_CYTOSKELETON
KEGG_FOLATE_BIOSYNTHESIS
KEGG_ENDOCYTOSIS
KEGG_TIGHT_JUNCTION
KEGG_LEUKOCYTE_TRANSENDOTHELIAL_MIGRATION
KEGG_ADHERENS_JUNCTION
KEGG_REGULATION_OF_AUTOPHAGY
KEGG_SYSTEMIC_LUPUS_ERYTHEMATOSUS
KEGG_CHEMOKINE_SIGNALING_PATHWAY
KEGG_MAPK_SIGNALING_PATHWAY
KEGG_ARGININE_AND_PROLINE_METABOLISM
KEGG_METABOLISM_OF_XENOBIOTICS_BY_CYTOCHROME_P450
KEGG_AMYOTROPHIC_LATERAL_SCLEROSIS_ALS
KEGG_NEUROTROPHIN_SIGNALING_PATHWAY
KEGG_RIBOSOME
KEGG_UBIQUITIN_MEDIATED_PROTEOLYSIS
KEGG_BASAL_CELL_CARCINOMA
KEGG_LEISHMANIA_INFECTION
KEGG_ASTHMA
KEGG_PATHOGENIC_ESCHERICHIA_COLI_INFECTION
KEGG_INSULIN_SIGNALING_PATHWAY
KEGG_SPLICEOSOME
Sex
F
M
Timepoint
1W
2W
4W
8W
NES
−10
−5
0
5
10
Tissue
SKM−GN
HEART
HIPPOC
KIDNEY
LIVER
LUNG
BAT
WAT−SC

Supplemental Figure S9: Variability in GSEA enrichment across tissue, sex and time point. Gene set enrichment analysis (GSEA)-generated normalized enrichment score (NES) for significantly enriched pathways among week 1 and week 8 training response genes in both sexes and eight tissues.

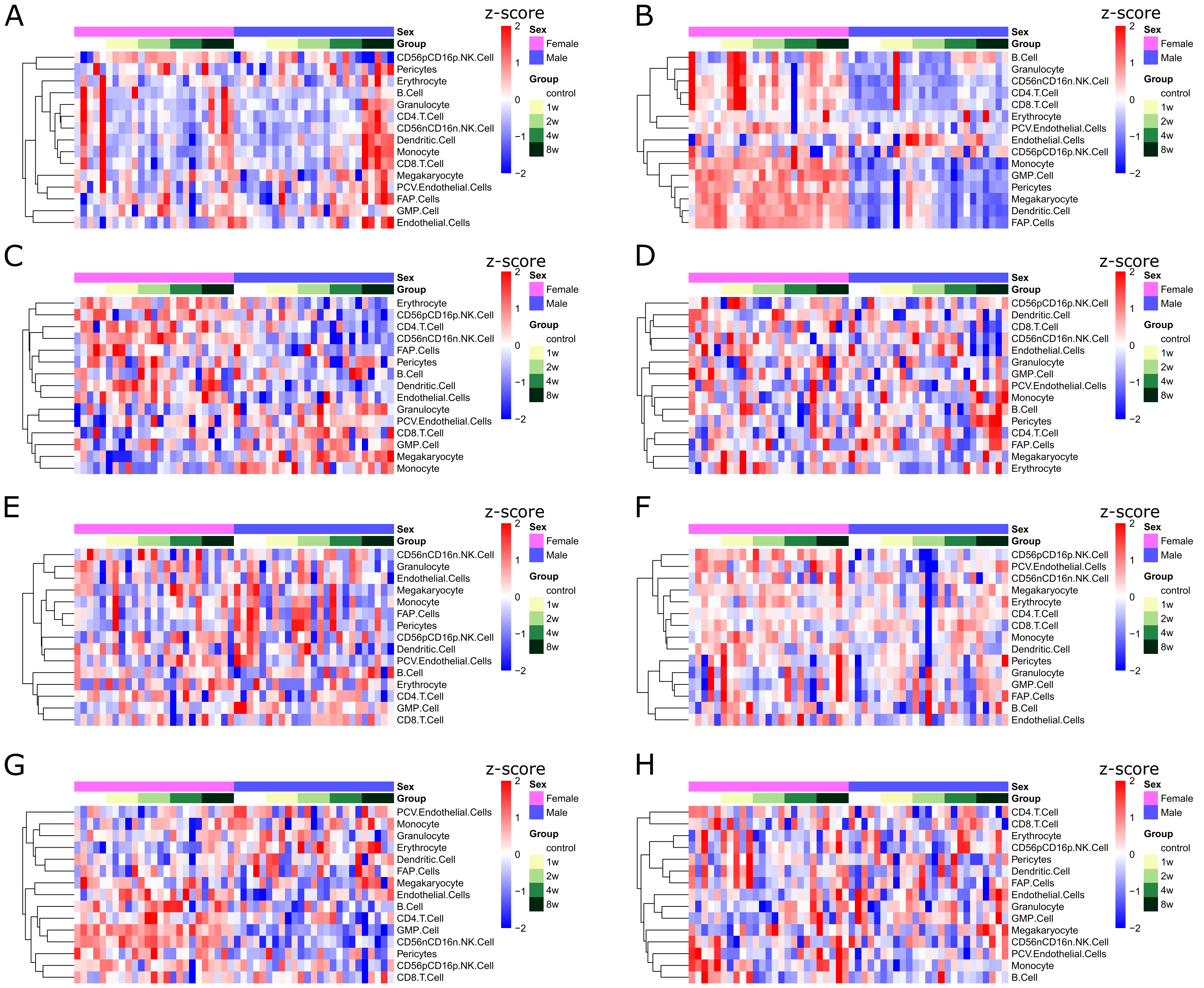
A
z-score
Sex
Group
2
1
0
−1
−2
Sex
Female
Male
Group
control
1w
2w
4w
8w
CD56pCD16p.NK.Cell
Pericytes
Erythrocyte
B.Cell
Granulocyte
CD4.T.Cell
CD56nCD16n.NK.Cell
Dendritic.Cell
Monocyte
CD8.T.Cell
Megakaryocyte
PCV.Endothelial.Cells
FAP.Cells
GMP.Cell
Endothelial.Cells
B
B.Cell
Granulocyte
CD56nCD16n.NK.Cell
CD4.T.Cell
CD8.T.Cell
Erythrocyte
PCV.Endothelial.Cells
Endothelial.Cells
CD56pCD16p.NK.Cell
Monocyte
GMP.Cell
Pericytes
Megakaryocyte
Dendritic.Cell
FAP.Cells
C
Erythrocyte
CD56pCD16p.NK.Cell
CD4.T.Cell
CD56nCD16n.NK.Cell
FAP.Cells
Pericytes
B.Cell
Dendritic.Cell
Endothelial.Cells
Granulocyte
PCV.Endothelial.Cells
CD8.T.Cell
GMP.Cell
Megakaryocyte
Monocyte
D
CD56pCD16p.NK.Cell
Dendritic.Cell
CD8.T.Cell
CD56nCD16n.NK.Cell
Endothelial.Cells
Granulocyte
GMP.Cell
PCV.Endothelial.Cells
Monocyte
B.Cell
Pericytes
CD4.T.Cell
FAP.Cells
Megakaryocyte
Erythrocyte
E
CD56nCD16n.NK.Cell
Granulocyte
Endothelial.Cells
Megakaryocyte
Monocyte
FAP.Cells
Pericytes
CD56pCD16p.NK.Cell
Dendritic.Cell
PCV.Endothelial.Cells
B.Cell
Erythrocyte
CD4.T.Cell
GMP.Cell
CD8.T.Cell
F
CD56pCD16p.NK.Cell
PCV.Endothelial.Cells
CD56nCD16n.NK.Cell
Megakaryocyte
Erythrocyte
CD4.T.Cell
CD8.T.Cell
Monocyte
Dendritic.Cell
Pericytes
Granulocyte
GMP.Cell
FAP.Cells
B.Cell
Endothelial.Cells
G
PCV.Endothelial.Cells
Monocyte
Granulocyte
Erythrocyte
Dendritic.Cell
FAP.Cells
Megakaryocyte
Endothelial.Cells
B.Cell
CD4.T.Cell
GMP.Cell
CD56nCD16n.NK.Cell
Pericytes
CD56pCD16p.NK.Cell
CD8.T.Cell
H
CD4.T.Cell
CD8.T.Cell
Erythrocyte
CD56pCD16p.NK.Cell
Pericytes
Dendritic.Cell
FAP.Cells
Endothelial.Cells
Granulocyte
GMP.Cell
Megakaryocyte
CD56nCD16n.NK.Cell
PCV.Endothelial.Cells
Monocyte
B.Cell

Supplemental Figure S10: Cell type deconvolution of RNA-seq data for eight tissues. (a-h) z-scores of predicted shifts in cell type proportion for each sample in brown adipose (a), white adipose (b), skeletal muscle (c), heart (d), hippocampus (e), kidney (f), liver (g), and lung (h).

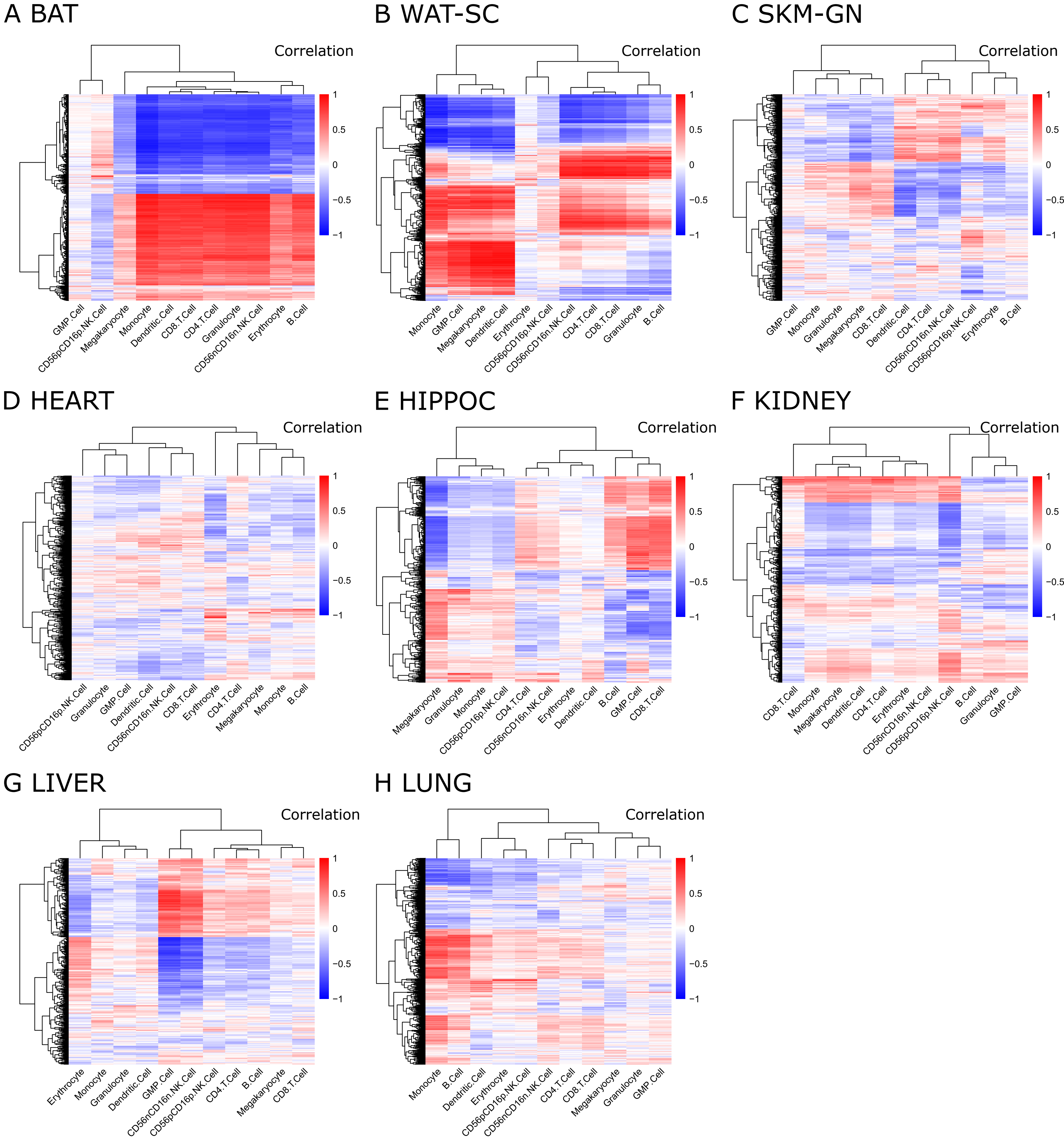
A BAT
Correlation
1
0.5
0
−0.5
−1
GMP.Cell
CD56pCD16p.NK.Cell
Megakaryocyte
Monocyte
Dendritic.Cell
CD8.T.Cell
CD4.T.Cell
Granulocyte
CD56nCD16n.NK.Cell
Erythrocyte
B.Cell
B WAT-SC
Correlation
Monocyte
GMP.Cell
Megakaryocyte
Dendritic.Cell
Erythrocyte
CD56pCD16p.NK.Cell
CD56nCD16n.NK.Cell
CD4.T.Cell
CD8.T.Cell
Granulocyte
B.Cell
C SKM-GN
Correlation
GMP.Cell
Monocyte
Granulocyte
Megakaryocyte
CD8.T.Cell
Dendritic.Cell
CD4.T.Cell
CD56nCD16n.NK.Cell
CD56pCD16p.NK.Cell
Erythrocyte
B.Cell
D HEART
Correlation
CD56pCD16p.NK.Cell
Granulocyte
GMP.Cell
Dendritic.Cell
CD56nCD16n.NK.Cell
CD8.T.Cell
Erythrocyte
CD4.T.Cell
Megakaryocyte
Monocyte
B.Cell
E HIPPOC
Correlation
Megakaryocyte
Granulocyte
Monocyte
CD56pCD16p.NK.Cell
CD4.T.Cell
CD56nCD16n.NK.Cell
Erythrocyte
Dendritic.Cell
B.Cell
GMP.Cell
CD8.T.Cell
F KIDNEY
Correlation
CD8.T.Cell
Monocyte
Megakaryocyte
Dendritic.Cell
CD4.T.Cell
Erythrocyte
CD56nCD16n.NK.Cell
CD56pCD16p.NK.Cell
B.Cell
Granulocyte
GMP.Cell
G LIVER
Correlation
Erythrocyte
Monocyte
Granulocyte
Dendritic.Cell
GMP.Cell
CD56nCD16n.NK.Cell
CD56pCD16p.NK.Cell
CD4.T.Cell
B.Cell
Megakaryocyte
CD8.T.Cell
H LUNG
Correlation
Monocyte
B.Cell
Dendritic.Cell
Erythrocyte
CD56pCD16p.NK.Cell
CD56nCD16n.NK.Cell
CD4.T.Cell
CD8.T.Cell
Megakaryocyte
Granulocyte
GMP.Cell

Supplemental Figure S11: Heatmaps of correlation between the EET response of DEGs and cell type composition changes in eight tissues: BAT (a), WAT-SC (b), SKM-GN (c), heart (d), HIPPOC (e), kidney (f), liver (g), lung (h).

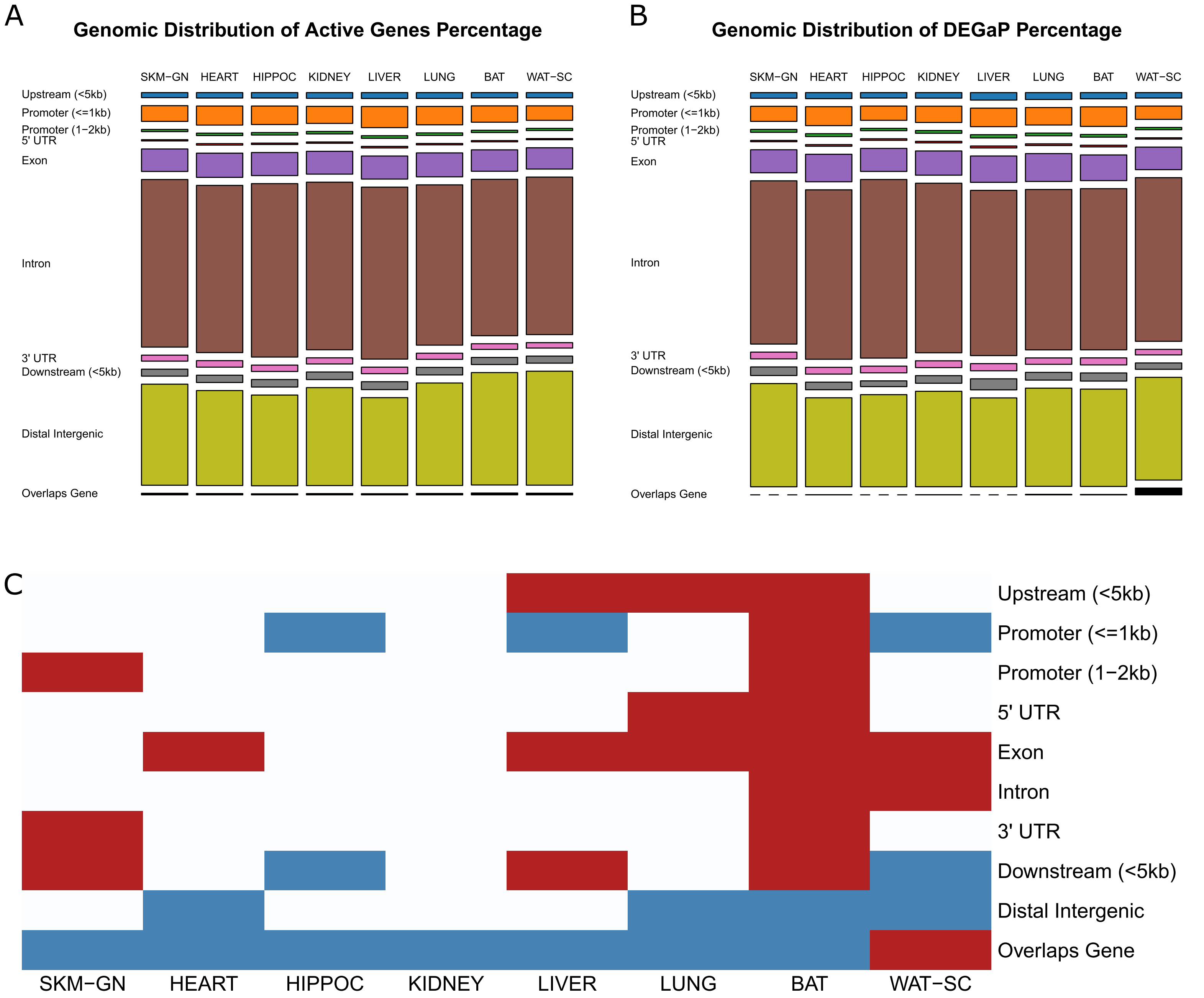
A
Genomic Distribution of Active Genes Percentage
SKM−GN
HEART
HIPPOC
KIDNEY
LIVER
LUNG
BAT
WAT−SC
Upstream (<5kb)
Promoter (<=1kb)
Promoter (1−2kb)
5' UTR
Exon
Intron
3' UTR
Downstream (<5kb)
Distal Intergenic
Overlaps Gene
B
Genomic Distribution of DEGaP Percentage
C

Supplemental Figure S12: Comparison of accessible peak distribution between expressed genes and DEGs. (A) Distribution of genomic regions among the accessible peaks for expressed genes in each tissue. (B) Distribution of genomic regions among the accessible peaks for DEGs (DEGaPs) in each tissue. (c) DEGaPs are enriched in exons for a majority of tissues and exhibit a decreased proportion in distal intergenic regions for half of tissues.

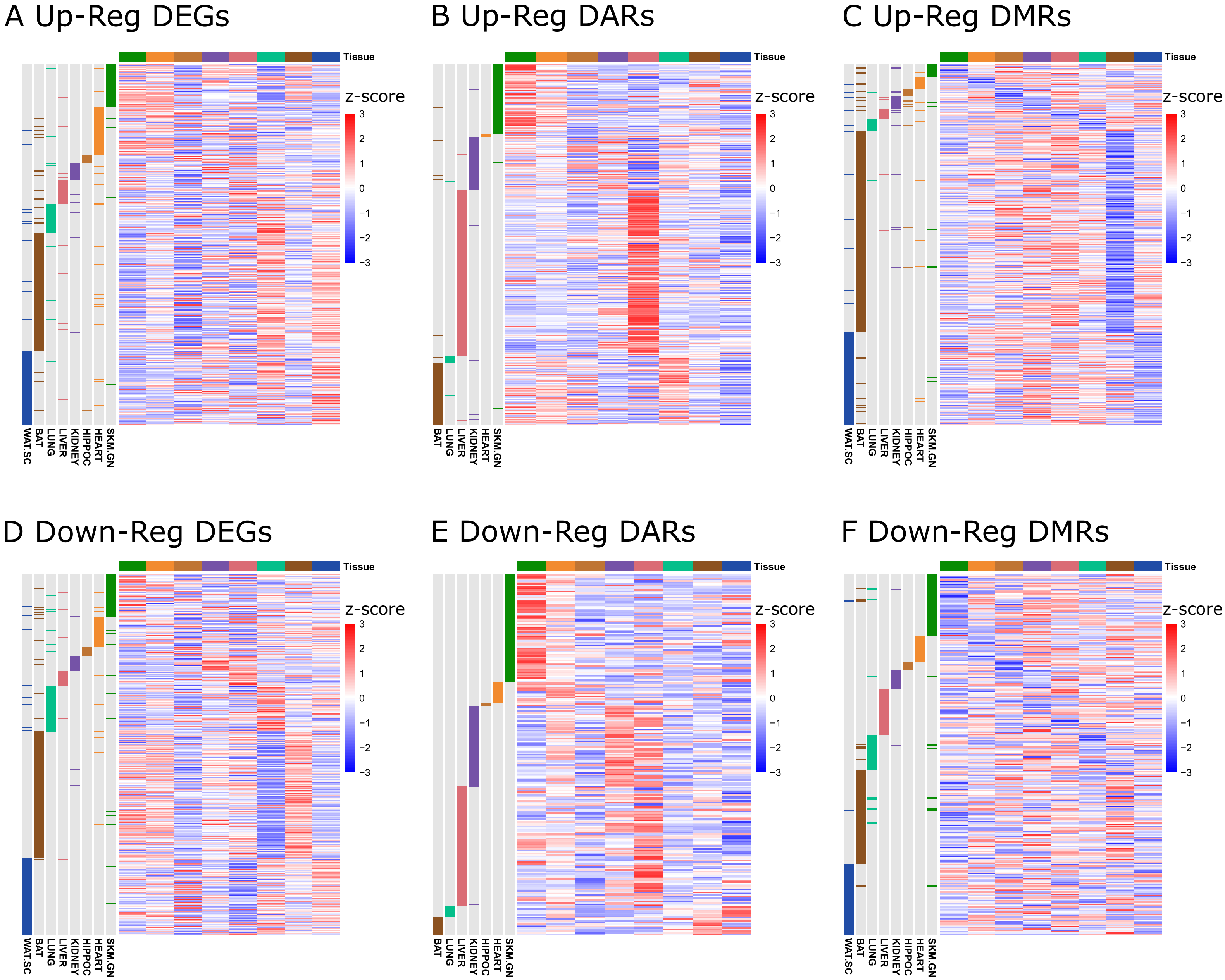
A Up-Reg DEGs
B Up-Reg DARs
C Up-Reg DMRs
D Down-Reg DEGs
E Down-Reg DARs
F Down-Reg DMRs
Tissue
z-score
3
2
1
0
−1
−2
−3
WAT.SC
BAT
LUNG
LIVER
KIDNEY
HIPPOC
HEART
SKM.GN

Supplemental Figure S13: (a-c) Heatmaps of all up-regulated differential analytes across eight tissues in RNA-seq (a), ATAC-seq (b), and RRBS (c). Columns reflect z scores of baseline expression (a), accessibility (b), or methylation (c), across the eight tissues. Rows are annotated by the tissues in which each analyte exhibits a differential training response. (d-f) Heatmaps of all down-regulated differential analytes across eight tissues in RNA-seq (d), ATAC-seq (e), and RRBS (f). Columns reflect z scores of baseline expression (d), accessibility (e), or methylation (f), across the eight tissues. Rows are annotated by the tissues in which each analyte exhibits a differential training response.

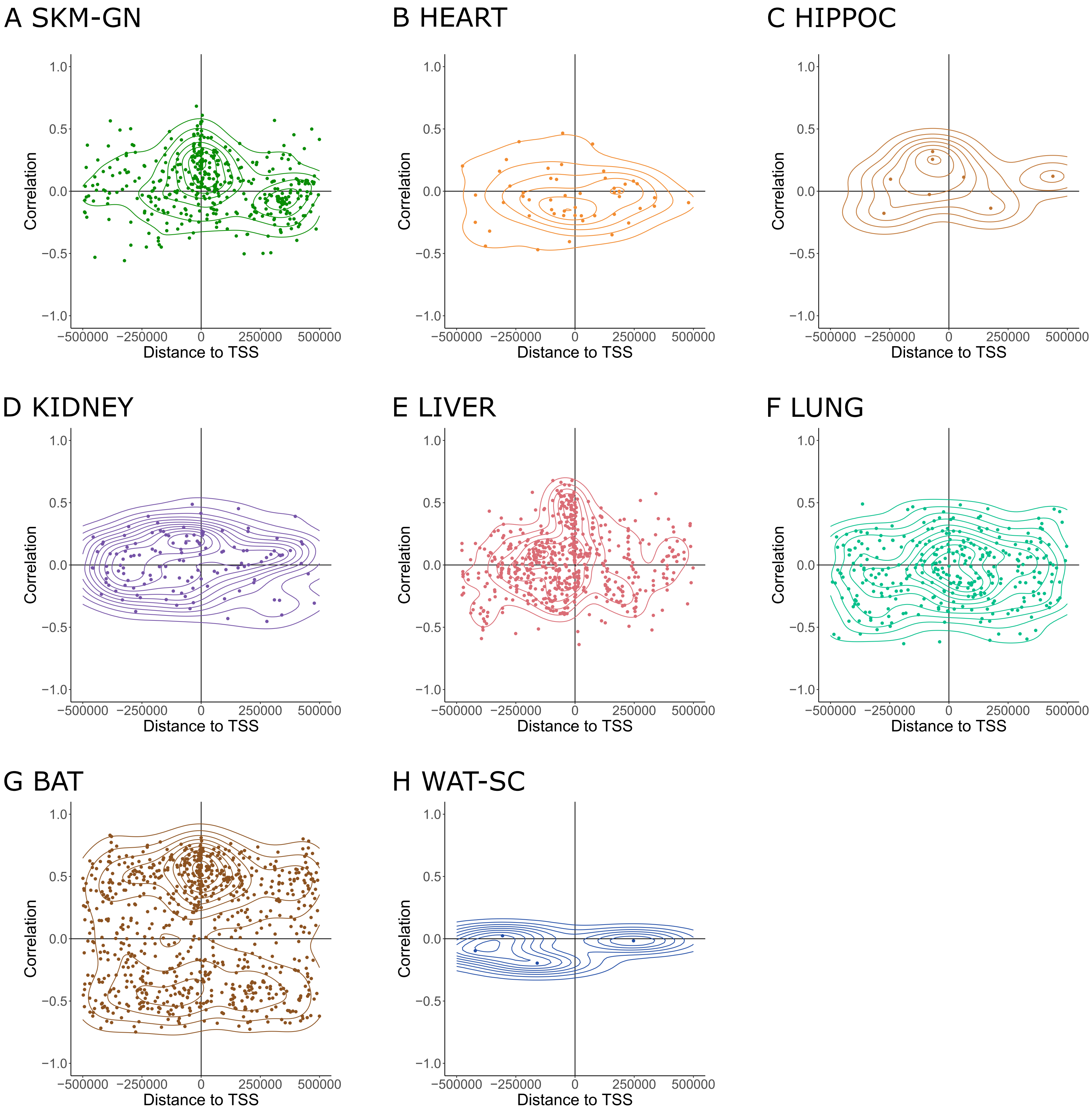

A SKM-GN
B HEART
C HIPPOC
D KIDNEY
E LIVER
F LUNG
G BAT
H WAT-SC
Correlation
Distance to TSS
1.0
0.5
0.0
−0.5
−1.0
−500000
−250000
0
250000
500000

Supplemental Figure S14: Density scatter plots of DAR-DEG distance and training response correlation for eight tissues: skeletal muscle (a), heart (b), hippocampus (c), kidney (d), liver (e), lung (f), brown adipose (e), and white adipose (f).

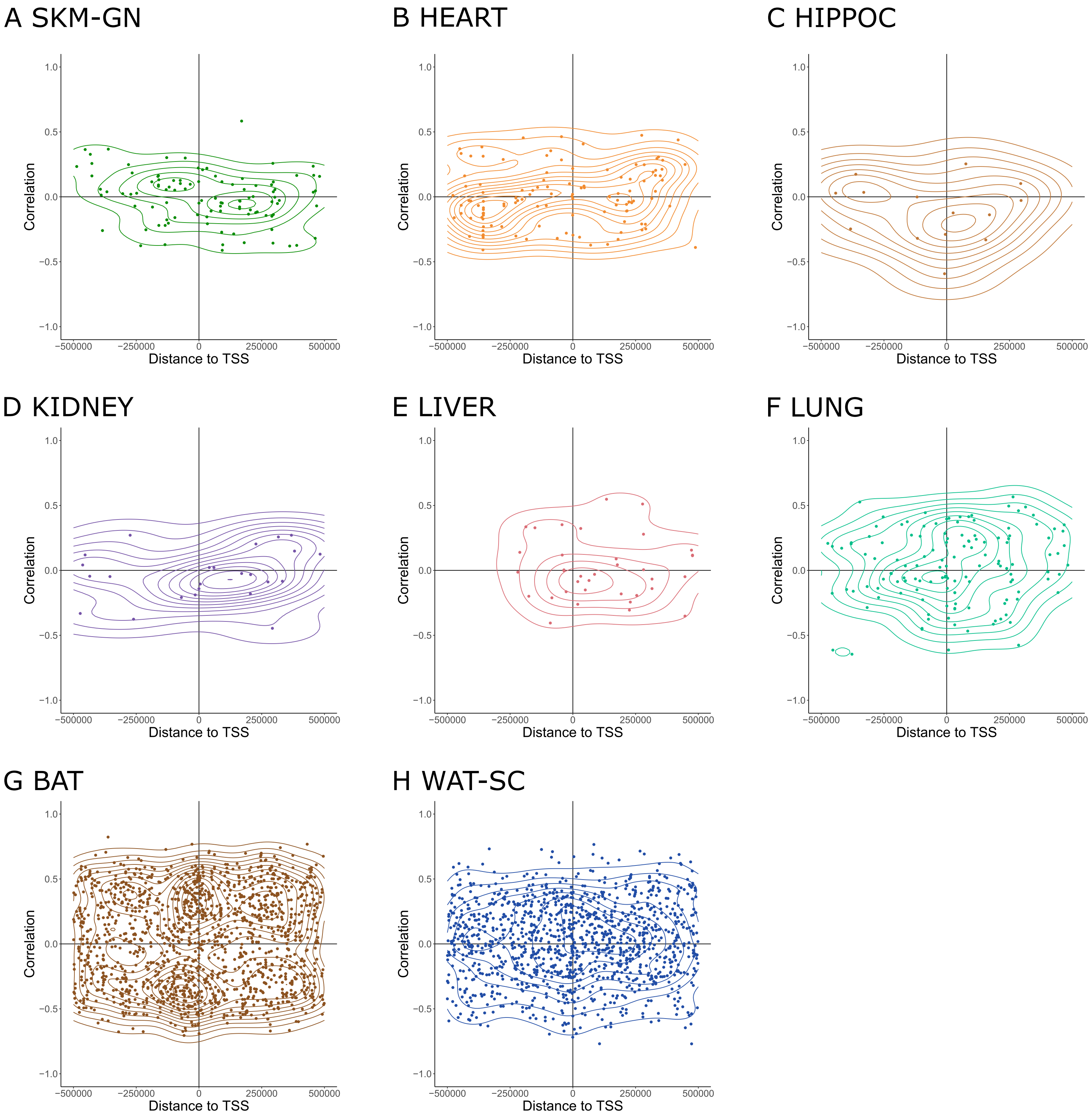

A SKM-GN
B HEART
C HIPPOC
D KIDNEY
E LIVER
F LUNG
G BAT
H WAT-SC
Correlation
Distance to TSS
1.0
0.5
0.0
−0.5
−1.0
−500000
−250000
0
250000
500000

Supplemental Figure S15: Density scatter plots of DMR-DEG distance and training response correlation for eight tissues: skeletal muscle (a), heart (b), hippocampus (c), kidney (d), liver (e), lung (f), brown adipose (e), and white adipose (f).

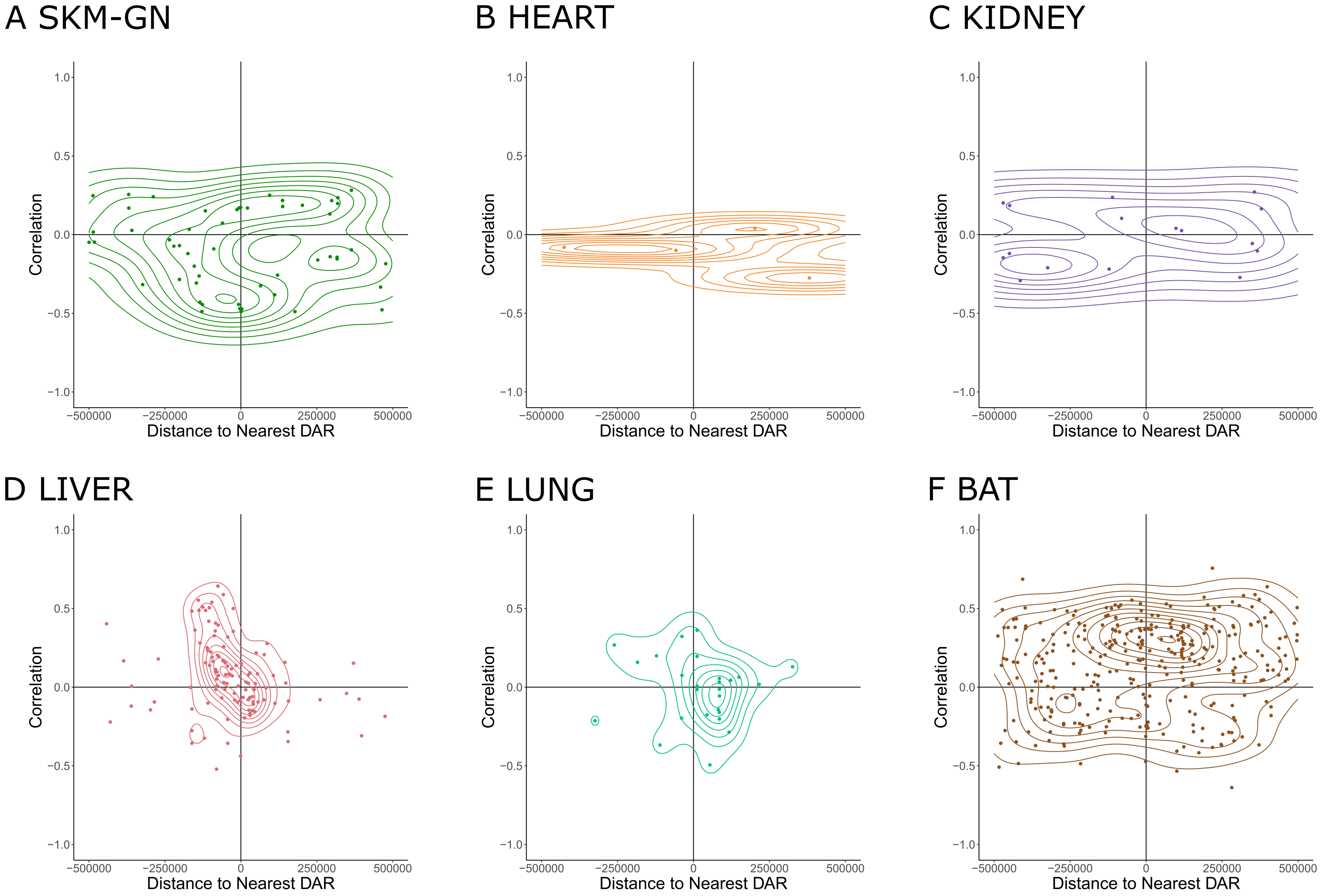
A SKM-GN
B HEART
C KIDNEY
D LIVER
E LUNG
F BAT
Correlation
Distance to Nearest DAR
1.0
0.5
0.0
−0.5
−1.0
−500000
−250000
0
250000
500000

Supplemental Figure S16: Density scatter plots of DMR-DAR distance and training response correlation for six tissues with non-zero measurements of neighboring DMR-DAR pairs: skeletal muscle (a), heart (b), kidney (c), liver (d), lung (e), and brown adipose (f).

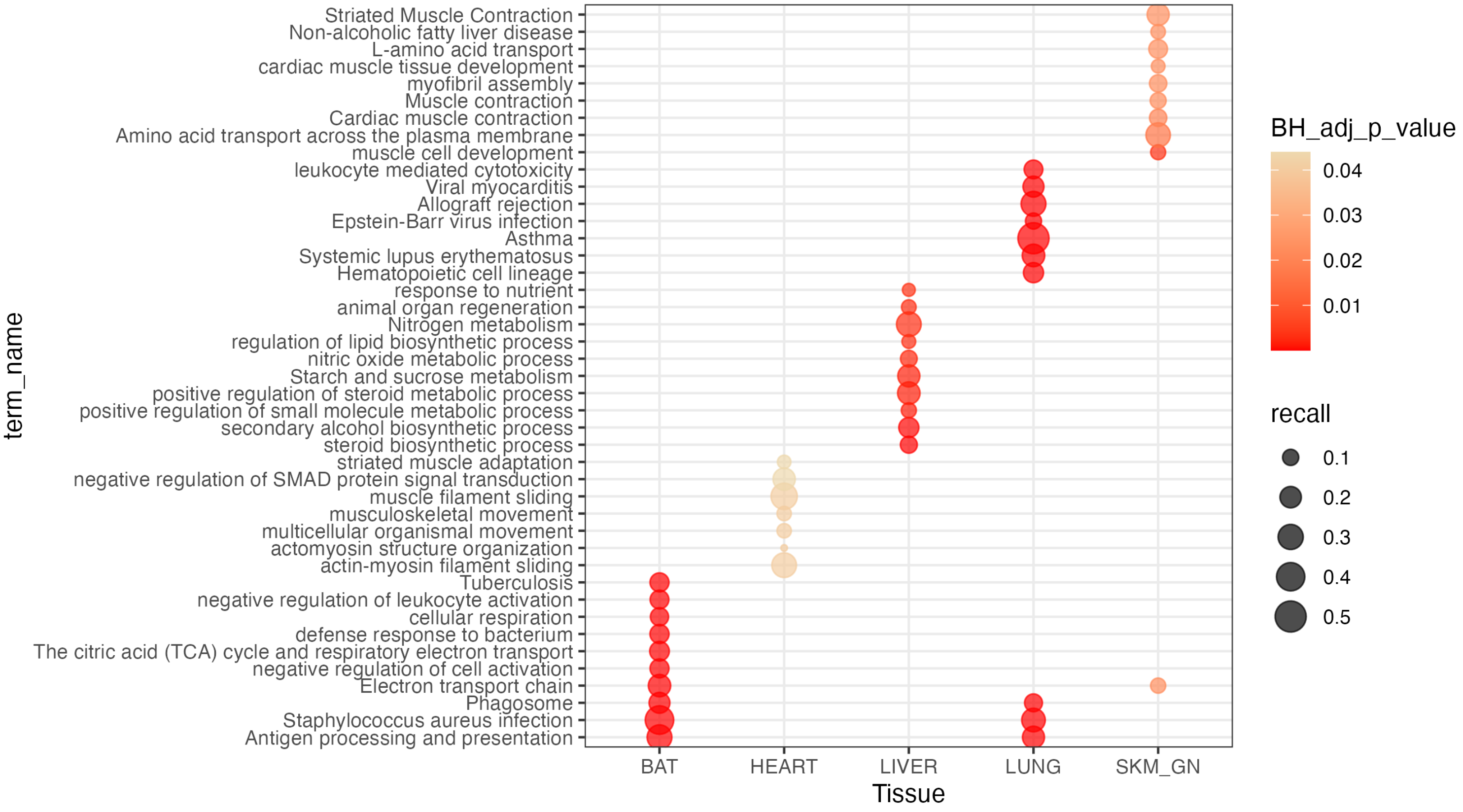

Striated Muscle Contraction
Non-alcoholic fatty liver disease
L-amino acid transport
cardiac muscle tissue development
myofibril assembly
Muscle contraction
Cardiac muscle contraction
Amino acid transport across the plasma membrane
muscle cell development
leukocyte mediated cytotoxicity
Viral myocarditis
Allograft rejection
Epstein-Barr virus infection
Asthma
Systemic lupus erythematosus
Hematopoietic cell lineage
response to nutrient
animal organ regeneration
Nitrogen metabolism
regulation of lipid biosynthetic process
nitric oxide metabolic process
Starch and sucrose metabolism
positive regulation of steroid metabolic process
positive regulation of small molecule metabolic process
secondary alcohol biosynthetic process
steroid biosynthetic process
striated muscle adaptation
negative regulation of SMAD protein signal transduction
muscle filament sliding
musculoskeletal movement
multicellular organismal movement
actomyosin structure organization
actin-myosin filament sliding
Tuberculosis
negative regulation of leukocyte activation
cellular respiration
defense response to bacterium
The citric acid (TCA) cycle and respiratory electron transport
negative regulation of cell activation
Electron transport chain
Phagosome
Staphylococcus aureus infection
Antigen processing and presentation
term_name
BAT
HEART
LIVER
LUNG
SKM_GN
Tissue
BH_adj_p_value
0.04
0.03
0.02
0.01
recall
0.1
0.2
0.3
0.4
0.5

Supplemental Figure S17: Per tissue pathway enrichment analysis of DEGs withDARs located within 500 kb. Top 10 enriched, non-redundant pathways are shown for each tissue.

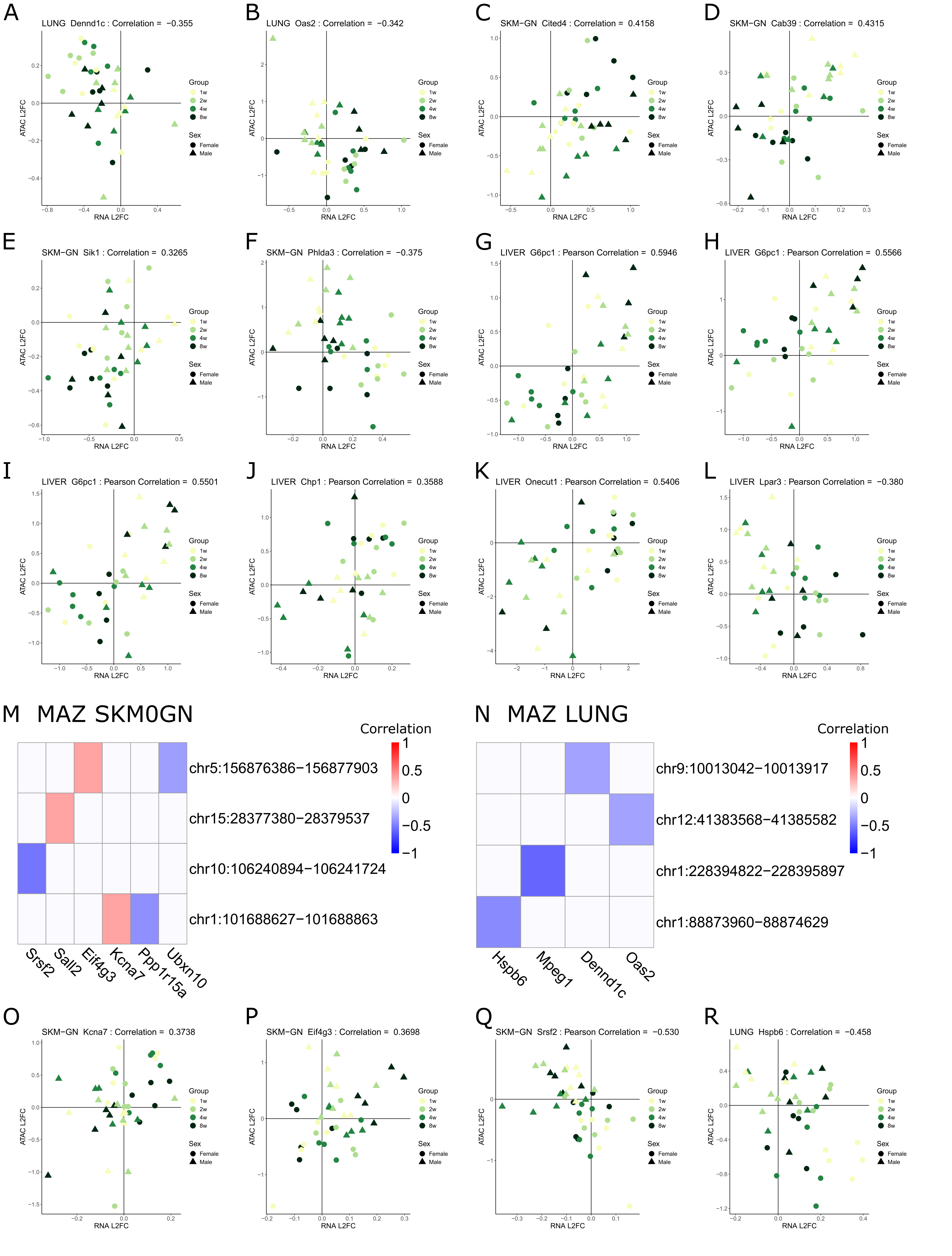
A
LUNG Dennd1c : Correlation = −0.355
B
LUNG Oas2 : Correlation = −0.342
C
SKM-GN Cited4 : Correlation = 0.4158
D
SKM-GN Cab39 : Correlation = 0.4315
E
SKM-GN Sik1 : Correlation = 0.3265
F
SKM-GN Phlda3 : Correlation = −0.375
G
LIVER G6pc1 : Pearson Correlation = 0.5946
H
LIVER G6pc1 : Pearson Correlation = 0.5566
I
LIVER G6pc1 : Pearson Correlation = 0.5501
J
LIVER Chp1 : Pearson Correlation = 0.3588
K
LIVER Onecut1 : Pearson Correlation = 0.5406
L
LIVER Lpar3 : Pearson Correlation = −0.380
ATAC L2FC
RNA L2FC
Group
1w
2w
4w
8w
Sex
Female
Male
M MAZ SKM0GN
Correlation
1
0.5
0
−0.5
−1
chr5:156876386−156877903
chr15:28377380−28379537
chr10:106240894−106241724
chr1:101688627−101688863
Srsf2
Sall2
Eif4g3
Kcna7
Ppp1r15a
Ubxn10
N MAZ LUNG
Correlation
chr9:10013042−10013917
chr12:41383568−41385582
chr1:228394822−228395897
chr1:88873960−88874629
Hspb6
Mpeg1
Dennd1c
Oas2
O
SKM-GN Kcna7 : Correlation = 0.3738
P
SKM-GN Eif4g3 : Correlation = 0.3698
Q
SKM-GN Srsf2 : Pearson Correlation = −0.530
R
LUNG Hspb6 : Correlation = −0.458

Supplemental Figure S18: (a-l) Training response L2FC scatter plots of correlated DAR-DEG examples. (a-b) SP2-target DARs in lung correlated with *Dennd1c* (a), *Oas2* (b); (c-f) BMYB-target DARs in SKM-GN correlated with *Cited4* (c), *Cab39* (d), *Sik1* (e), *Phlda3* (f); (g-l) BMAL1-target DARs in liver correlated with *G6pc1* (g-i), *Chp1* (j), *Onecut1* (k), *Lpar3* (l). (m-n) DAR-DEG pairs containing the binding motif of MAZ in SKM-GN (m), and lung (n). (o-q) MAZ-target DARs in SKM-GN correlated with *Kcna7* (o), *Eif4g3* (p), *Srsf2* (q); (r) MAZ-target DAR in lung correlated with *Hspb6*.

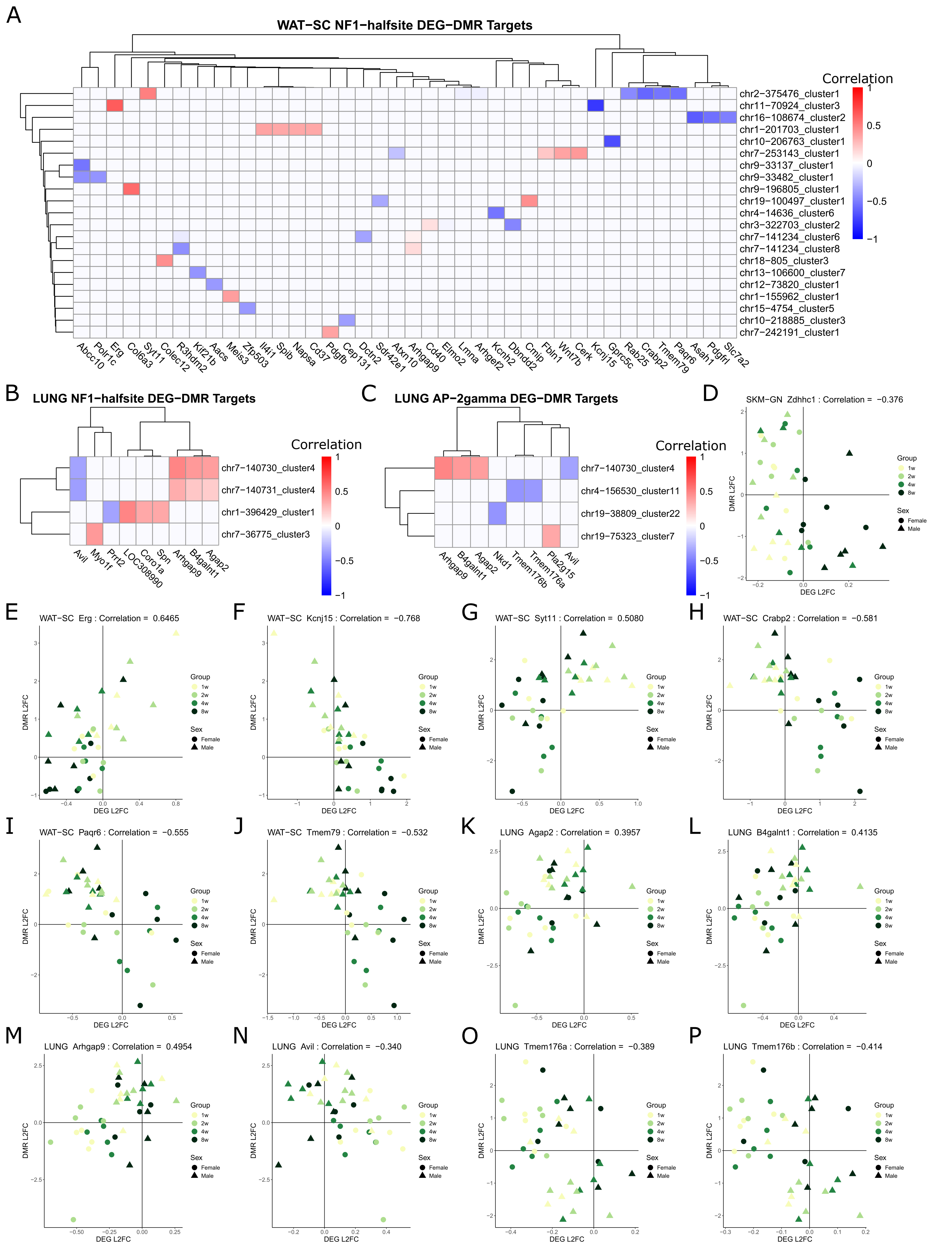
A
WAT−SC NF1−halfsite DEG−DMR Targets
Correlation
1
0.5
0
−0.5
−1
chr2−375476_cluster1
chr11−70924_cluster3
chr16−108674_cluster2
chr1−201703_cluster1
chr10−206763_cluster1
chr7−253143_cluster1
chr9−33137_cluster1
chr9−33482_cluster1
chr9−196805_cluster1
chr19−100497_cluster1
chr4−14636_cluster6
chr3−322703_cluster2
chr7−141234_cluster6
chr7−141234_cluster8
chr18−805_cluster3
chr13−106600_cluster7
chr12−73820_cluster1
chr1−155962_cluster1
chr15−4754_cluster5
chr10−218885_cluster3
chr7−242191_cluster1
Abcc10 Polr1c Erg Col6a3 Syt11 Colec12 R3hdm2 Kif21b Aacs Meis3 Zfp503 Il4i1 Spib Napsa Cd37 Pdgfb Cep131 Dctn2 Sdr42e1 Atxn10 Arhgap9 Cd40 Elmo2 Lmna Arhgef2 Kcnh2 Dbndd2 Cmip Fbln1 Wnt7b Cerk Kcnj15 Gprc5c Rab25 Crabp2 Tmem79 Paqr6 Asah1 Pdgfrl Slc7a2
B
LUNG NF1−halfsite DEG−DMR Targets
Correlation
chr7−140730_cluster4
chr7−140731_cluster4
chr1−396429_cluster1
chr7−36775_cluster3
Avil Myo1f Prrt2 LOC308990 Coro1a Spn Arhgap9 B4galnt1 Agap2
C
LUNG AP−2gamma DEG−DMR Targets
Correlation
chr7−140730_cluster4
chr4−156530_cluster11
chr19−38809_cluster22
chr19−75323_cluster7
Arhgap9 B4galnt1 Agap2 Nkd1 Tmem176b Tmem176a Pla2g15 Avil
D
SKM−GN Zdhhc1 : Correlation = −0.376
E
WAT−SC Erg : Correlation = 0.6465
F
WAT−SC Kcnj15 : Correlation = −0.768
G
WAT−SC Syt11 : Correlation = 0.5080
H
WAT−SC Crabp2 : Correlation = −0.581
I
WAT−SC Paqr6 : Correlation = −0.555
J
WAT−SC Tmem79 : Correlation = −0.532
K
LUNG Agap2 : Correlation = 0.3957
L
LUNG B4galnt1 : Correlation = 0.4135
M
LUNG Arhgap9 : Correlation = 0.4954
N
LUNG Avil : Correlation = −0.340
O
LUNG Tmem176a : Correlation = −0.389
P
LUNG Tmem176b : Correlation = −0.414
DMR L2FC
DEG L2FC
Group
1w
2w
4w
8w
Sex
Female
Male

Supplemental Figure S19: (a-b) TF NF1-halfsite correlated DMR-DEG targets in WAT-SC (a), and lung (b). (c) TF AP-2gamma correlated DMR-DEG targets in lung. (d-p) Training response L2FC scatter plots of correlated DMR-DEG examples. (d) SP2-target DMR in SKM-GN correlated with *Zdhhc1*. (e-j) NF1-halfsite-target DMRs in WAT-SC correlated with *Erg* (e), *Kcnj15* (f), *Syt11* (g), *Crabp2* (h), *Paqr6* (i), *Tmem79* (j); (k-n) NF1-halfsite and AP-2gamma targeted DMRs in lung correlated with *Agap2* (k), *B4galnt1* (l), *Arhgap9* (m), *Avil* (n); (o-p) AP-2gamma-target DMRs in lung correlated with *Tmem176a* (o), *Tmem176b* (p).

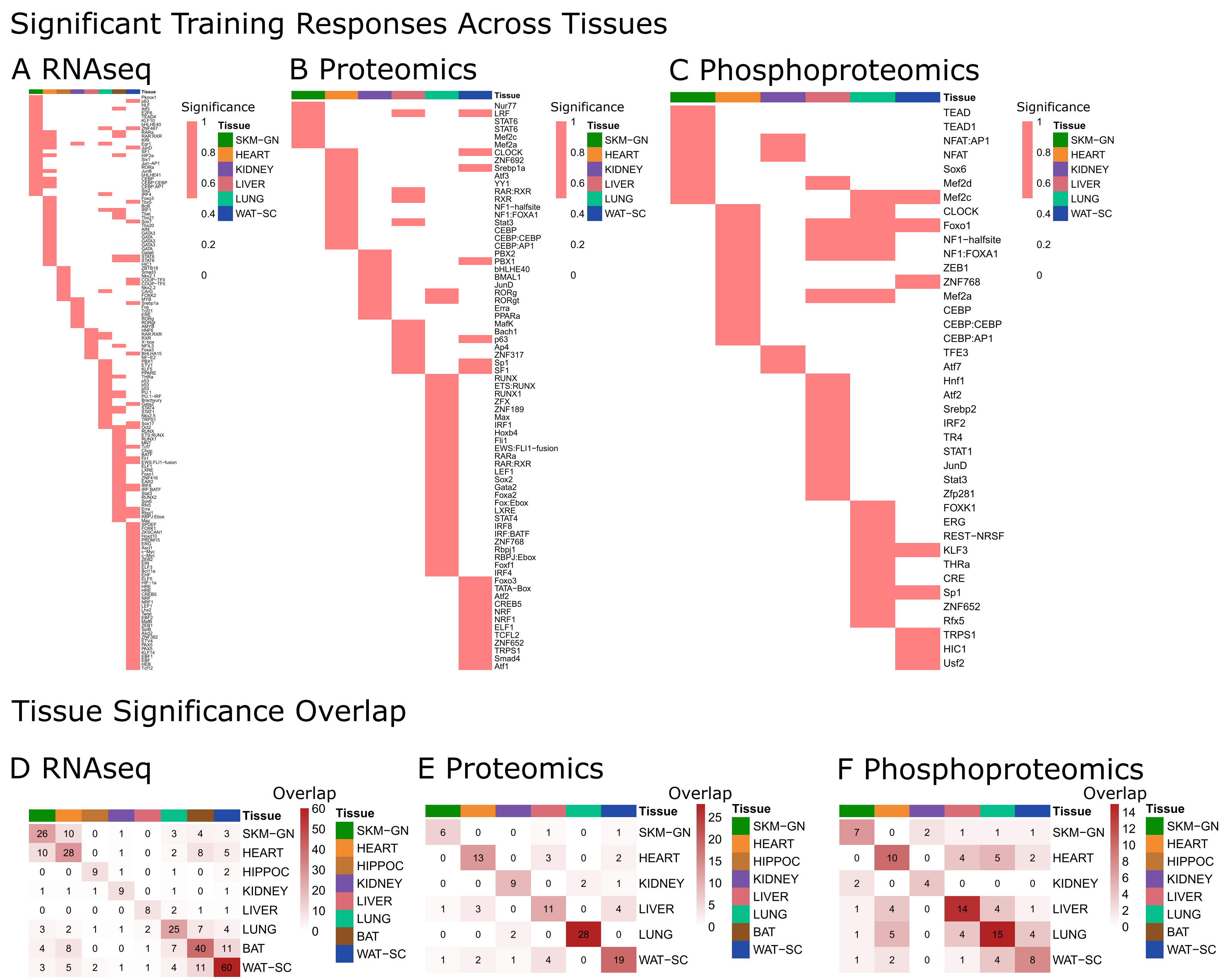

Significant Training Responses Across Tissues
A RNAseq
B Proteomics
C Phosphoproteomics
Significance
Tissue
SKM−GN
HEART
KIDNEY
LIVER
LUNG
WAT−SC
Nur77
LRF
STAT6
STAT6
Mef2c
Mef2a
CLOCK
ZNF692
Srebp1a
Atf3
YY1
RAR:RXR
RXR
NF1−halfsite
NF1:FOXA1
Stat3
CEBP
CEBP:CEBP
CEBP:AP1
PBX2
PBX1
bHLHE40
BMAL1
JunD
RORg
RORgt
Erra
PPARa
MafK
Bach1
p63
Ap4
ZNF317
Sp1
SF1
RUNX
ETS:RUNX
RUNX1
ZFX
ZNF189
Max
IRF1
Hoxb4
Fli1
EWS:FLI1−fusion
RARa
RAR:RXR
LEF1
Sox2
Gata2
Foxa2
Fox:Ebox
LXRE
STAT4
IRF8
IRF:BATF
ZNF768
Rbpj1
RBPJ:Ebox
Foxf1
IRF4
Foxo3
TATA−Box
Atf2
CREB5
NRF
NRF1
ELF1
TCFL2
ZNF652
TRPS1
Smad4
Atf1
TEAD
TEAD1
NFAT:AP1
NFAT
Sox6
Mef2d
Mef2c
CLOCK
Foxo1
NF1−halfsite
NF1:FOXA1
ZEB1
ZNF768
Mef2a
CEBP
CEBP:CEBP
CEBP:AP1
TFE3
Atf7
Hnf1
Atf2
Srebp2
IRF2
TR4
STAT1
JunD
Stat3
Zfp281
FOXK1
ERG
REST−NRSF
KLF3
THRa
CRE
Sp1
ZNF652
Rfx5
TRPS1
HIC1
Usf2
Tissue Significance Overlap
D RNAseq
E Proteomics
F Phosphoproteomics
Overlap
Tissue
SKM−GN
HEART
HIPPOC
KIDNEY
LIVER
LUNG
BAT
WAT−SC
26 10 0 1 0 3 4 3 SKM−GN
10 28 0 1 0 2 8 5 HEART
0 0 9 1 0 1 0 2 HIPPOC
1 1 1 9 0 1 0 1 KIDNEY
0 0 0 0 8 2 1 1 LIVER
3 2 1 1 2 25 7 4 LUNG
4 8 0 0 1 7 40 11 BAT
3 5 2 1 1 4 11 60 WAT−SC
6 0 0 1 0 1 SKM−GN
0 13 0 3 0 2 HEART
0 0 9 0 2 1 KIDNEY
1 3 0 11 0 4 LIVER
0 0 2 0 28 0 LUNG
1 2 1 4 0 19 WAT−SC
7 0 2 1 1 1 SKM−GN
0 10 0 4 5 2 HEART
2 0 4 0 0 0 KIDNEY
1 4 0 14 4 1 LIVER
1 5 0 4 15 4 LUNG
1 2 0 1 4 8 WAT−SC

Supplemental Figure S20: Identifying cross-tissue conserved differential signal in TFs. (a-c) Heatmaps of TFs displaying significance at the RNA-seq (a), proteome (b), and phosphoproteome (c) level. In each heatmap, a grid square is red if the TF is significant in that ome in that tissue. (d-f) Quantifying the overlap of significant TFs at the RNA-seq (d), proteome (e), and phosphoproteome (f) level.

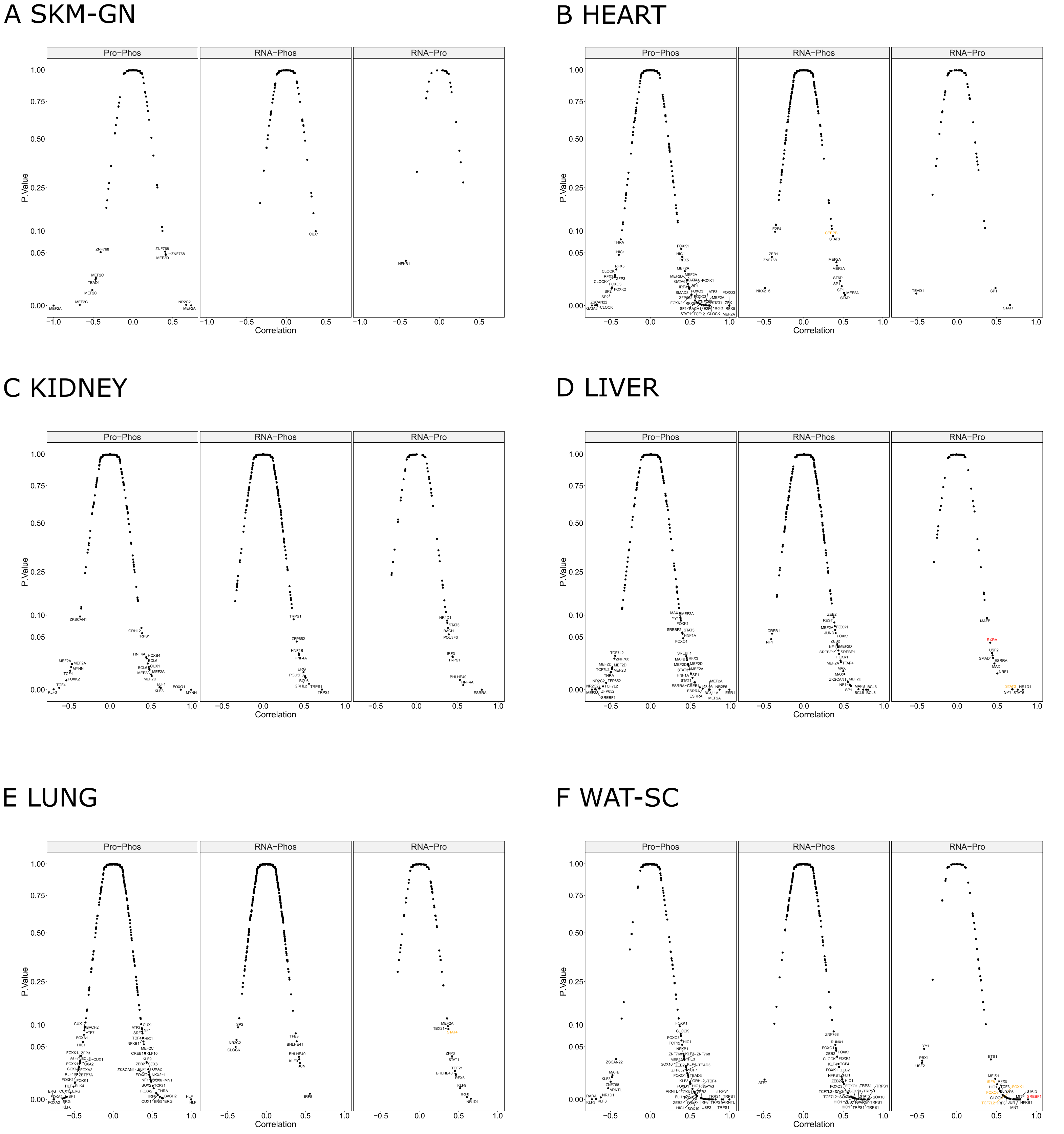
A SKM-GN
B HEART
C KIDNEY
D LIVER
E LUNG
F WAT-SC
Pro−Phos
RNA−Phos
RNA−Pro
P.Value
Correlation
1.00
0.75
0.50
0.25
0.10
0.05
0.00
−1.0
−0.5
0.0
0.5
1.0

Supplemental Figure S21: Scatter plots of magnitude (x-axis) and significance (y-axis) of correlation in EET response for TFs across omes and tissues. Correlation is measured for EET response between protein abundance and protein phosphorylation, gene expression and protein phosphorylation and gene expression and protein abundance for each tissue: skeletal muscle (a), heart (b), kidney (c), liver (d), lung (e), white adipose (f). Names of TFs on the scatter plots are colored orange if the TF has a significant (adjusted p.val < 0.1) EET response in one of the two compared omes, and colored red if the TF has a significant EET response in both compared omes.

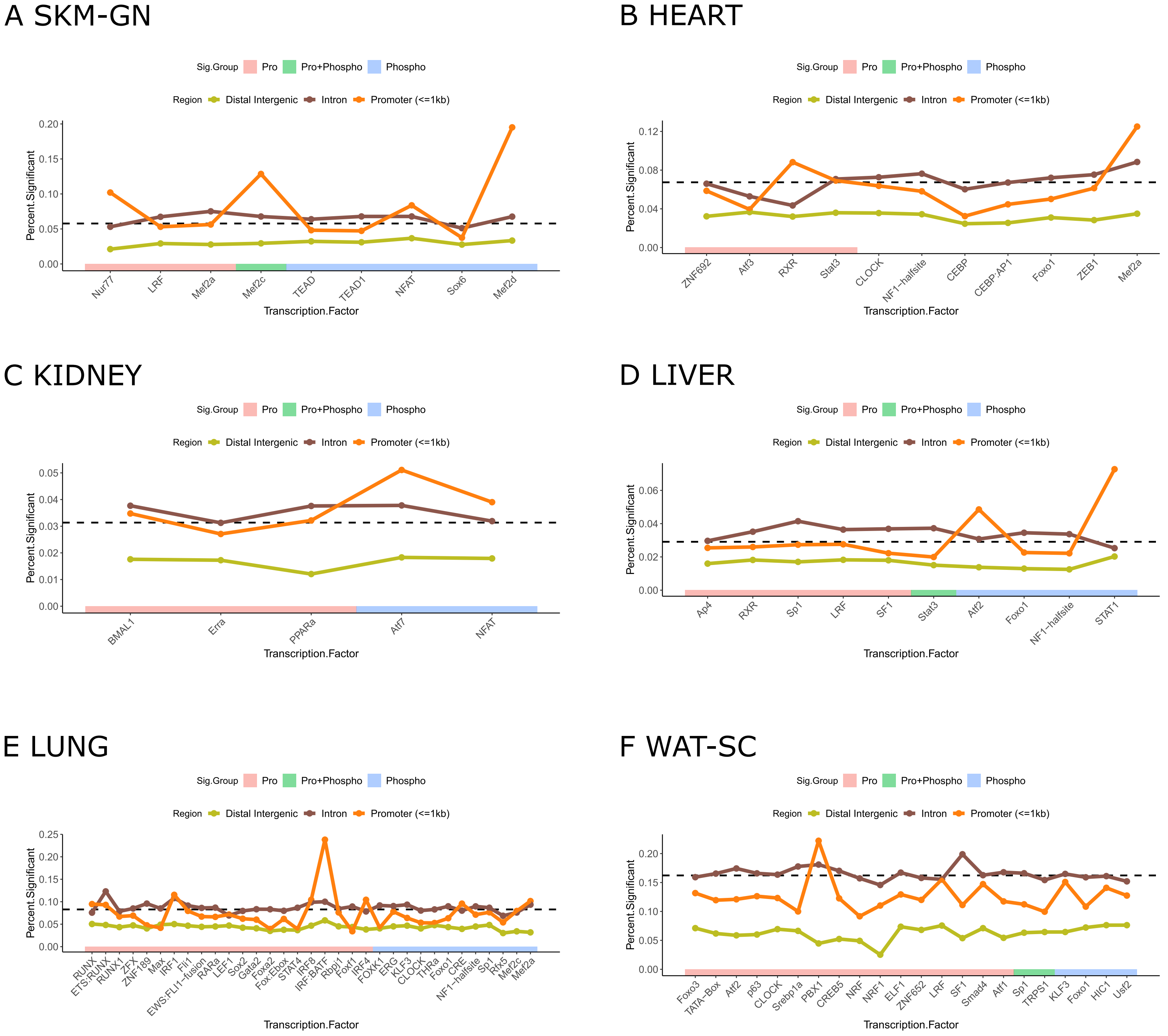
A SKM-GN
Sig.Group Pro Pro+Phospho Phospho
Region Distal Intergenic Intron Promoter (<=1kb)
Percent.Significant
0.00 0.05 0.10 0.15 0.20
Nur77 LRF Mef2a Mef2c TEAD TEAD1 NFAT Sox6 Mef2d
Transcription.Factor
B HEART
Sig.Group Pro Pro+Phospho Phospho
Region Distal Intergenic Intron Promoter (<=1kb)
Percent.Significant
0.00 0.04 0.08 0.12
ZNF692 Atf3 RXR Stat3 CLOCK NF1−halfsite CEBP CEBP:AP1 Foxo1 ZEB1 Mef2a
Transcription.Factor
C KIDNEY
Sig.Group Pro Pro+Phospho Phospho
Region Distal Intergenic Intron Promoter (<=1kb)
Percent.Significant
0.00 0.01 0.02 0.03 0.04 0.05
BMAL1 Erra PPARa Atf7 NFAT
Transcription.Factor
D LIVER
Sig.Group Pro Pro+Phospho Phospho
Region Distal Intergenic Intron Promoter (<=1kb)
Percent.Significant
0.00 0.02 0.04 0.06
Ap4 RXR Sp1 LRF SF1 Stat3 Atf2 Foxo1 NF1−halfsite STAT1
Transcription.Factor
E LUNG
Sig.Group Pro Pro+Phospho Phospho
Region Distal Intergenic Intron Promoter (<=1kb)
Percent.Significant
0.00 0.05 0.10 0.15 0.20 0.25
RUNX ETS:RUNX RUNX1 ZFX ZNF189 Max IRF1 Fli1 EWS:FLI1−fusion RARa LEF1 Sox2 Gata2 Foxa2 Fox:Ebox STAT4 IRF8 IRF:BATF Rbpj1 Foxf1 IRF4 FOXK1 ERG KLF3 CLOCK THRa Foxo1 CRE NF1−halfsite Sp1 Rfx5 Mef2c Mef2a
Transcription.Factor
F WAT-SC
Sig.Group Pro Pro+Phospho Phospho
Region Distal Intergenic Intron Promoter (<=1kb)
Percent.Significant
0.00 0.05 0.10 0.15 0.20
Foxo3 TATA−Box Atf2 p63 CLOCK Srebp1a PBX1 CREB5 NRF NRF1 ELF1 ZNF652 LRF SF1 Smad4 Atf1 Sp1 TRPS1 KLF3 Foxo1 HIC1 Usf2
Transcription.Factor

Supplemental Figure S22: Frequency of DEGs among gene targets of TFs with significant proteomic or phosphoproteomic training response in (a) skeletal muscle, (b) heart, (c) kidney, (d) liver, (e) lung, (f) white adipose. TFs are color-annotated by the ome of their training response: protein (red), phosphoprotein (blue) or both (green). The dashed line represents the frequency of DEGs among active genes in each tissue. Gene targets are divided by presence of TF motif within active peaks annotated to the proximal promoter (orange), intron (brown) or distal intergenic (gold) features.

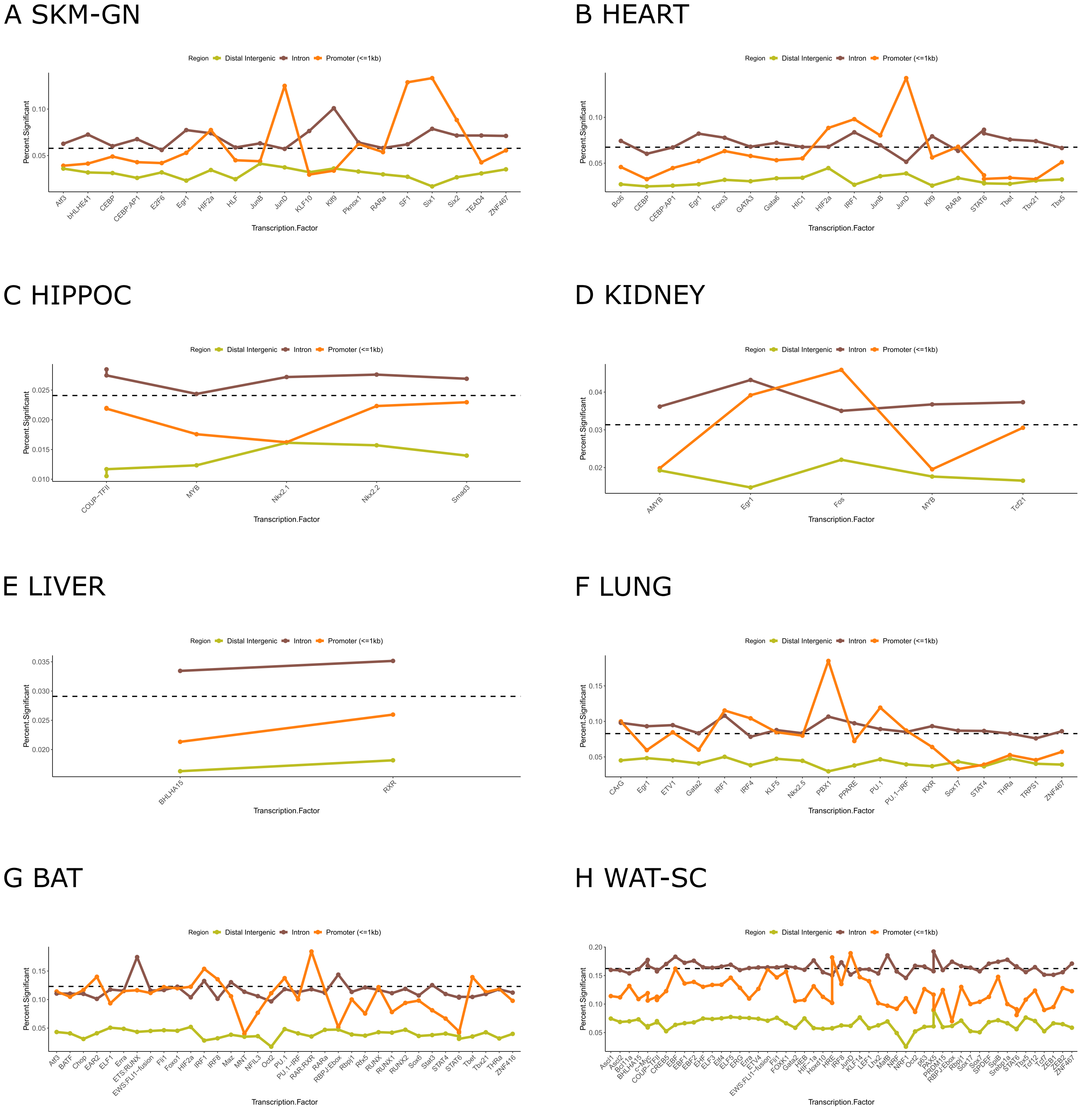
A SKM-GN
Region Distal Intergenic Intron Promoter (<=1kb)
Percent.Significant
0.05
0.10
Atf3 bHLHE41 CEBP CEBP:AP1 E2F6 Egr1 HIF2a HLF JunB JunD KLF10 Klf9 Pknox1 RARa SF1 Six1 Six2 TEAD4 ZNF467
Transcription.Factor
B HEART
Region Distal Intergenic Intron Promoter (<=1kb)
Percent.Significant
0.05
0.10
Bcl6 CEBP CEBP:AP1 Egr1 Foxo3 GATA3 Gata6 HIC1 HIF2a IRF1 JunB JunD Klf9 RARa STAT6 Tbet Tbx21 Tbx5
Transcription.Factor
C HIPPOC
Region Distal Intergenic Intron Promoter (<=1kb)
Percent.Significant
0.010
0.015
0.020
0.025
COUP−TFII MYB Nkx2.1 Nkx2.2 Smad3
Transcription.Factor
D KIDNEY
Region Distal Intergenic Intron Promoter (<=1kb)
Percent.Significant
0.02
0.03
0.04
AMYB Egr1 Fos MYB Tcf21
Transcription.Factor
E LIVER
Region Distal Intergenic Intron Promoter (<=1kb)
Percent.Significant
0.020
0.025
0.030
0.035
BHLHA15 RXR
Transcription.Factor
F LUNG
Region Distal Intergenic Intron Promoter (<=1kb)
Percent.Significant
0.05
0.10
0.15
CArG Egr1 ETV1 Gata2 IRF1 IRF4 KLF5 Nkx2.5 PBX1 PPARE PU.1 PU.1−IRF RXR Sox17 STAT4 THRa TRPS1 ZNF467
Transcription.Factor
G BAT
Region Distal Intergenic Intron Promoter (<=1kb)
Percent.Significant
0.05
0.10
0.15
Atf3 BATF Chop EAR2 ELF1 Erra ETS:RUNX EWS:FLI1−fusion Fli1 Foxo1 HIF2a IRF1 IRF8 Maz MNT NFIL3 Oct2 PU.1 PU.1−IRF RAR:RXR RARa RBPJ:Ebox Rbpj1 Rfx5 RUNX RUNX1 RUNX2 Sox6 Stat3 STAT4 STAT6 Tbet Tbx21 THRa ZNF416
Transcription.Factor
H WAT-SC
Region Distal Intergenic Intron Promoter (<=1kb)
Percent.Significant
0.05
0.10
0.15
0.20
Transcription.Factor

Supplemental Figure S23: Frequency of DEGs among gene targets of TFs with significant transcriptomic  training response in (a) skeletal muscle, (b) heart, (c) hippocampus, (d) kidney, (e) liver, (f) lung, (g) brown adipose, and (f) white adipose. The dashed line represents the frequency of DEGs among active genes in each tissue. Gene targets are divided by presence of TF motif within active peaks annotated to the proximal promoter (orange), intron (brown) or distal intergenic (gold) features.

### TFs with Significant Changes in Gene Expression

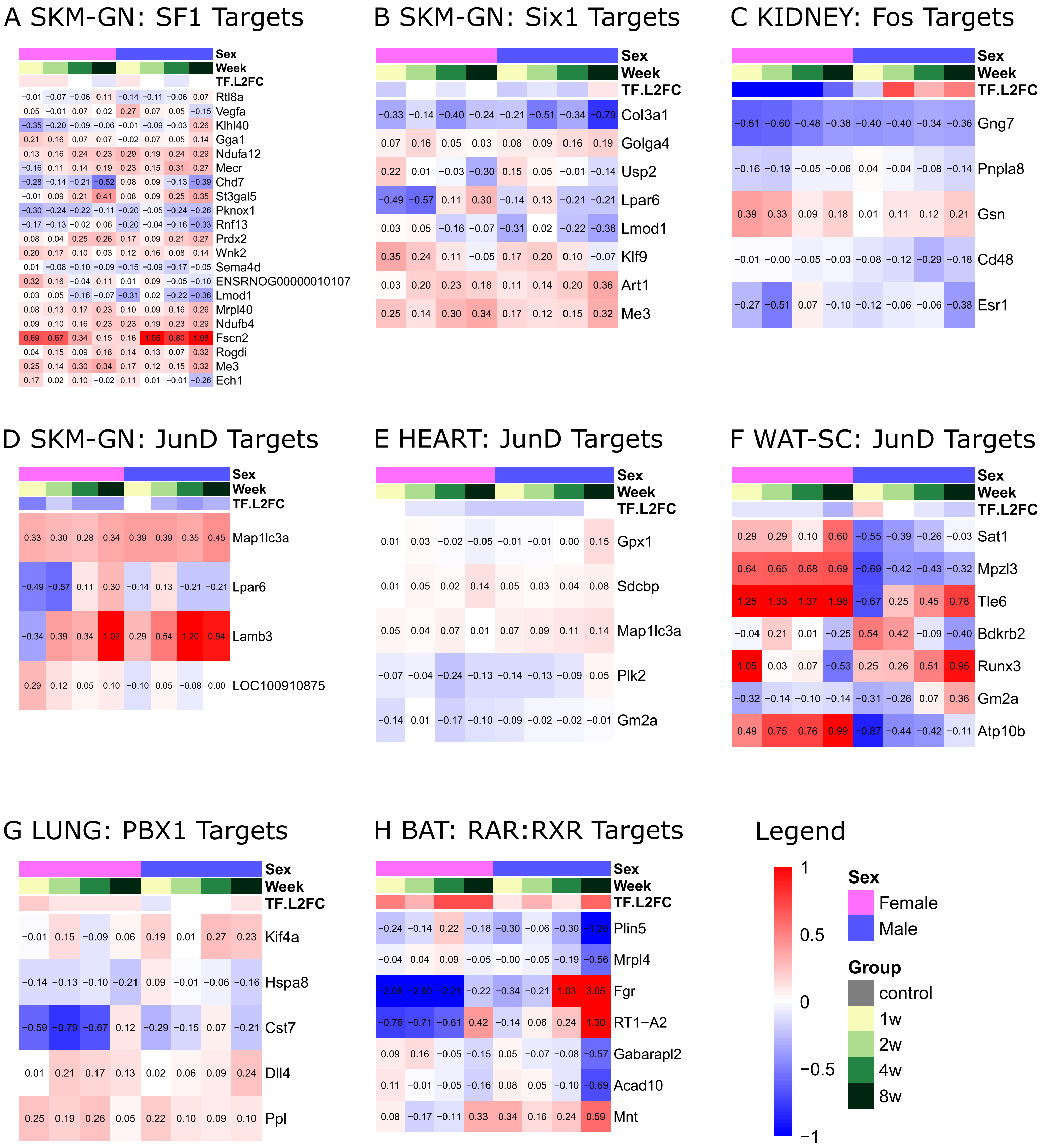

Supplemental Figure S24: Heatmaps of EET response for DEGs witih proximal promoter motifs of TFs with significant changes at the RNA-seq level including (a) SF1, (b) Six1 in SKM-GN, (c) Fos in kidney, (d) JunD in SKM-GN, (e) JunD in heart, (f) JunD in WAT-SC, (g) PBX1 in lung, (h) RAR:RXR in BAT.

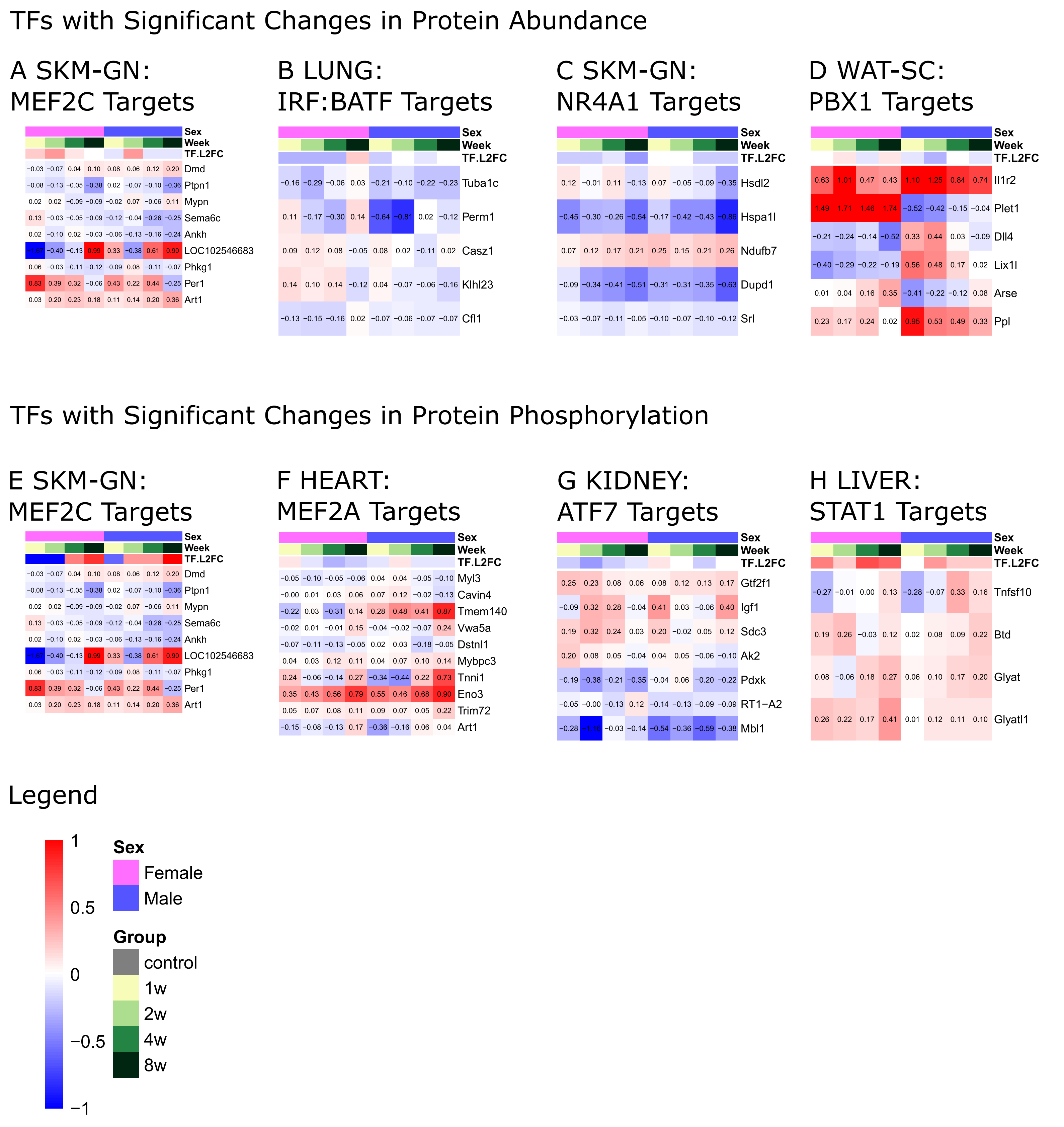
TFs with Significant Changes in Protein Abundance
A SKM-GN: MEF2C Targets
B LUNG: IRF:BATF Targets
C SKM-GN: NR4A1 Targets
D WAT-SC: PBX1 Targets
Sex
Week
TF.L2FC
Dmd
Ptpn1
Mypn
Sema6c
Ankh
LOC102546683
Phkg1
Per1
Art1
Tuba1c
Perm1
Casz1
Klhl23
Cfl1
Hsdl2
Hspa1l
Ndufb7
Dupd1
Srl
Il1r2
Plet1
Dll4
Lix1l
Arse
Ppl
TFs with Significant Changes in Protein Phosphorylation
E SKM-GN: MEF2C Targets
F HEART: MEF2A Targets
G KIDNEY: ATF7 Targets
H LIVER: STAT1 Targets
Myl3
Cavin4
Tmem140
Vwa5a
Dstnl1
Mybpc3
Tnni1
Eno3
Trim72
Art1
Gtf2f1
Igf1
Sdc3
Ak2
Pdxk
RT1−A2
Mbl1
Tnfsf10
Btd
Glyat
Glyatl1
Legend
1
0.5
0
−0.5
−1
Sex
Female
Male
Group
control
1w
2w
4w
8w

Supplemental Figure S25: Heatmaps of EET response for DEGs with proximal promoter motifs of TFs with significant changes at the proteomic (a-d) or phosphoproteomic (e-h) level including (a) MEF2C in SKM-GN, (b) IRF:BATF in lung, (c) NR4A1 in SKM-GN, (d) PBX1 in WAT-SC among proteomic TF changes, and (e) MEF2C in SKM-GN, (f) MEF2A in heart, (g) ATF7 in kidney, and (h) STAT1 in liver among phosphoproteomic TF changes.

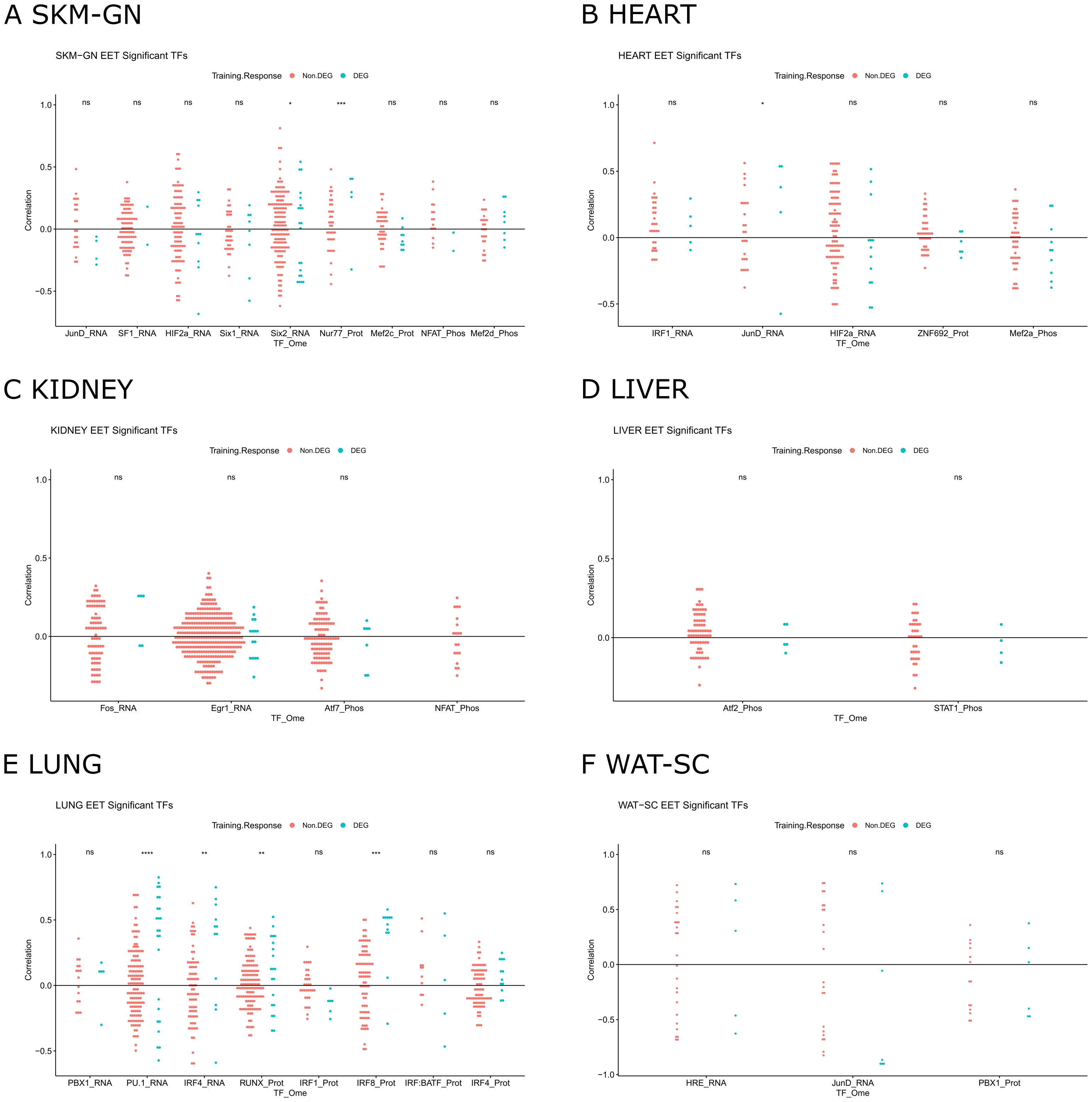
A SKM-GN
SKM−GN EET Significant TFs
Training.Response Non.DEG DEG
ns ns ns ns * *** ns ns ns
Correlation
1.0
0.5
0.0
−0.5
JunD_RNA SF1_RNA HIF2a_RNA Six1_RNA Six2_RNA Nur77_Prot Mef2c_Prot NFAT_Phos Mef2d_Phos
TF_Ome
B HEART
HEART EET Significant TFs
Training.Response Non.DEG DEG
ns * ns ns ns
Correlation
1.0
0.5
0.0
−0.5
IRF1_RNA JunD_RNA HIF2a_RNA ZNF692_Prot Mef2a_Phos
TF_Ome
C KIDNEY
KIDNEY EET Significant TFs
Training.Response Non.DEG DEG
ns ns ns
Correlation
1.0
0.5
0.0
Fos_RNA Egr1_RNA Atf7_Phos NFAT_Phos
TF_Ome
D LIVER
LIVER EET Significant TFs
Training.Response Non.DEG DEG
ns ns
Correlation
1.0
0.5
0.0
Atf2_Phos STAT1_Phos
TF_Ome
E LUNG
LUNG EET Significant TFs
Training.Response Non.DEG DEG
ns **** ** ** ns *** ns ns
Correlation
1.0
0.5
0.0
−0.5
PBX1_RNA PU.1_RNA IRF4_RNA RUNX_Prot IRF1_Prot IRF8_Prot IRF:BATF_Prot IRF4_Prot
TF_Ome
F WAT-SC
WAT−SC EET Significant TFs
Training.Response Non.DEG DEG
ns ns ns
Correlation
1.0
0.5
0.0
−0.5
−1.0
HRE_RNA JunD_RNA PBX1_Prot
TF_Ome

Supplemental Figure S26: Dotplots measuring correlation between TF response to EET and proximal promoter targets' response to EET for select TFs with significant training responses at the RNA, proteome or phosphoproteome levels in skeletal muscle (a), heart (b), kidney (c), liver (d), lung (e), white adipose (f). Target genes are divided into non-DEGs (red) and DEGs (blue). * represents $p < 0.05$ of t test for difference between correlation of TF with DEG targets and non-DEG targets.

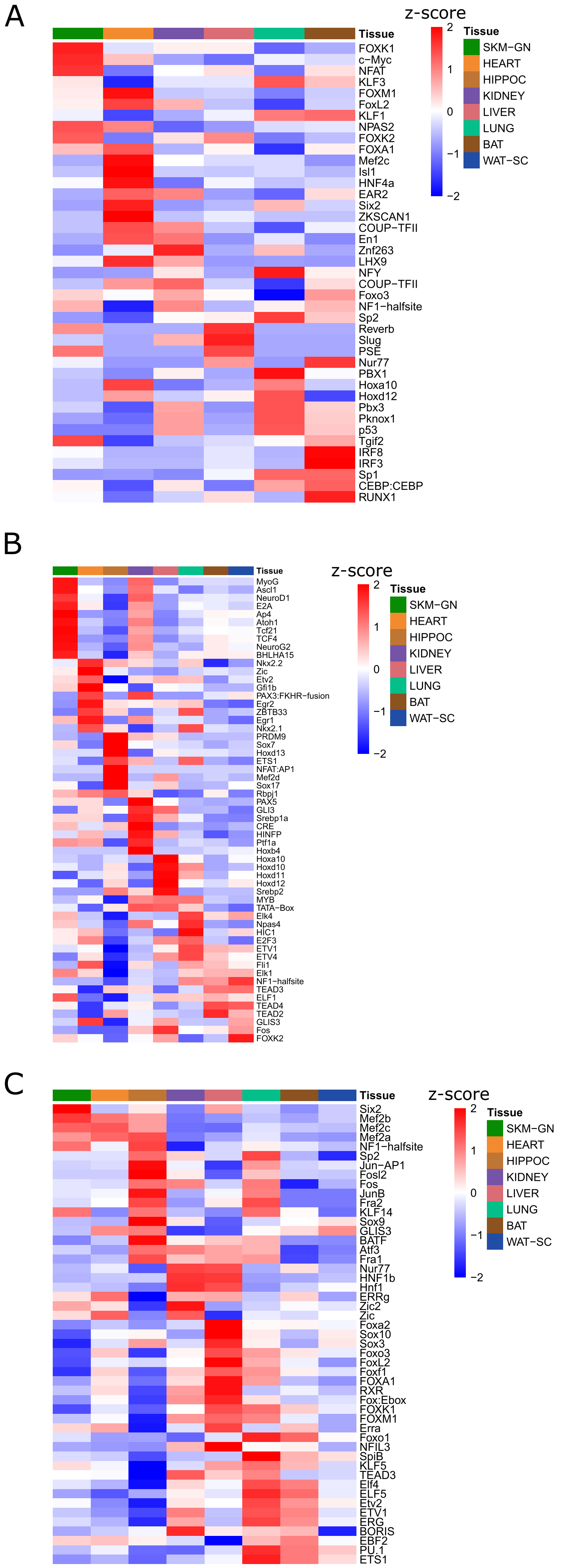
A
Tissue
z-score
2
1
0
−1
−2
Tissue
SKM−GN
HEART
HIPPOC
KIDNEY
LIVER
LUNG
BAT
WAT−SC
FOXK1
c−Myc
NFAT
KLF3
FOXM1
FoxL2
KLF1
NPAS2
FOXK2
FOXA1
Mef2c
Isl1
HNF4a
EAR2
Six2
ZKSCAN1
COUP−TFII
En1
Znf263
LHX9
NFY
COUP−TFII
Foxo3
NF1−halfsite
Sp2
Reverb
Slug
PSE
Nur77
PBX1
Hoxa10
Hoxd12
Pbx3
Pknox1
p53
Tgif2
IRF8
IRF3
Sp1
CEBP:CEBP
RUNX1
B
Tissue
z-score
2
1
0
−1
−2
Tissue
SKM−GN
HEART
HIPPOC
KIDNEY
LIVER
LUNG
BAT
WAT−SC
MyoG
Ascl1
NeuroD1
E2A
Ap4
Atoh1
Tcf21
TCF4
NeuroG2
BHLHA15
Nkx2.2
Zic
Etv2
Gfi1b
PAX3:FKHR−fusion
Egr2
ZBTB33
Egr1
Nkx2.1
PRDM9
Sox7
Hoxd13
ETS1
NFAT:AP1
Mef2d
Sox17
Rbpj1
PAX5
GLI3
Srebp1a
CRE
HINFP
Ptf1a
Hoxb4
Hoxa10
Hoxd10
Hoxd11
Hoxd12
Srebp2
MYB
TATA−Box
Elk4
Npas4
HIC1
E2F3
ETV1
ETV4
Fli1
Elk1
NF1−halfsite
TEAD3
ELF1
TEAD4
TEAD2
GLIS3
Fos
FOXK2
C
Tissue
z-score
2
1
0
−1
−2
Tissue
SKM−GN
HEART
HIPPOC
KIDNEY
LIVER
LUNG
BAT
WAT−SC
Six2
Mef2b
Mef2c
Mef2a
NF1−halfsite
Sp2
Jun−AP1
Fosl2
Fos
JunB
Fra2
KLF14
Sox9
GLIS3
BATF
Atf3
Fra1
Nur77
HNF1b
Hnf1
ERRg
Zic2
Zic
Foxa2
Sox10
Sox3
Foxo3
FoxL2
Foxf1
FOXA1
RXR
Fox:Ebox
FOXK1
FOXM1
Erra
Foxo1
NFIL3
SpiB
KLF5
TEAD3
Elf4
ELF5
Etv2
ETV1
ERG
BORIS
EBF2
PU.1
ETS1

Supplemental Figure S27: (a) z-scores of TF motif frequency among DARs across tissues for top DAR-enriched TFs. (b) z-scores of TF motif frequency among DMRs across tissues for top DMR-enriched TFs. (c) z-scores of TF motif frequency among DEGaPs across tissues for top DEGaP-enriched TFs.

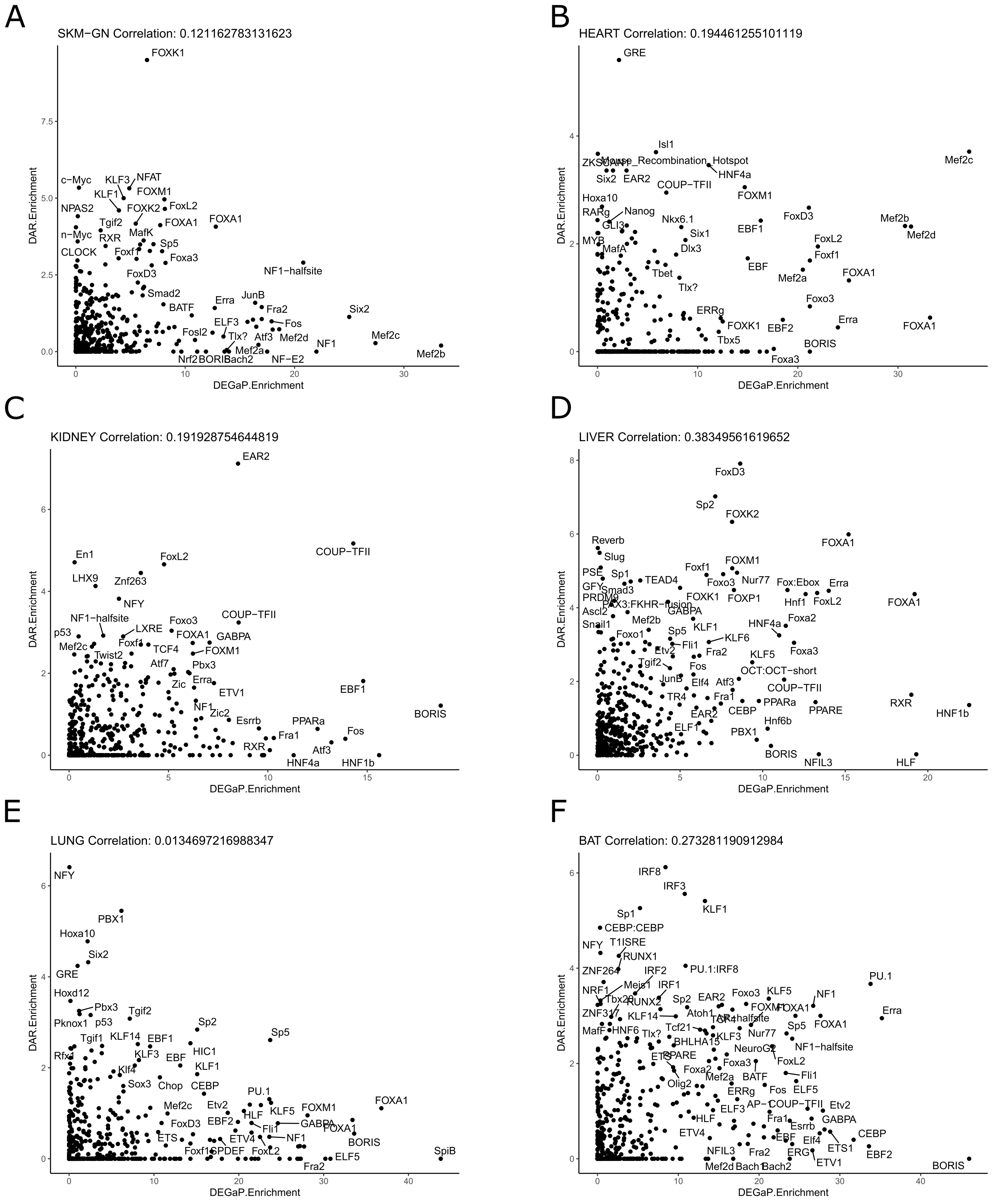
A
SKM−GN Correlation: 0.121162783131623
DAR.Enrichment
DEGaP.Enrichment
FOXK1
c−Myc
KLF3
NFAT
KLF1
FOXM1
NPAS2
FOXK2
FoxL2
Tgif2
FOXA1
FOXA1
n−Myc
MafK
RXR
Foxf1
Sp5
CLOCK
Foxa3
NF1−halfsite
FoxD3
Smad2
JunB
Erra
BATF
Fra2
Six2
ELF3
Fos
Fosl2
Tlx?
Atf3
Mef2d
Mef2c
NF1
Mef2a
Nrf2
BORIS
Bach2
NF−E2
Mef2b
B
HEART Correlation: 0.194461255101119
GRE
Isl1
ZKSCAN
Mouse_Recombination_Hotspot
HNF4a
Mef2c
Six2
EAR2
COUP−TFII
FOXM1
Hoxa10
Nanog
FoxD3
RARg
Nkx6.1
EBF1
Mef2b
GLI3
Six1
Mef2d
MYB
FoxL2
MafA
Dlx3
Foxf1
EBF
Tbet
Mef2a
FOXA1
Tlx?
Foxo3
ERRg
FOXK1
EBF2
Erra
FOXA1
Tbx5
BORIS
Foxa3
C
KIDNEY Correlation: 0.191928754644819
EAR2
COUP−TFII
En1
FoxL2
LHX9
Znf263
NFY
NF1−halfsite
COUP−TFII
Foxo3
p53
LXRE
FOXA1
GABPA
Mef2c
Foxf1
TCF4
FOXM1
Twist2
Atf7
Pbx3
Erra
EBF1
Zic
ETV1
NF1
Zic2
Esrrb
PPARa
BORIS
Fra1
Fos
RXR
Atf3
HNF4a
HNF1b
D
LIVER Correlation: 0.38349561619652
FoxD3
Sp2
FOXK2
FOXA1
Reverb
Slug
FOXM1
PSE
Foxf1
Sp1
Nur77
TEAD4
Foxo3
GFY
Fox:Ebox
Erra
Smad3
Hnf1
FoxL2
PRDM9
FOXK1
FOXP1
PAX3:FKHR−fusion
FOXA1
Ascl2
GABPA
Mef2b
HNF4a
Foxa2
Snail1
KLF1
Foxo1
Sp5
KLF6
Fli1
Etv2
Fra2
Foxa3
KLF5
Tgif2
Fos
OCT:OCT−short
JunB
Atf3
Elf4
COUP−TFII
TR4
Fra1
PPARa
PPARE
RXR
CEBP
HNF1b
EAR2
Hnf6b
ELF1
PBX1
BORIS
NFIL3
HLF
E
LUNG Correlation: 0.0134697216988347
NFY
PBX1
Hoxa10
Six2
GRE
Hoxd12
Pbx3
Tgif2
Pknox1
p53
Sp2
Sp5
Tgif1
KLF14
EBF1
Rfx1
KLF3
EBF
HIC1
Klf4
KLF1
Sox3
Chop
CEBP
PU.1
FOXA1
Mef2c
Etv2
KLF5
FOXM1
HLF
FoxD3
EBF2
GABPA
Fli1
FOXA1
ETS
ETV4
NF1
BORIS
Foxf1
SPDEF
FoxL2
ELF5
SpiB
Fra2
F
BAT Correlation: 0.273281190912984
IRF8
IRF3
KLF1
Sp1
CEBP:CEBP
NFY
T1ISRE
RUNX1
PU.1:IRF8
ZNF264
IRF2
PU.1
IRF1
Meis1
Foxo3
KLF5
NF1
NRF1
Tbx20
Sp2
EAR2
FOXM1
FOXA1
Erra
RUNX2
Atoh1
ZNF317
KLF14
TR4
NF1−halfsite
FOXA1
MafF
HNF6
Tcf21
Sp5
Tlx?
Nur77
KLF3
BHLHA15
ETS
NeuroG2
PPARE
NF1−halfsite
Foxa3
FoxL2
Foxa2
BATF
Fli1
Mef2a
Olig2
Fos
ELF5
ERRg
AP−1
COUP−TFII
Etv2
ELF3
HLF
Fra1
Esrrb
GABPA
ETV4
EBF
Elf4
CEBP
ETS1
NFIL3
Fra2
ERG
EBF2
ETV1
Mef2d
Bach1
Bach2
BORIS

Supplemental Figure S28: Comparing TF enrichment among DARs and DEGaPs. Scatter plots of -log10 p-value of TF motif enrichment among DARs and DEGaPs in six tissues: (a) skeletal muscle, (b) heart, (c) kidney, (d) liver, (e) lung, (f) brown adipose. Pearson correlation is calculated for each tissue.

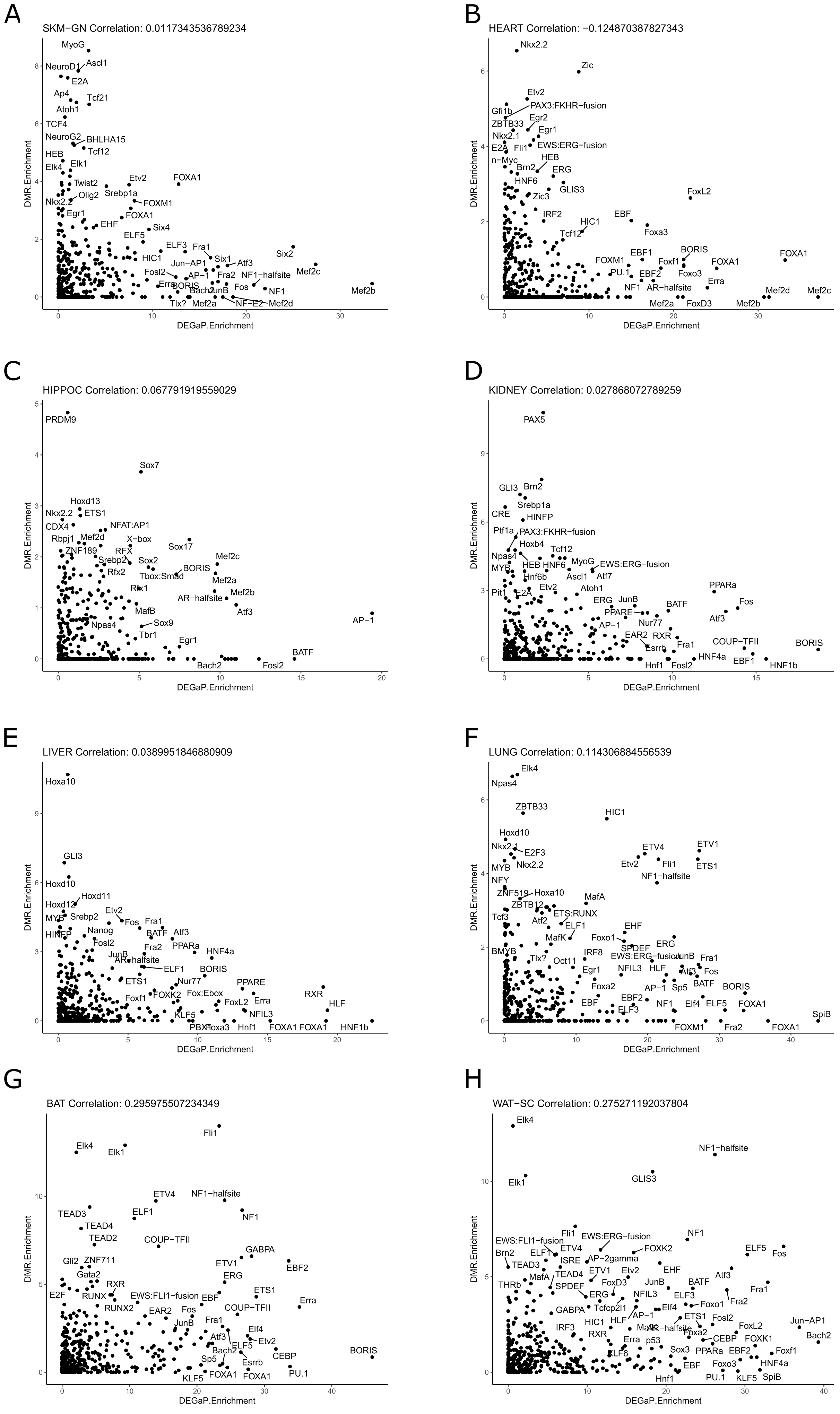
A
SKM−GN Correlation: 0.0117343536789234
MyoG
NeuroD1
Ascl1
E2A
Ap4
Tcf21
Atoh1
TCF4
NeuroG2
BHLHA15
Tcf12
HEB
Elk4
Elk1
Twist2
Etv2
FOXA1
Olig2
Srebp1a
Nkx2.2
FOXM1
Egr1
FOXA1
EHF
Six4
ELF5
ELF3
Fra1
Six2
HIC1
Six1
Atf3
Jun−AP1
Mef2c
Fosl2
AP−1
Fra2
NF1−halfsite
Erra
BORIS
Fos
NF1
Mef2b
Bach2
JunB
Tlx?
Mef2a
NF−E2
Mef2d
DMR.Enrichment
DEGaP.Enrichment
B
HEART Correlation: −0.124870387827343
Nkx2.2
Zic
Etv2
PAX3:FKHR−fusion
Gfi1b
ZBTB33
Egr2
Egr1
Nkx2.1
EWS:ERG−fusion
E2A
Fli1
HEB
n−Myc
Brn2
ERG
HNF6
GLIS3
FoxL2
Zic3
IRF2
EBF
HIC1
Tcf12
Foxa3
EBF1
BORIS
FOXA1
FOXM1
Foxf1
FOXA1
PU.1
Foxo3
EBF2
NF1
AR−halfsite
Erra
Mef2d
Mef2c
Mef2a
FoxD3
Mef2b
DMR.Enrichment
DEGaP.Enrichment
C
HIPPOC Correlation: 0.067791919559029
PRDM9
Sox7
Hoxd13
Nkx2.2
ETS1
CDX4
NFAT:AP1
Rbpj1
Mef2d
X−box
ZNF189
RFX
Sox17
Srebp2
Sox2
Mef2c
Rfx2
Tbox:Smad
BORIS
Mef2a
Rfx1
AR−halfsite
Mef2b
MafB
Atf3
Npas4
Sox9
AP−1
Tbr1
Egr1
BATF
Bach2
Fosl2
DMR.Enrichment
DEGaP.Enrichment
D
KIDNEY Correlation: 0.027868072789259
PAX5
GLI3
Brn2
Srebp1a
CRE
HINFP
Ptf1a
PAX3:FKHR−fusion
Hoxb4
Tcf12
Npas4
MyoG
EWS:ERG−fusion
MYB
HEB
HNF6
Hnf6b
Ascl1
Atf7
Pit1
E2A
Etv2
Atoh1
PPARa
ERG
JunB
BATF
Fos
PPARE
Nur77
Atf3
AP−1
EAR2
RXR
Esrrb
Fra1
COUP−TFII
BORIS
HNF4a
EBF1
Hnf1
Fosl2
HNF1b
DMR.Enrichment
DEGaP.Enrichment
E
LIVER Correlation: 0.0389951846880909
Hoxa10
GLI3
Hoxd10
Hoxd11
Hoxd12
Etv2
MYB
Srebp2
Fos
Fra1
Nanog
BATF
Atf3
HINFP
Fosl2
Fra2
PPARa
HNF4a
JunB
AR−halfsite
ELF1
BORIS
ETS1
Nur77
PPARE
RXR
Foxf1
FOXK2
Fox:Ebox
FoxL2
Erra
HLF
KLF5
NFIL3
PBX1
Foxa3
Hnf1
FOXA1
FOXA1
HNF1b
DMR.Enrichment
DEGaP.Enrichment
F
LUNG Correlation: 0.114306884556539
Elk4
Npas4
ZBTB33
HIC1
Hoxd10
ETV4
ETV1
Nkx2.1
E2F3
Etv2
Fli1
ETS1
MYB
Nkx2.2
NF1−halfsite
NFY
ZNF519
Hoxa10
MafA
ZBTB12
Tcf3
ETS:RUNX
Atf2
ELF1
EHF
MafK
Foxo1
SPDEF
ERG
BMYB
IRF8
EWS:ERG−fusion
JunB
Fra1
Tlx?
Oct11
Egr1
NFIL3
HLF
Atf3
Fos
BATF
BORIS
Foxa2
AP−1
Sp5
EBF
EBF2
NF1
Elf4
ELF5
FOXA1
ELF3
SpiB
FOXM1
Fra2
FOXA1
DMR.Enrichment
DEGaP.Enrichment
G
BAT Correlation: 0.295975507234349
Fli1
Elk4
Elk1
ETV4
NF1−halfsite
TEAD3
ELF1
NF1
TEAD4
TEAD2
COUP−TFII
GABPA
Gli2
ZNF711
ETV1
EBF2
Gata2
ERG
RXR
E2F
RUNX
EWS:FLI1−fusion
EBF
ETS1
Erra
RUNX2
EAR2
Fos
COUP−TFII
JunB
Fra1
Elf4
Atf3
Bach2
ELF5
Etv2
CEBP
BORIS
Sp5
Esrrb
KLF5
FOXA1
FOXA1
PU.1
DMR.Enrichment
DEGaP.Enrichment
H
WAT−SC Correlation: 0.275271192037804
Elk4
NF1−halfsite
Elk1
GLIS3
Fli1
EWS:ERG−fusion
NF1
EWS:FLI1−fusion
FOXK2
ELF5
Brn2
ELF1
ETV4
Fos
ISRE
AP−2gamma
TEAD3
EHF
TEAD4
ETV1
Etv2
Atf3
MafA
BATF
THRb
JunB
Fra1
FoxD3
SPDEF
ERG
NFIL3
ELF3
Fra2
Foxo1
Tcfcp2l1
GABPA
Elf4
AP−1
ETS1
Fosl2
Jun−AP1
IRF3
HIC1
HLF
MafK
AR−halfsite
FoxL2
RXR
Foxa2
FOXK1
Bach2
CEBP
KLF6
Erra
p53
Sox3
PPARa
EBF2
Foxf1
HNF4a
Hnf1
EBF
Foxo3
PU.1
KLF5
SpiB
DMR.Enrichment
DEGaP.Enrichment

Supplemental Figure S29: Comparing TF enrichment among DMRs and DEGaPs. Scatter plots of -log10 p-value of TF motif enrichment among DARs and DEGaPs in six tissues: (a) skeletal muscle, (b) heart, (c) hippocampus, (d) kidney, (e) liver, (f) lung, (g) brown adipose, and (h) white adipose. Pearson correlation is calculated for each tissue.

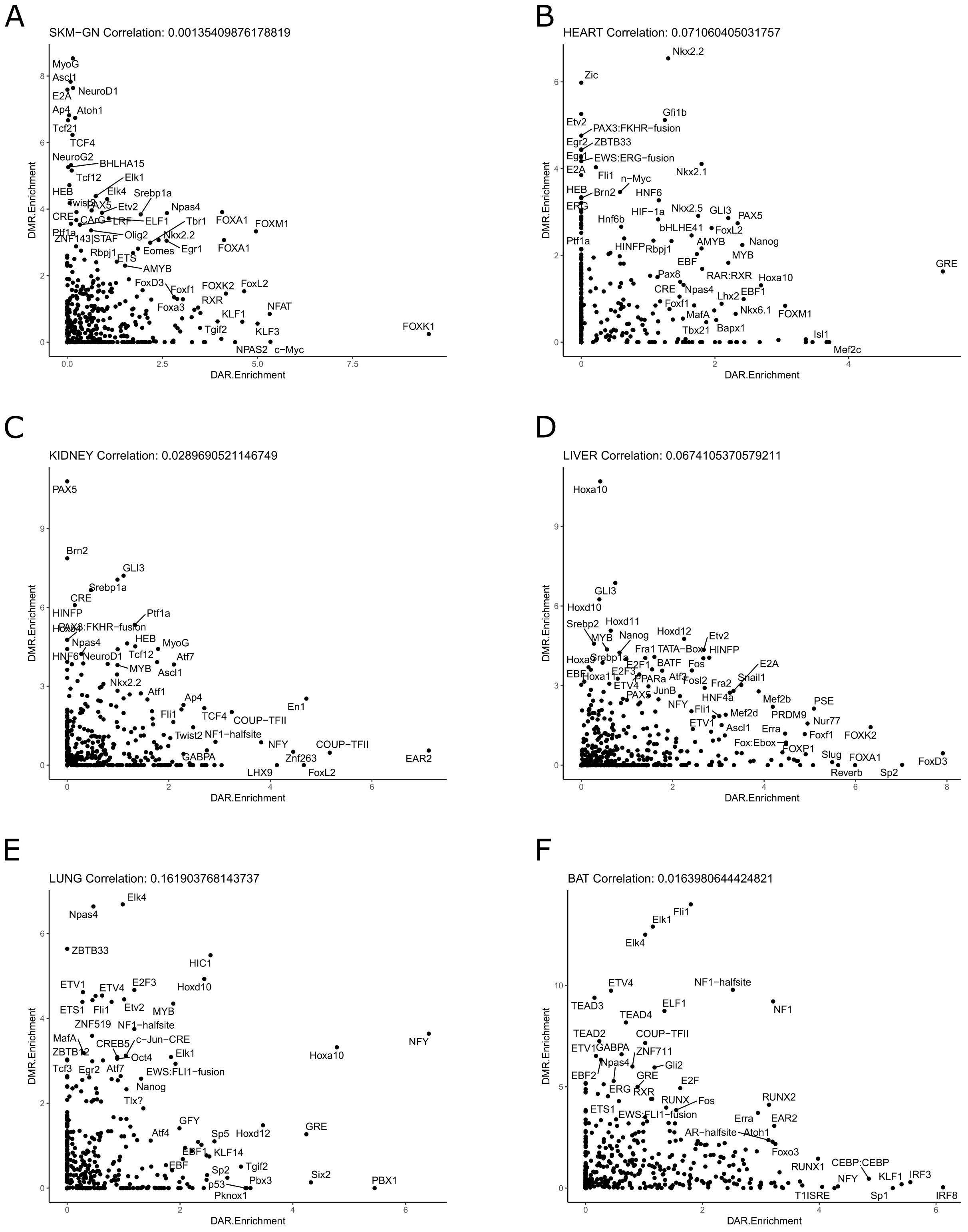
A
SKM−GN Correlation: 0.00135409876178819
DMR.Enrichment
DAR.Enrichment
MyoG
Ascl1
E2A
NeuroD1
Ap4
Atoh1
Tcf21
TCF4
NeuroG2
BHLHA15
Tcf12
Elk1
HEB
Elk4
Srebp1a
Etv2
Npas4
CRE
ELF1
Tbr1
FOXA1
FOXM1
Ptf1a
Olig2
Nkx2.2
Eomes
Egr1
Rbpj1
ETS
AMYB
FoxD3
Foxf1
FOXK2
FoxL2
Foxa3
RXR
NFAT
KLF1
Tgif2
KLF3
NPAS2
c−Myc
FOXK1
B
HEART Correlation: 0.071060405031757
Nkx2.2
Zic
Gfi1b
Etv2
PAX3:FKHR−fusion
ZBTB33
EWS:ERG−fusion
E2A
Fli1
n−Myc
Nkx2.1
HEB
Brn2
HNF6
ERG
HIF−1a
Nkx2.5
GLI3
PAX5
Hnf6b
bHLHE41
FoxL2
Ptf1a
AMYB
Nanog
HINFP
Rbpj1
MYB
EBF
GRE
Pax8
RAR:RXR
Hoxa10
CRE
Npas4
EBF1
Foxf1
Lhx2
Nkx6.1
FOXM1
MafA
Tbx21
Bapx1
Isl1
Mef2c
C
KIDNEY Correlation: 0.0289690521146749
PAX5
Brn2
GLI3
Srebp1a
CRE
HINFP
Ptf1a
PAX3:FKHR−fusion
HEB
Npas4
MyoG
HNF6
NeuroD1
Tcf12
Atf7
MYB
Ascl1
Nkx2.2
Atf1
Ap4
En1
Fli1
TCF4
COUP−TFII
Twist2
NF1−halfsite
NFY
COUP−TFII
GABPA
Znf263
EAR2
LHX9
FoxL2
D
LIVER Correlation: 0.0674105370579211
Hoxa10
GLI3
Hoxd10
Srebp2
Hoxd11
Hoxd12
MYB
Nanog
Etv2
Fra1
TATA−Box
HINFP
Srebp1a
E2F1
BATF
Fos
E2A
Hoxa11
E2F3
PPARa
Atf3
Fosl2
Fra2
Snail1
ETV4
JunB
PAX5
HNF4a
Mef2b
NFY
PSE
Fli1
Mef2d
PRDM9
ETV1
Ascl1
Nur77
Erra
Foxf1
FOXK2
Fox:Ebox
FOXP1
Slug
FOXA1
Reverb
Sp2
FoxD3
E
LUNG Correlation: 0.161903768143737
Elk4
Npas4
ZBTB33
HIC1
ETV1
ETV4
E2F3
Hoxd10
ETS1
Fli1
Etv2
MYB
ZNF519
NF1−halfsite
c−Jun−CRE
NFY
MafA
CREB5
Hoxa10
Oct4
Elk1
Tcf3
Egr2
Atf7
EWS:FLI1−fusion
Nanog
Tlx?
GFY
Atf4
Sp5
Hoxd12
GRE
EBF1
KLF14
EBF
Tgif2
Sp2
Six2
p53
Pbx3
PBX1
Pknox1
F
BAT Correlation: 0.0163980644424821
Fli1
Elk1
Elk4
ETV4
NF1−halfsite
TEAD3
ELF1
NF1
TEAD4
TEAD2
COUP−TFII
ETV1
GABPA
ZNF711
Npas4
Gli2
EBF2
GRE
E2F
ERG
RXR
RUNX2
RUNX
Fos
ETS1
EWS:FLI1−fusion
Erra
EAR2
AR−halfsite
Atoh1
Foxo3
RUNX1
CEBP:CEBP
IRF3
NFY
KLF1
T1ISRE
Sp1
IRF8

Supplemental Figure S30: Comparing TF enrichment among DARs and DMRs. Scatter plots of -log10 p-value of TF motif enrichment among DARs and DEGaPs in six tissues: (a) skeletal muscle, (b) heart, (c) kidney, (d) liver, (e) lung, (f) brown adipose. Pearson correlation is calculated for each tissue.

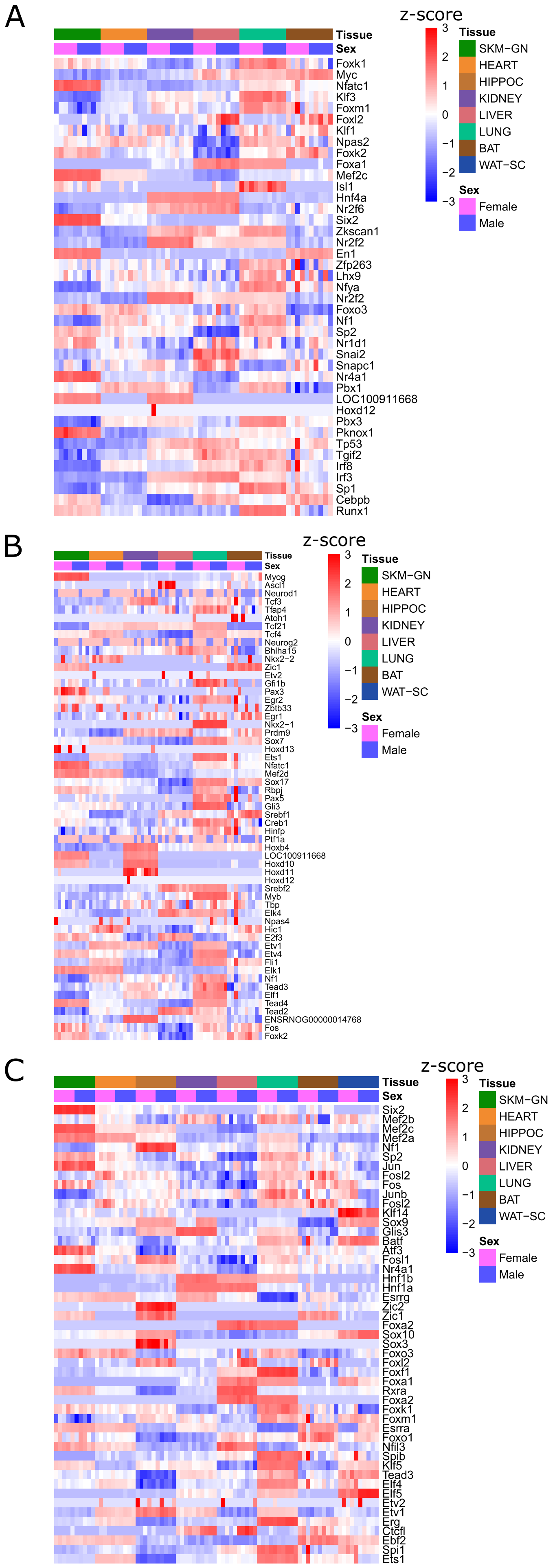

A
z-score
3
2
1
0
−1
−2
−3
Tissue
Sex
Tissue
SKM−GN
HEART
HIPPOC
KIDNEY
LIVER
LUNG
BAT
WAT−SC
Sex
Female
Male
Foxk1
Myc
Nfatc1
Klf3
Foxm1
Foxl2
Klf1
Npas2
Foxk2
Foxa1
Mef2c
Isl1
Hnf4a
Nr2f6
Six2
Zkscan1
Nr2f2
En1
Zfp263
Lhx9
Nfya
Nr2f2
Foxo3
Nf1
Sp2
Nr1d1
Snai2
Snapc1
Nr4a1
Pbx1
LOC100911668
Hoxd12
Pbx3
Pknox1
Tp53
Tgif2
Irf8
Irf3
Sp1
Cebpb
Runx1
B
z-score
Myog
Ascl1
Neurod1
Tcf3
Tfap4
Atoh1
Tcf21
Tcf4
Neurog2
Bhlha15
Nkx2−2
Zic1
Etv2
Gfi1b
Pax3
Egr2
Zbtb33
Egr1
Nkx2−1
Prdm9
Sox7
Hoxd13
Ets1
Nfatc1
Mef2d
Sox17
Rbpj
Pax5
Gli3
Srebf1
Creb1
Hinfp
Ptf1a
Hoxb4
LOC100911668
Hoxd10
Hoxd11
Hoxd12
Srebf2
Myb
Tbp
Elk4
Npas4
Hic1
E2f3
Etv1
Etv4
Fli1
Elk1
Nf1
Tead3
Elf1
Tead4
Tead2
ENSRNOG00000014768
Fos
Foxk2
C
z-score
Six2
Mef2b
Mef2c
Mef2a
Nf1
Sp2
Jun
Fosl2
Fos
Junb
Fosl2
Klf14
Sox9
Glis3
Batf
Atf3
Fosl1
Nr4a1
Hnf1b
Hnf1a
Esrrg
Zic2
Zic1
Foxa2
Sox10
Sox3
Foxo3
Foxl2
Foxf1
Foxa1
Rxra
Foxa2
Foxk1
Foxm1
Esrra
Foxo1
Nfil3
Spib
Klf5
Tead3
Elf4
Elf5
Etv2
Etv1
Erg
Ctcfl
Ebf2
Spi1
Ets1

Supplemental Figure S31: (a) z-scores of TF gene expression in control samples across tissues for top DAR-enriched TFs. (b) z-scores of TF gene expression in control samples across tissues for top DMR-enriched TFs. (c) z-scores of TF gene expression in control samples across tissues for top DEGaP-enriched TFs.

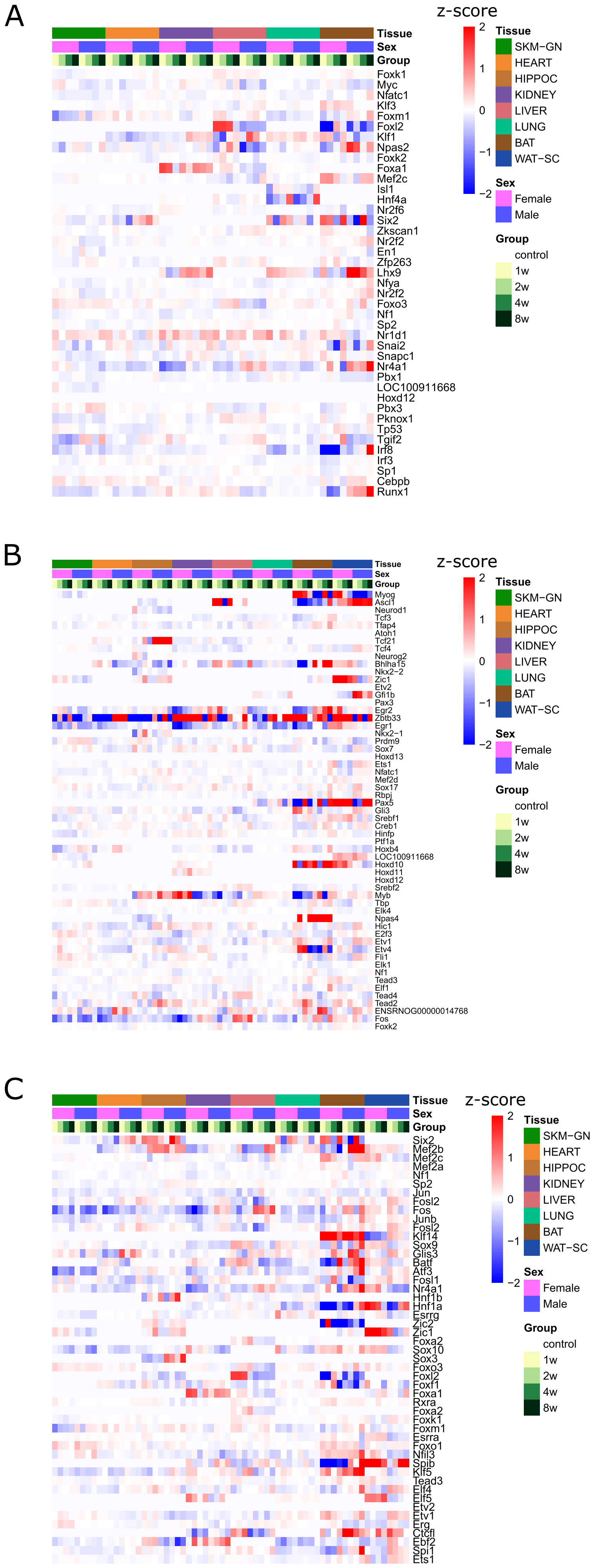
A
Tissue
Sex
Group
Foxk1
Myc
Nfatc1
Klf3
Foxm1
Foxl2
Klf1
Npas2
Foxk2
Foxa1
Mef2c
Isl1
Hnf4a
Nr2f6
Six2
Zkscan1
Nr2f2
En1
Zfp263
Lhx9
Nfya
Nr2f2
Foxo3
Nf1
Sp2
Nr1d1
Snai2
Snapc1
Nr4a1
Pbx1
LOC100911668
Hoxd12
Pbx3
Pknox1
Tp53
Tgif2
Irf8
Irf3
Sp1
Cebpb
Runx1
z-score
2
1
0
−1
−2
Tissue
SKM−GN
HEART
HIPPOC
KIDNEY
LIVER
LUNG
BAT
WAT−SC
Sex
Female
Male
Group
control
1w
2w
4w
8w
B
Tissue
Sex
Group
Myog
Ascl1
Neurod1
Tcf3
Tfap4
Atoh1
Tcf21
Tcf4
Neurog2
Bhlha15
Nkx2−2
Zic1
Etv2
Gfi1b
Pax3
Egr2
Zbtb33
Egr1
Nkx2−1
Prdm9
Sox7
Hoxd13
Ets1
Nfatc1
Mef2d
Sox17
Rbpj
Pax5
Gli3
Srebf1
Creb1
Hinfp
Ptf1a
Hoxb4
LOC100911668
Hoxd10
Hoxd11
Hoxd12
Srebf2
Myb
Tbp
Elk4
Npas4
Hic1
E2f3
Etv1
Etv4
Fli1
Elk1
Nf1
Tead3
Elf1
Tead4
Tead2
ENSRNOG00000014768
Fos
Foxk2
z-score
2
1
0
−1
−2
Tissue
SKM−GN
HEART
HIPPOC
KIDNEY
LIVER
LUNG
BAT
WAT−SC
Sex
Female
Male
Group
control
1w
2w
4w
8w
C
Tissue
Sex
Group
Six2
Mef2b
Mef2c
Mef2a
Nf1
Sp2
Jun
Fosl2
Fos
Junb
Fosl2
Klf14
Sox9
Glis3
Batf
Atf3
Fosl1
Nr4a1
Hnf1b
Hnf1a
Esrrg
Zic2
Zic1
Foxa2
Sox10
Sox3
Foxo3
Foxl2
Foxf1
Foxa1
Rxra
Foxa2
Foxk1
Foxm1
Esrra
Foxo1
Nfil3
Spib
Klf5
Tead3
Elf4
Elf5
Etv2
Etv1
Erg
Ctcfl
Ebf2
Spi1
Ets1
z-score
2
1
0
−1
−2
Tissue
SKM−GN
HEART
HIPPOC
KIDNEY
LIVER
LUNG
BAT
WAT−SC
Sex
Female
Male
Group
control
1w
2w
4w
8w

Supplemental Figure S32: (a) Training response L2FC across sexes, time points and tissues for top DAR-enriched TFs. (b) Training response L2FC across sexes, time points and tissues for top DMR-enriched TFs. (c) Training response L2FC across sexes, time points and tissues for top DEGaP-enriched TFs.

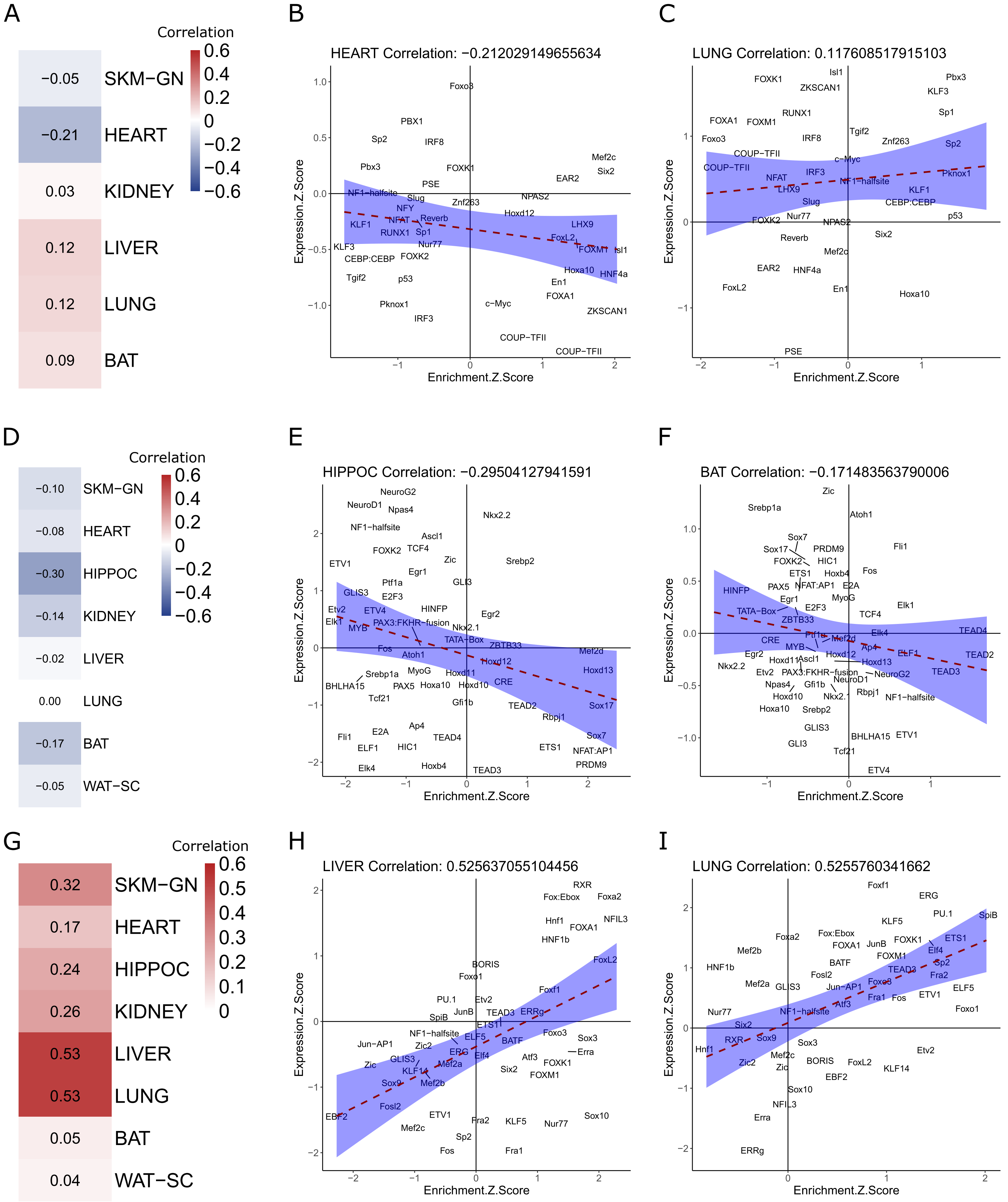
A
Correlation
−0.05 SKM−GN
−0.21 HEART
0.03 KIDNEY
0.12 LIVER
0.12 LUNG
0.09 BAT
B
HEART Correlation: −0.212029149655634
C
LUNG Correlation: 0.117608517915103
D
Correlation
−0.10 SKM−GN
−0.08 HEART
−0.30 HIPPOC
−0.14 KIDNEY
−0.02 LIVER
0.00 LUNG
−0.17 BAT
−0.05 WAT−SC
E
HIPPOC Correlation: −0.29504127941591
F
BAT Correlation: −0.171483563790006
G
Correlation
0.32 SKM−GN
0.17 HEART
0.24 HIPPOC
0.26 KIDNEY
0.53 LIVER
0.53 LUNG
0.05 BAT
0.04 WAT−SC
H
LIVER Correlation: 0.525637055104456
I
LUNG Correlation: 0.5255760341662
Expression.Z.Score
Enrichment.Z.Score

Supplemental Figure S33: (a) Correlation, for each tissue, of z-scores of TF motif frequency among DARs across tissues (S27a) and z-scores of TF gene expression in control samples across tissues (S28a). (b,c) Scatter plots of enrichment z score vs expression z score for top DAR-enriched TFs in heart (b) and lung (c). (d) Correlation, for each tissue, of z-scores of TF motif frequency among DMRs across tissues (S27b) and z-scores of TF gene expression in control samples across tissues (S28b). (e,f) Scatter plots of enrichment z score vs expression z score for top DAR-enriched TFs in hippocampus (e) and brown adipose (f). (g) Correlation, for each tissue, of z-scores of TF motif frequency among DEGaPs across tissues (S27c) and z-scores of TF gene expression in control samples across tissues (S28c). (h,i) Scatter plots of enrichment z score vs expression z score for top DEGaP-enriched TFs in liver (h) and lung (i). Pearson correlation is calculated for each tissue and a linear model is fit to the data.

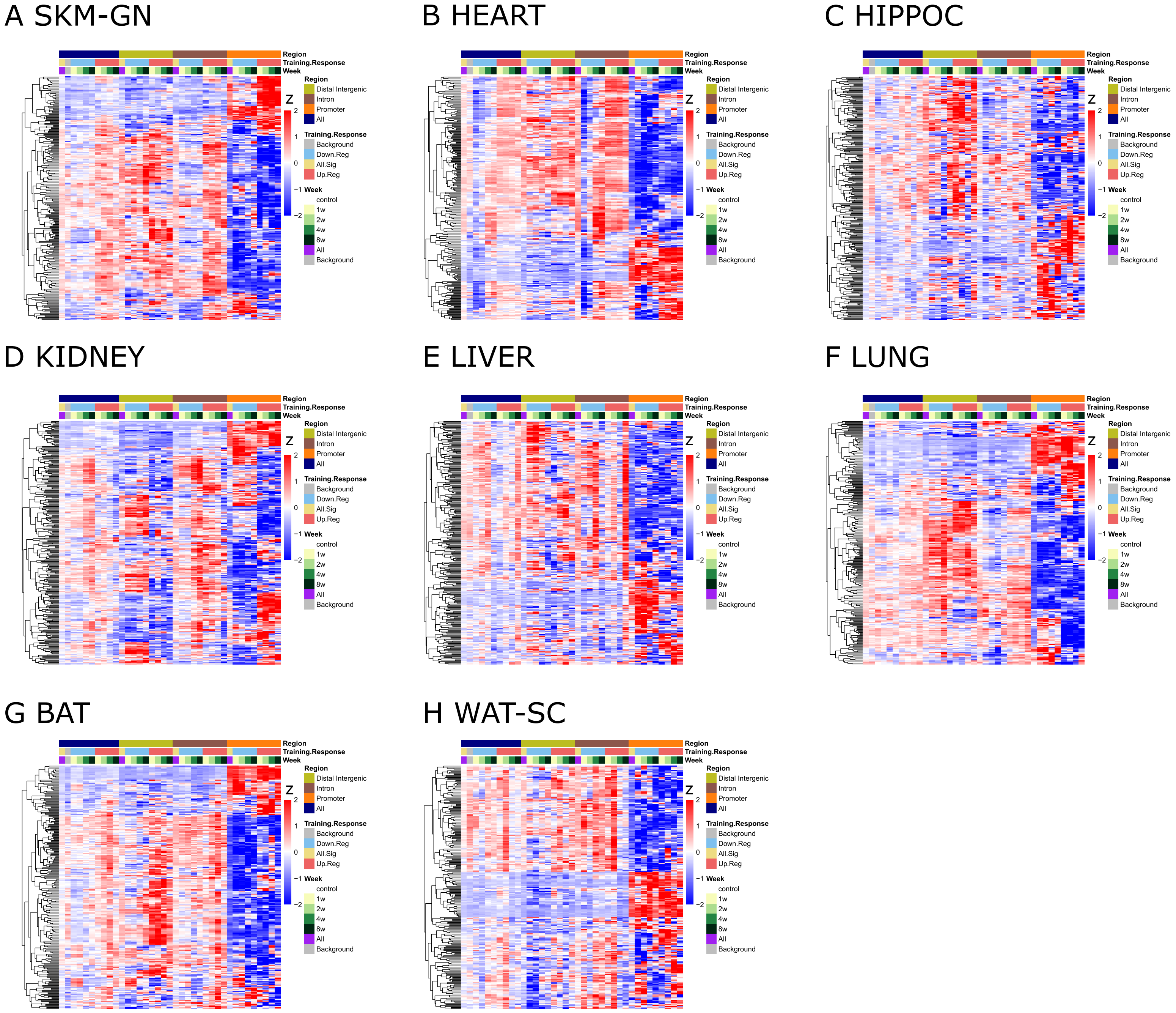
A SKM-GN
B HEART
C HIPPOC
D KIDNEY
E LIVER
F LUNG
G BAT
H WAT-SC
Region
Training.Response
Week
Z
2
1
0
−1
−2
Region
Distal Intergenic
Intron
Promoter
All
Training.Response
Background
Down.Reg
All.Sig
Up.Reg
Week
control
1w
2w
4w
8w
All
Background

Supplemental Figure S34: z-scores of TF motif frequency among DEGaPs divided into up-regulated and down-regulated genes at each training time point and split by genomic feature in: (a) skeletal muscle, (b) heart, (c) hippocampus, (d) kidney, (e) liver, (f) lung, (g) brown adipose, (h) white adipose. For comparison, each panel includes DEGaP sets not divided by genomic feature or training response. Distinct patterns of TF motif frequency among the up-regulated and down-regulated DEG promoter peaks.

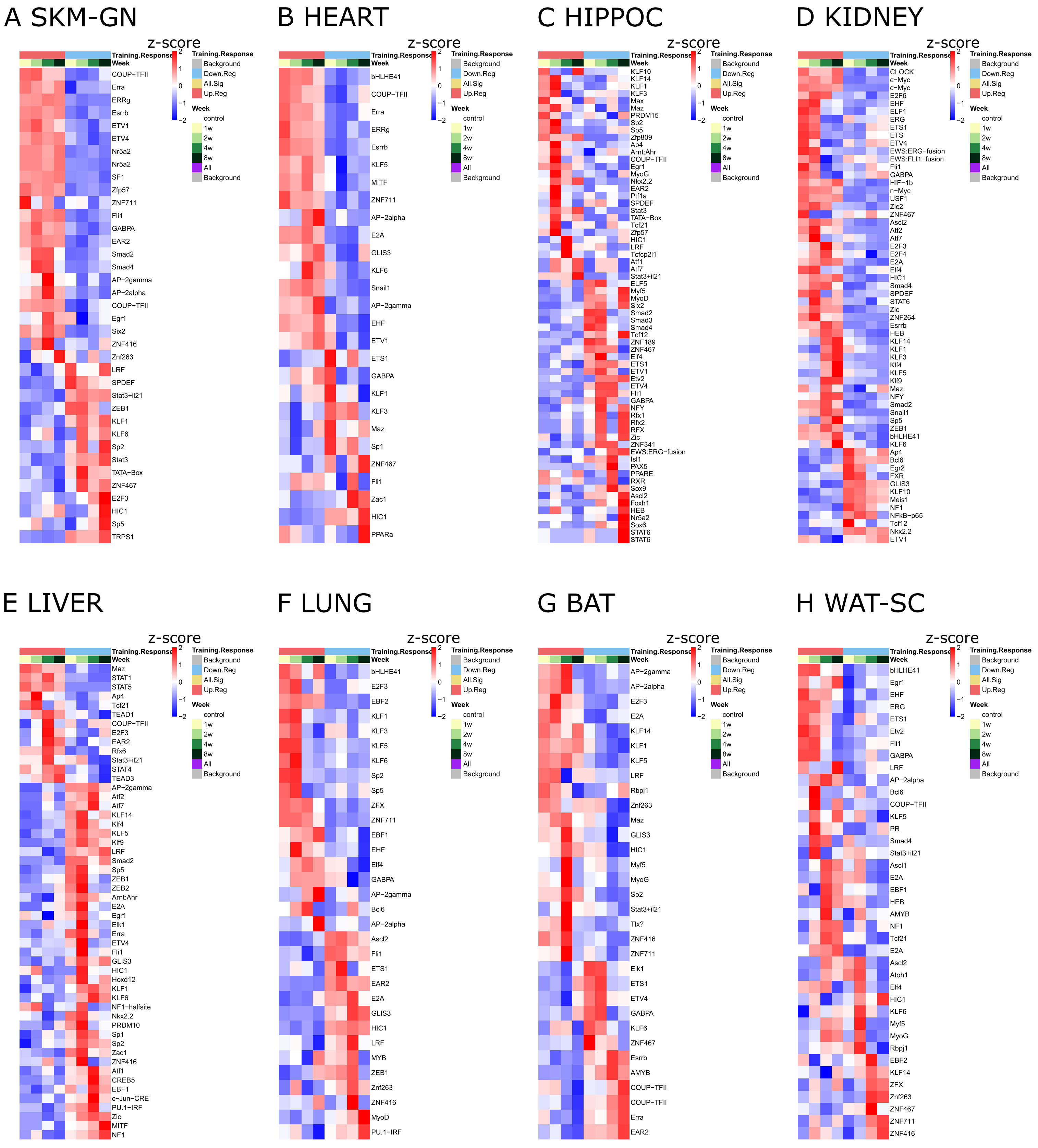
A SKM-GN
z-score
2
1
0
−1
−2
Training.Response
Week
Training.Response
Background
Down.Reg
All.Sig
Up.Reg
Week
control
1w
2w
4w
8w
All
Background
COUP−TFII
Erra
ERRg
Esrrb
ETV1
ETV4
Nr5a2
Nr5a2
SF1
Zfp57
ZNF711
Fli1
GABPA
EAR2
Smad2
Smad4
AP−2gamma
AP−2alpha
COUP−TFII
Egr1
Six2
ZNF416
Znf263
LRF
SPDEF
Stat3+il21
ZEB1
KLF1
KLF6
Sp2
Stat3
TATA−Box
ZNF467
E2F3
HIC1
Sp5
TRPS1
B HEART
z-score
2
1
0
−1
−2
Training.Response
Week
Training.Response
Background
Down.Reg
All.Sig
Up.Reg
Week
control
1w
2w
4w
8w
All
Background
bHLHE41
COUP−TFII
Erra
ERRg
Esrrb
KLF5
MITF
ZNF711
AP−2alpha
E2A
GLIS3
KLF6
Snail1
AP−2gamma
EHF
ETV1
ETS1
GABPA
KLF1
KLF3
Maz
Sp1
ZNF467
Fli1
Zac1
HIC1
PPARa
C HIPPOC
z-score
2
1
0
−1
−2
Training.Response
Week
Training.Response
Background
Down.Reg
All.Sig
Up.Reg
Week
control
1w
2w
4w
8w
All
Background
KLF10
KLF14
KLF1
KLF3
Max
Maz
PRDM15
Sp2
Sp5
Zfp809
Ap4
Arnt:Ahr
COUP−TFII
Egr1
MyoG
Nkx2.2
EAR2
Ptf1a
SPDEF
Stat3
TATA−Box
Tcf21
Zfp57
HIC1
LRF
Tcfcp2l1
Atf1
Atf7
Stat3+il21
ELF5
Myf5
MyoD
Six2
Smad2
Smad3
Smad4
Tcf12
ZNF189
ZNF467
Elf4
ETS1
ETV1
Etv2
ETV4
Fli1
GABPA
NFY
Rfx1
Rfx2
RFX
Zic
ZNF341
EWS:ERG−fusion
Isl1
PAX5
PPARE
RXR
Sox9
Ascl2
Foxh1
HEB
Nr5a2
Sox6
STAT6
STAT6
D KIDNEY
z-score
2
1
0
−1
−2
Training.Response
Week
Training.Response
Background
Down.Reg
All.Sig
Up.Reg
Week
control
1w
2w
4w
8w
All
Background
CLOCK
c−Myc
c−Myc
E2F6
EHF
ELF1
ERG
ETS1
ETS
ETV4
EWS:ERG−fusion
EWS:FLI1−fusion
Fli1
GABPA
HIF−1b
n−Myc
USF1
Zic2
ZNF467
Ascl2
Atf2
Atf7
E2F3
E2F4
E2A
Elf4
HIC1
Smad4
SPDEF
STAT6
Zic
ZNF264
Esrrb
HEB
KLF14
KLF1
KLF3
Klf4
KLF5
Klf9
Maz
NFY
Smad2
Snail1
Sp5
ZEB1
bHLHE41
KLF6
Ap4
Bcl6
Egr2
FXR
GLIS3
KLF10
Meis1
NF1
NFkB−p65
Tcf12
Nkx2.2
ETV1
E LIVER
z-score
2
1
0
−1
−2
Training.Response
Week
Training.Response
Background
Down.Reg
All.Sig
Up.Reg
Week
control
1w
2w
4w
8w
All
Background
Maz
STAT1
STAT5
Ap4
Tcf21
TEAD1
COUP−TFII
E2F3
EAR2
Rfx6
Stat3+il21
STAT4
TEAD3
AP−2gamma
Atf2
Atf7
KLF14
Klf4
KLF5
Klf9
LRF
Smad2
Sp5
ZEB1
ZEB2
Arnt:Ahr
E2A
Egr1
Elk1
Erra
ETV4
Fli1
GLIS3
HIC1
Hoxd12
KLF1
KLF6
NF1−halfsite
Nkx2.2
PRDM10
Sp1
Sp2
Zac1
ZNF416
Atf1
CREB5
EBF1
c−Jun−CRE
PU.1−IRF
Zic
MITF
NF1
F LUNG
z-score
2
1
0
−1
−2
Training.Response
Week
Training.Response
Background
Down.Reg
All.Sig
Up.Reg
Week
control
1w
2w
4w
8w
All
Background
bHLHE41
E2F3
EBF2
KLF1
KLF3
KLF5
KLF6
Sp2
Sp5
ZFX
ZNF711
EBF1
EHF
Elf4
GABPA
AP−2gamma
Bcl6
AP−2alpha
Ascl2
Fli1
ETS1
EAR2
E2A
GLIS3
HIC1
LRF
MYB
ZEB1
Znf263
ZNF416
MyoD
PU.1−IRF
G BAT
z-score
2
1
0
−1
−2
Training.Response
Week
Training.Response
Background
Down.Reg
All.Sig
Up.Reg
Week
control
1w
2w
4w
8w
All
Background
AP−2gamma
AP−2alpha
E2F3
E2A
KLF14
KLF1
KLF5
LRF
Rbpj1
Znf263
Maz
GLIS3
HIC1
Myf5
MyoG
Sp2
Stat3+il21
Tlx?
ZNF416
ZNF711
Elk1
ETS1
ETV4
GABPA
KLF6
ZNF467
Esrrb
AMYB
COUP−TFII
COUP−TFII
Erra
EAR2
H WAT-SC
z-score
2
1
0
−1
−2
Training.Response
Week
Training.Response
Background
Down.Reg
All.Sig
Up.Reg
Week
control
1w
2w
4w
8w
All
Background
bHLHE41
Egr1
EHF
ERG
ETS1
Etv2
Fli1
GABPA
LRF
AP−2alpha
Bcl6
COUP−TFII
KLF5
PR
Smad4
Stat3+il21
Ascl1
E2A
EBF1
HEB
AMYB
NF1
Tcf21
E2A
Ascl2
Atoh1
Elf4
HIC1
KLF6
Myf5
MyoG
Rbpj1
EBF2
KLF14
ZFX
Znf263
ZNF467
ZNF711
ZNF416

Supplemental Figure S35: Conserved TF motif frequency of up-regulated and down-regulated promoter peaks in each tissue. z-scores of TF motif frequency among DEG promoter peaks divided by up-regulated and down-regulated training responses at each training time point in: (a) skeletal muscle, (b) heart, (c) hippocampus, (d) kidney, (e) liver, (f) lung, (g) brown adipose, (h) white adipose. TFs were selected in each tissue for enrichment in up-regulated or down-regulated DEG promoter peaks and differentiation of enrichment between up-regulated and down-regulated DEG promoter peaks.

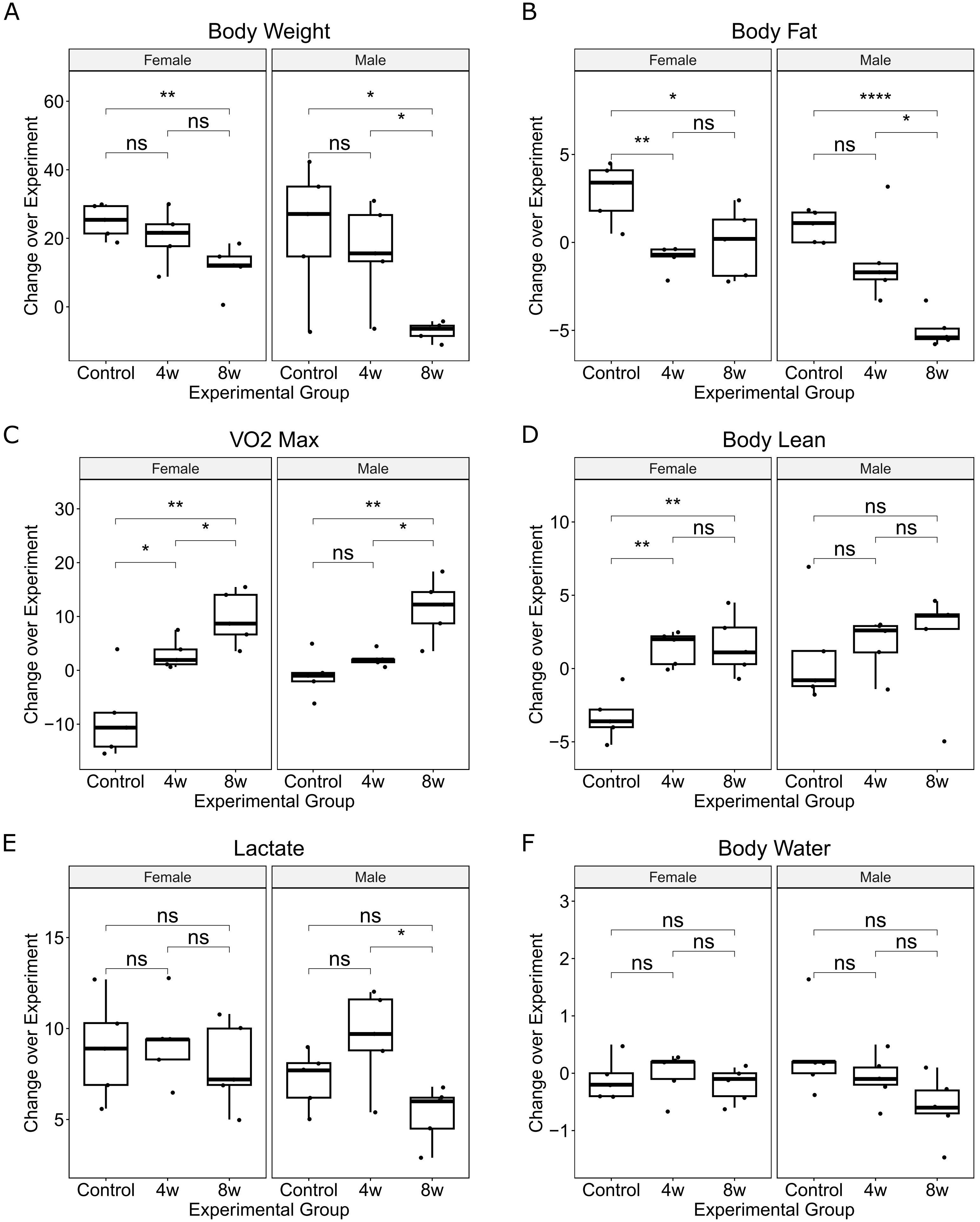
A
Body Weight
Female
Male
Change over Experiment
Experimental Group
Control
4w
8w
**
*
ns
B
Body Fat
****
C
VO2 Max
D
Body Lean
E
Lactate
F
Body Water

Supplemental Figure S36: Boxplots of phenotypic changes over training for body weight (a), body fat (b), vo2 max (c), body lean (d), lactate (e), and body water(f). Statistical measurements were generated with a t test. ns (not significant) represents $p >= 0.05$, *$p < 0.05$, **$p < 0.01$, ***$p < 0.001$, ****$p < 0.0001$.

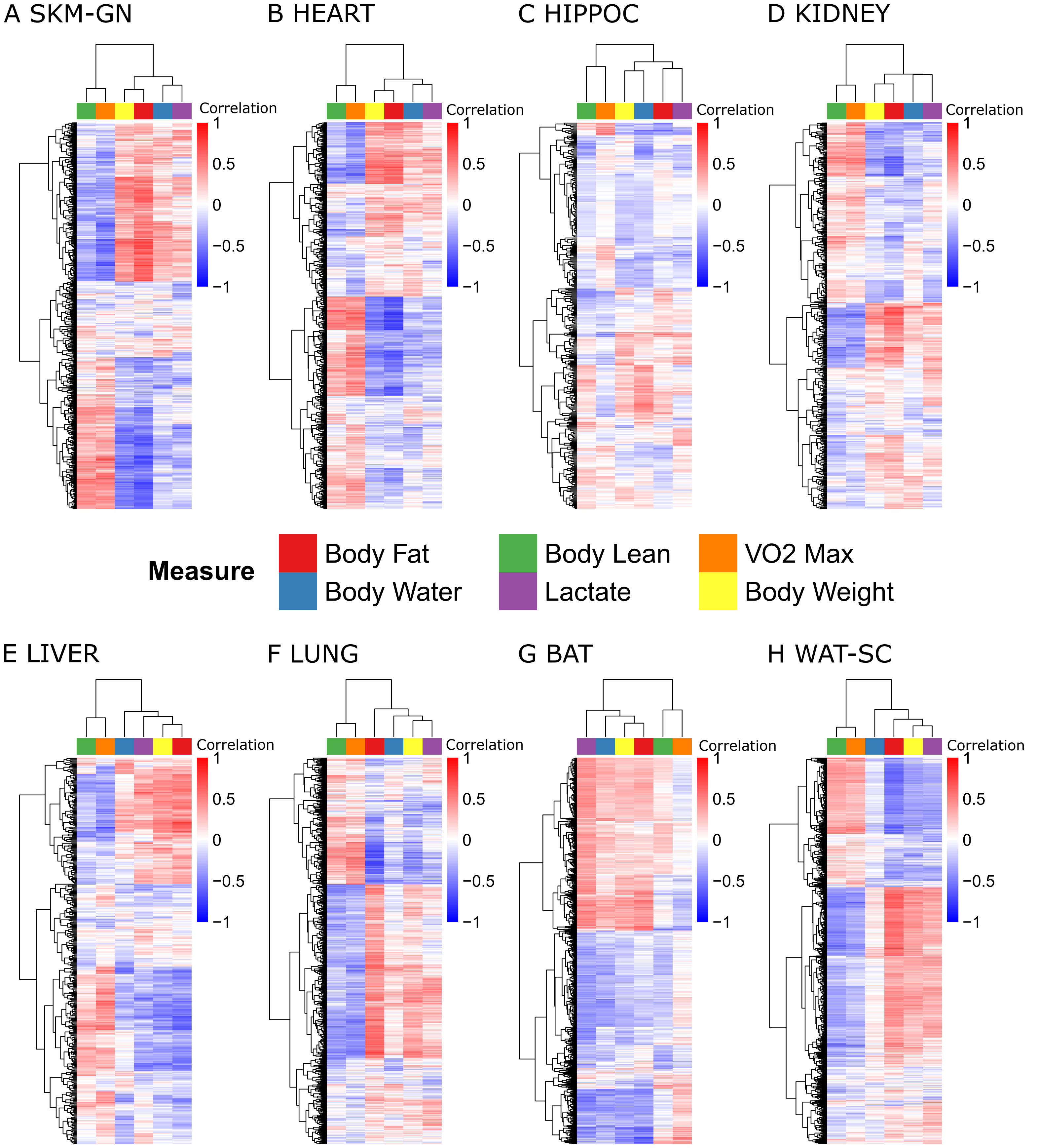
A SKM-GN
B HEART
C HIPPOC
D KIDNEY
Correlation
1
0.5
0
−0.5
−1
Measure
Body Fat
Body Water
Body Lean
Lactate
VO2 Max
Body Weight
E LIVER
F LUNG
G BAT
H WAT-SC

Supplemental Figure S37: Heatmaps of correlation between DEGs and phenotypic measures for eight tissues: skeletal muscle (a), heart (b), hippocampus (c), kidney (d), liver (e), lung (f), brown adipose (e), and white adipose (f). Most tissues show distinct groups of DEGs positively correlated with vo2 max and body lean vs body fat and weight gain, with the exception of brown adipose (DEGs driven by cell type changes).

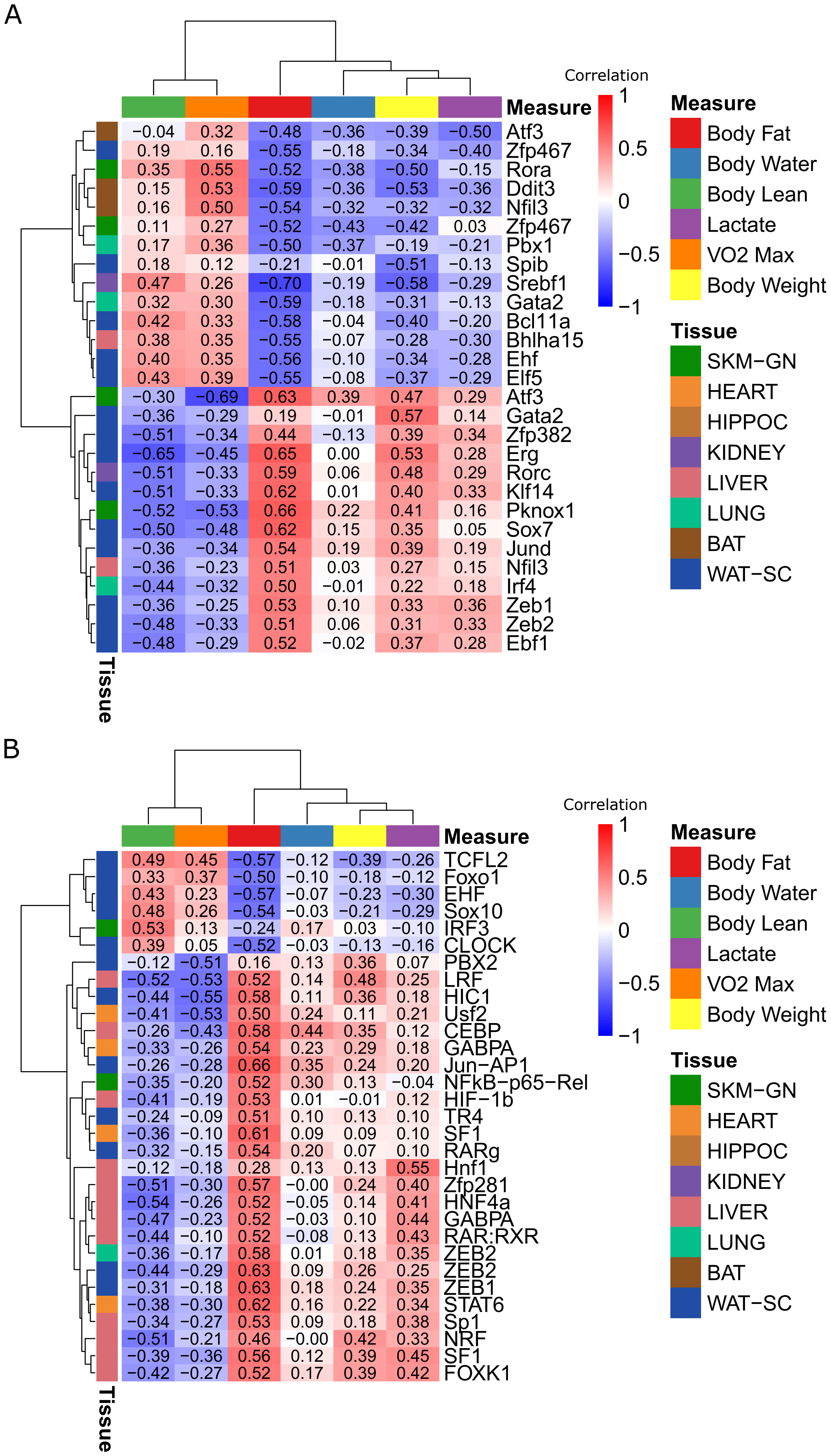
A
Correlation
1
0.5
0
−0.5
−1
Measure
Body Fat
Body Water
Body Lean
Lactate
VO2 Max
Body Weight
Tissue
SKM−GN
HEART
HIPPOC
KIDNEY
LIVER
LUNG
BAT
WAT−SC
−0.04 0.32 −0.48 −0.36 −0.39 −0.50 Atf3
0.19 0.16 −0.55 −0.18 −0.34 −0.40 Zfp467
0.35 0.55 −0.52 −0.38 −0.50 −0.15 Rora
0.15 0.53 −0.59 −0.36 −0.53 −0.36 Ddit3
0.16 0.50 −0.54 −0.32 −0.32 −0.32 Nfil3
0.11 0.27 −0.52 −0.43 −0.42 0.03 Zfp467
0.17 0.36 −0.50 −0.37 −0.19 −0.21 Pbx1
0.18 0.12 −0.21 −0.01 −0.51 −0.13 Spib
0.47 0.26 −0.70 −0.19 −0.58 −0.29 Srebf1
0.32 0.30 −0.59 −0.18 −0.31 −0.13 Gata2
0.42 0.33 −0.58 −0.04 −0.40 −0.20 Bcl11a
0.38 0.35 −0.55 −0.07 −0.28 −0.30 Bhlha15
0.40 0.35 −0.56 −0.10 −0.34 −0.28 Ehf
0.43 0.39 −0.55 −0.08 −0.37 −0.29 Elf5
−0.30 −0.69 0.63 0.39 0.47 0.29 Atf3
−0.36 −0.29 0.19 −0.01 0.57 0.14 Gata2
−0.51 −0.34 0.44 −0.13 0.39 0.34 Zfp382
−0.65 −0.45 0.65 0.00 0.53 0.28 Erg
−0.51 −0.33 0.59 0.06 0.48 0.29 Rorc
−0.51 −0.33 0.62 0.01 0.40 0.33 Klf14
−0.52 −0.53 0.66 0.22 0.41 0.16 Pknox1
−0.50 −0.48 0.62 0.15 0.35 0.05 Sox7
−0.36 −0.34 0.54 0.19 0.39 0.19 Jund
−0.36 −0.23 0.51 0.03 0.27 0.15 Nfil3
−0.44 −0.32 0.50 −0.01 0.22 0.18 Irf4
−0.36 −0.25 0.53 0.10 0.33 0.36 Zeb1
−0.48 −0.33 0.51 0.06 0.31 0.33 Zeb2
−0.48 −0.29 0.52 −0.02 0.37 0.28 Ebf1
Tissue
B
Correlation
1
0.5
0
−0.5
−1
Measure
Body Fat
Body Water
Body Lean
Lactate
VO2 Max
Body Weight
Tissue
SKM−GN
HEART
HIPPOC
KIDNEY
LIVER
LUNG
BAT
WAT−SC
0.49 0.45 −0.57 −0.12 −0.39 −0.26 TCFL2
0.33 0.37 −0.50 −0.10 −0.18 −0.12 Foxo1
0.43 0.23 −0.57 −0.07 −0.23 −0.30 EHF
0.48 0.26 −0.54 −0.03 −0.21 −0.29 Sox10
0.53 0.13 −0.24 0.17 0.03 −0.10 IRF3
0.39 0.05 −0.52 −0.03 −0.13 −0.16 CLOCK
−0.12 −0.51 0.16 0.13 0.36 0.07 PBX2
−0.52 −0.53 0.52 0.14 0.48 0.25 LRF
−0.44 −0.55 0.58 0.11 0.36 0.18 HIC1
−0.41 −0.53 0.50 0.24 0.11 0.21 Usf2
−0.26 −0.43 0.58 0.44 0.35 0.12 CEBP
−0.33 −0.26 0.54 0.23 0.29 0.18 GABPA
−0.26 −0.28 0.66 0.35 0.24 0.20 Jun−AP1
−0.35 −0.20 0.52 0.30 0.13 −0.04 NFkB−p65−Rel
−0.41 −0.19 0.53 0.01 −0.01 0.12 HIF−1b
−0.24 −0.09 0.51 0.10 0.13 0.10 TR4
−0.36 −0.10 0.61 0.09 0.09 0.10 SF1
−0.32 −0.15 0.54 0.20 0.07 0.10 RARg
−0.12 −0.18 0.28 0.13 0.13 0.55 Hnf1
−0.51 −0.30 0.57 −0.00 0.24 0.40 Zfp281
−0.54 −0.26 0.52 −0.05 0.14 0.41 HNF4a
−0.47 −0.23 0.52 −0.03 0.10 0.44 GABPA
−0.44 −0.10 0.52 −0.08 0.13 0.43 RAR:RXR
−0.36 −0.17 0.58 0.01 0.18 0.35 ZEB2
−0.44 −0.29 0.63 0.09 0.26 0.25 ZEB2
−0.31 −0.18 0.63 0.18 0.24 0.35 ZEB1
−0.38 −0.30 0.62 0.16 0.22 0.34 STAT6
−0.34 −0.27 0.53 0.09 0.18 0.38 Sp1
−0.51 −0.21 0.46 −0.00 0.42 0.33 NRF
−0.39 −0.36 0.56 0.12 0.39 0.45 SF1
−0.42 −0.27 0.52 0.17 0.39 0.42 FOXK1
Tissue

Supplemental Figure S38: Significant correlations of TF training response with phenotypic measures in RNA-seq (a) and proteomic (b) data. All TFs listed have at least one phenotypic correlation > 0.5 or < -0.5 in the annotated tissue (annotated at the left of each row).
